# Repeated disuse atrophy imprints a molecular memory in skeletal muscle: transcriptional resilience in young adults and susceptibility in aged muscle

**DOI:** 10.1101/2025.10.16.681134

**Authors:** Daniel C. Turner, Truls Raastad, Max Ullrich, Stian F. Christiansen, Hazel Sutherland, James Boot, Eva Wozniak, Charles Mein, Emilie Dalbram, Jonas T. Treebak, Daniel J. Owens, David C. Hughes, Sue C. Bodine, Jonathan C. Jarvis, Adam P. Sharples

**Affiliations:** Department for Physical Performance, Norwegian School of Sport Sciences, Oslo, Norway; Department of Molecular Medicine, University of Pavia, Italy; Research Institute for Sport and Exercise Sciences, Liverpool John Moores University (LJMU), Liverpool, United Kingdom; Genome Centre, Queen Mary University of London (QMUL), London, United Kingdom; Francis Crick Institute, London, United Kingdom; Novo Nordisk Foundation Center for Basic Metabolic Research, Faculty of Health and Medical Sciences, University of Copenhagen (UCPH), Copenhagen, Denmark; Aging and Metabolism Research Program, Oklahoma Medical Research Foundation (OMRF), Oklahoma City, Oklahoma, United States

**Keywords:** muscle memory, DNA methylation, aerobic metabolism, NAD^+^ metabolism, NR4A1, NR4A3 AChR (CHRNA1, *CHRND*, *CHRNG*), *NMRK2*, nicotinamide riboside, muscle stem cells

## Abstract

Disuse-induced muscle atrophy is common after illness, injury, or falls and becomes increasingly frequent with ageing. Whether skeletal muscle retains a “memory” of repeated disuse remains unknown. We investigated repeated lower-limb immobilization in young adults and a refined aged rat model, integrating physiological, multi-omic, immunohistochemical, biochemical, and primary human muscle stem cell (MuSC) analyses. To enable robust age comparisons, we integrated published young rat data with newly generated aged rat data. In young human muscle, repeated disuse elicited attenuated transcriptional perturbations in oxidative and mitochondrial pathways, suggestive of a protective molecular memory, despite similar atrophy to initial disuse. In contrast, aged muscle exhibited a detrimental memory, characterised by greater atrophy, exaggerated suppression of aerobic metabolism genes despite recovery after initial disuse, NAD⁺ and mitochondrial DNA depletion, and activation of proteasomal, extracellular-matrix, and DNA-damage pathways. Whereas young rats recovered muscle mass after initial disuse, aged rats did not. Repeated disuse induced DNA hypermethylation and downregulation of aerobic metabolism and mitochondrial genes across species. *NR4A1* remained suppressed into recovery, while AChR subunit genes (*CHRNA1*, *CHRND*) were epigenetically primed. *NMRK2*, an NAD⁺ biosynthesis gene, was among the most downregulated after both atrophy periods, and nicotinamide riboside improved myotube size in MuSCs post-atrophy. Repeated disuse atrophy imprints a molecular memory in skeletal muscle shaping transcriptional resilience in young adults and exaggerated susceptibility in aged muscle.

## Introduction

Age-related skeletal muscle loss (termed sarcopenia) is one of the main contributors to falling injury ^1, 2, 3^, often resulting in prolonged periods of disuse. Consequently, repeated falls in the elderly become increasingly common, leading to recurrent loss of skeletal muscle and thus increased risk of frailty, morbidity, hospital admittance, reduced quality of life and ultimately earlier mortality ^2, 4, 5, 6, 7^. Restoration of skeletal muscle following periods of disuse is also impaired in aged muscle ^5, 8, 9, 10, 11, 12^, further increasing the risk of repeated falls ^2, 3^. Despite the well-characterised skeletal muscle wasting (atrophy) response to a single period of limb immobilisation ^8, 9, 10, 13, 14, 15, 16, 17, 18, 19, 20, 21^, bed rest ^22, 23,24, 25, 26^ and spaceflight ^27, 28, 29^, whether muscle atrophy is exacerbated upon repeated disuse in both adult and aged muscle is currently unknown.

Skeletal muscle “memory” refers to the differential tissue response to repeated exposure to the same (or similar) environmental stimuli ^30, 31^. While it is evident that skeletal muscle exhibits a ‘positive’ memory of anabolic stimuli, resistance ^32, 33, 34, 35, 36, 37, 38, 39^ and endurance ^34, 40^ training in both humans and rodents, it is unknown whether skeletal muscle possesses a ‘negativ’ memory of disuse-induced muscle atrophy. Specifically, whether an earlier encounter of disuse renders the muscle more susceptible to further wasting if reencountered in the future following injury reoccurrence or repeated falling in the elderly.

Epigenetics is considered a key mechanism underpinning skeletal muscle memory, whereby retention of epigenetic modifications in skeletal muscle are either; a) modified following exercise training and retained during detraining, leading to sustained elevations in gene transcription ^33, 34, 36, 37, 39, 40^ and protein abundance ^38^ and/or; b) enhanced upon later retraining, resulting in amplified changes in gene expression ^33, 34, 36, 37, 39, 40^. Among the various epigenetic modifications, DNA methylation is one of the most prominent emerging regulators of skeletal muscle memory. DNA methylation undergoes genome-wide alterations and modulates the transcriptional response to acute exercise and chronic training in divergent pathways according to the mode of exercise performed ^33, 36, 37, 39, 40, 41, 42, 43, 44, 45, 46, 47, 48, 49, 50, 51, 52, 53, 54, 55^. Moreover, DNA methylation is dynamic and can both precede and modulate the time course of transient gene expression in skeletal muscle in the minutes to hours following a single bout of exercise (reviewed in ^56^). However, some DNA methylation changes are long-lasting, particularly those induced by chronic training, thus providing a key mechanism mediating skeletal muscle memory of exercise ^31, 33, 34, 36, 37, 39, 40^.

Despite the epigenetic role in modulating gene transcription in response to positive stimuli, studies examining the DNA methylation response to negative stimuli such as muscle disuse atrophy remain scarce. Nevertheless, there is clear evidence of a dysregulated methylome in skeletal muscle of aged ^57, 58, 59, 60, 61^ and diseased populations (e.g., type-II diabetes ^42, 51, 62^ and cancer ^48^), in which greater aerobic fitness (higher V̇O_2max_) ^57^, higher physical activity, or exercise training can recalibrate the nuclear ^39, 48, 57, 58, 59^ and mitochondrial methylome ^63^ toward epigenetic signatures resembling that of youthful muscle. Following disuse atrophy, hypermethylation and reduced expression of targeted genes such as *PPARGC1A* after nine days of bed rest in human muscle ^64^, and *Nos1* after acute immobilisation in mice, have been reported ^65^. Our group have also demonstrated hypomethylation and increased expression of E3 ligases (*Murf1* and *MAFbx/Fbxo32*), the acetylochline receptor subunit (*Chrna1*), and the histone deacetylase (*Hdac4*) following tetrodotoxin (TTX)-included disuse atrophy in young adult rat muscle ^66^. To date, however, most studies have predominantly focused on targeted gene methylation with limited data available on genome-wide methylation. A recent study combined methylome and transcriptome analyses after 2 weeks of disuse atrophy in young adult humans (via lower limb casting) and reported predominance of global hypermethylation that was associated with a larger number of downregulated versus upregulated genes ^39^. However, leg casting-induced disuse was preceded by resistance training, and therefore it is unknown whether such alterations occur following disuse atrophy without prior resistance training that is more representative of the general population ^39^. A recent study analysing genome-wide DNA methylation in purified muscle stem cells (MuSCs) following injury in adult mice demonstrated sustained differential methylation long after recovery ^67^, whereas myoblasts retain DNA methylation changes after repeated inflammatory exposure *in vitro* ^68^. However, no studies have investigated the combined methylome and transcriptome following repeated atrophy to determine whether skeletal muscle imprints a ‘negativ’ memory of disuse-induced atrophy in both young adult and aged muscle.

We aimed to determine whether skeletal muscle exhibits a ‘negative memory’ of disuse atrophy. We hypothesised that prior disuse episodes would leave persistent molecular imprints that exacerbate subsequent atrophy, particularly in aged muscle. Because repeated atrophy and thus frailty in elderly humans raises ethical concerns, we combined physiological, multi-omic, immunohistochemical, biochemical, and primary muscle stem cell analyses in parallel young adult human and aged animal models of disuse atrophy, recovery and repeated atrophy. To reduce animal use and enable a direct age comparison, we also integrated our previously published young rat data ^66^ with newly generated aged rat data, confirming that young rats recover muscle mass after disuse, whereas aged rats do not. We demonstrate that skeletal muscle retains a molecular memory of disuse. In young adults, repeated immobilization elicited a resilient and ‘protective’ transcriptional memory, characterised by attenuated changes in aerobic metabolism and mitochondrial pathways after repeated disuse, while muscle mass had fully recovered. However, this did not mitigate muscle loss after repeated atrophy. In contrast, repeated disuse in aged muscle exhibited a heightened susceptibility, with further suppression of aerobic metabolism and mitochondrial pathways, depletion of NAD⁺ and mtDNA content, and greater muscle wasting despite recovery of gene expression after initial disuse. Proteasomal, extracellular matrix (ECM), and DNA-damage pathways were uniquely elevated in aged muscle after repeated disuse. *NR4A3*, a nuclear receptor critical for muscle metabolism, showed the greatest downregulation among all genes after disuse atrophy in humans. Epigenomic profiling revealed conserved DNA hypermethylation in aerobic and mitochondrial gene networks across species following atrophy and repeated atrophy, alongside locus-specific memory genes, including *NR4A1* and AChR (*CHRNA1*, *CHRND*) subunit genes. Finally, NMRK2, a key NAD⁺ biosynthesis gene, was among most downregulated after both periods of atrophy in young adult humans and nicotinamide riboside (NR) partially restored NMRK2 and improved MYOG expression and myotube size in primary human MuSCs post-atrophy.

## Methods

### Human ethics and participant information

Ethical approval was granted by the Regional Committees for Medical and Health Research Ethics (REK, Application ID 227924). The study was registered with the Norwegian Agency for Shared Services in Education and Research (Sikt) under reference number 627824. Participants were familiarised with the study intervention and the possible associated risks. Informed written consent and completion of an approved health and pre-biopsy medical questionnaire were obtained prior to participation. Ten young adults, males (*n* = 7) and females (*n* = 3) living in Oslo, Norway participated in the study. Participants were free from any known injury, illness and disease and had no previous history or predisposition to deep vein thrombosis (DVT). Participant baseline characteristics are presented in **Table 1**. All human data collection was conducted at the Norwegian School of Sport Sciences.

**Table 1.** Young adult human baseline characteristics. Values presented as mean ± standard deviation (SD).

| | Male ($n = 7$) | Female ($n = 3$) | All ($n = 10$) |
| --- | --- | --- | --- |
| Age (yrs) | 27.1 ± 4.1 | 23.7 ± 3 | 26.5 ± 4.1 |
| Height (cm) | 181.7 ± 2.7 | 167.9 ± 7.5 | 177.5 ± 7.9 |
| Weight (kg) | 93.4 ± 16 | 62 ± 12.6 | 84 ± 20.9 |
| Body mass index, BMI (kg/m<sup>2</sup>) | 28.4 ± 5.3 | 21.8 ± 2.6 | 26.4 ± 5.5 |
| Total Lean body mass, LBM (kg) | 64.8 ± 4.4 | 41.5 ± 7.2 | 57.8 ± 12.3 |
| Fat mass, FM (kg) | 24.8 ± 13.1 | 18.3 ± 5.7 | 22.3 ± 11.4 |
| Fat free mass, FFM (kg) | 68.2 ± 4.5 | 43.9 ± 7.3 | 60.9 ± 12.8 |
| Body fat, BF (%) | 25.4 ± 9.4 | 29.1 ± 4.5 | 26.5 ± 8.2 |
| Immobilised leg lean mass, LM (kg) | 11.4 ± 1.2 | 7.4 ± 1.5 | 10.2 ± 2.3 |
| Non-immobilised leg LM (kg) | 11.5 ± 1.3 | 7.1 ± 1.6 | 8.9 ± 2.5 |

### Assessment of physical activity levels and dietary intake in humans

The International Physical Activity Questionnaire Short Form (IPAQ-SF) was used to assess short-term physical activity levels prior to the study ^68^. Participants also completed an additional non-validated questionnaire including questions related to the specific type of sport (e.g., team, endurance, speed/power) or activity (e.g. cardiorespiratory fitness, strength, concurrent), duration of participation and competition level (i.e. recreational, national, international) to assess long-term physical activity levels.

### Human study design

In a repeated measures design, participants underwent 2 weeks (wks) (14 days) disuse atrophy (via unilateral limb immobilization) → ≈ 7 wks (49 ± 5 d) recovery → 2 wks repeated disuse atrophy (**Figure 1A**). During immobilisation, participants were fitted with a knee brace (DonJoy^®^ X-Act ROM, Enovis/DJO LLC, USA) fixed at 30° knee flexion to prevent any weight bearing of the non-dominant leg. Crutches were provided to enable ambulation without weight bearing on the braced immobilised leg. Following immobilisation, the knee brace and crutches were removed and participants returned to their normal ambulatory activity for ≈ 7 wks to allow recovery. Finally, the same non-dominant leg underwent a second period of 2 wks of immobilisation (repeated atrophy). Before immobilisation, participants were trained using the crutches and knee brace and were instructed to only remove the knee brace for showering and bedtime. Daily exercises (e.g., ankle rotation/dorsiflexion/plantarflexion) were encouraged to reduce the risk of DVT^69^. There were no reported cases throughout the present study. Before and after each immobilization, muscle biopsies and measurements of SkM size and strength were conducted in the morning (± 1 hr between timepoints) under fasted conditions following the same testing order (**Figure 1A**). Participants were also advised to refrain from strenuous exercise and alcohol consumption 24 hrs prior to muscle biopsy and testing procedures. Muscle biopsies and the testing protocol are detailed below. Upon completion of the study intervention, for ethical reasons, all participants were provided with the opportunity to undertake supervised resistance training programme (3 d/wk for 7 wks consisting of 2 lower body and 1 upper body sessions each wk) to restore any loss of SkM. Two to four weeks of strength training is sufficient for restoring SkM following a similar duration of atrophy in young adult humans ^8, 11, 70^.

**Figure 1.**
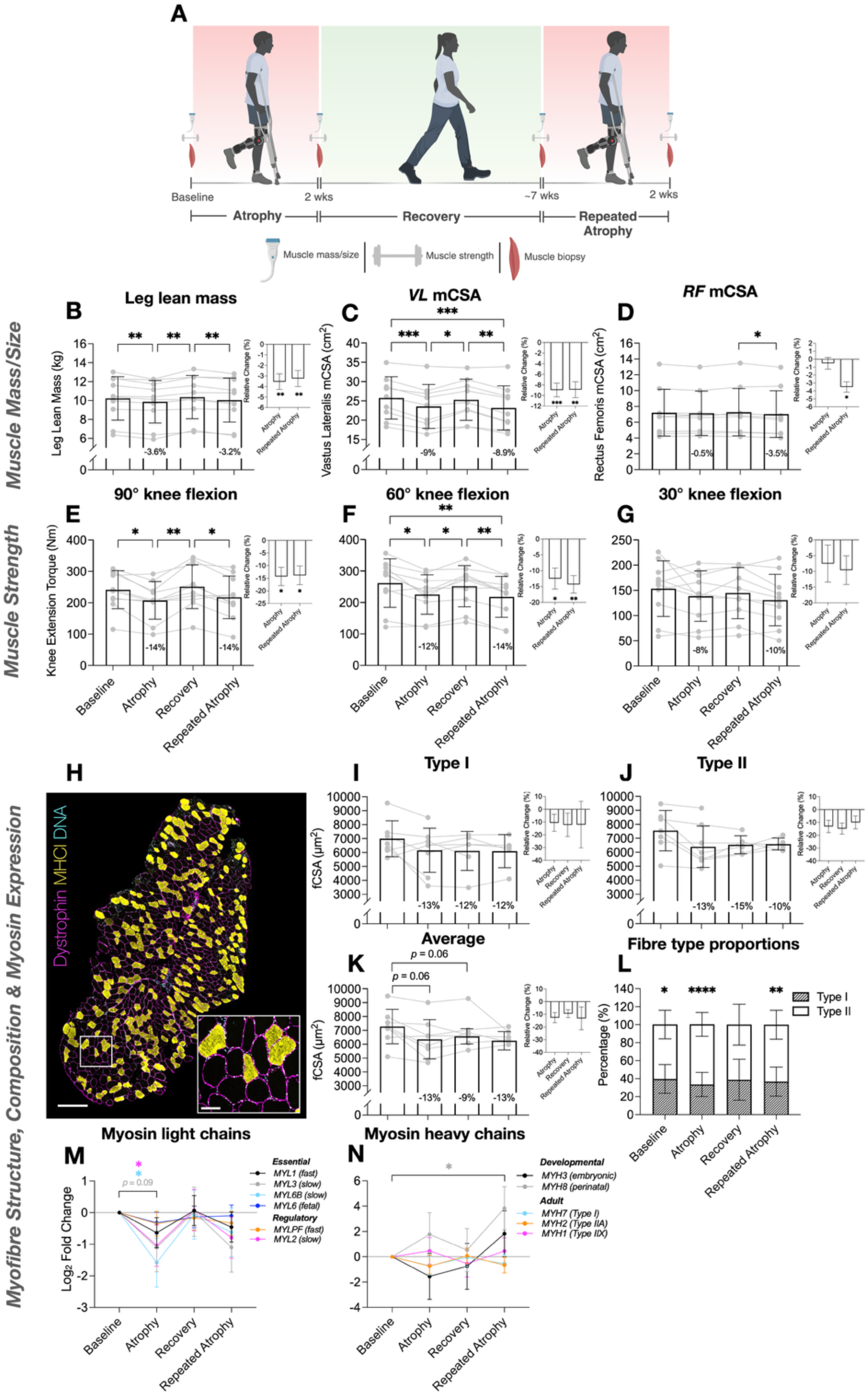
Young adult human intervention study design (created using BioRender.com) **(A).** Changes in leg lean mass **(B),** *Vastus Lateralis* (*VL*) mCSA **(C)**, *Rectus Femoris* (*RF)* mCSA **(D)** and strength/MVC across knee flexion angles 90, 60 and 30° **(E, F, G** respectively) after disuse (atrophy), recovery and repeated disuse (repeated atrophy). Significant reductions in muscle size and strength were observed after disuse atrophy, restored during recovery and reduced to a similar extent after repeated atrophy. *RF* mCSA only occurred after repeated disuse **(D).** Bar graphs depict relative change (%) from the previous timepoint (i.e., atrophy versus baseline, repeated atrophy versus recovery). *N* = 10 for leg lean mass **(A)** and strength **(E-J)**; *n* = 9 for mCSA (**B-D**). Myofibre analyses **(H-M)** revealed a non-significant reduction in fCSA **(H-K).** *N* = 8 baseline and atrophy, *n =* 6 recovery, *n* = 5 repeated atrophy. **(H)** Representative fluorescence microscopic image (10× magnification; magenta = dystrophin/myofibre membrane, yellow = MyHCI/type I myofibres, cyan = DNA, non-staining = putative type II myofibres; Scale bar within tile scan image = 500 µm, scale bar within magnified inset image = 100 µm). FibeRtypeR^82^ analysis suggested a larger proportion of fast/type II versus slow/type I fibres at baseline, atrophy and repeated atrophy with no significant differences at recovery (**L**). Despite no significant reduction in fCSA, there was reduced mRNA levels of slow isoform *MYL* and *MYH* genes after atrophy and repeated atrophy as well as increased fast type IIX (*MYH1*) and developmental *MYH* genes after repeated atrophy (**M & N**). *N* = 9 for each timepoint derived from RNA-seq data. \**p* ≤ 0.05, \*\**p* ≤ 0.01, \*\**p* ≤ 0.001, \*\*\*\**p* ≤ 0.0001.

### Human skeletal muscle biopsy procedures

Muscle biopsies were obtained from the immobilised leg according to our procedures described elsewhere ^48, 49^. Briefly, local anaesthesia (Xylocaine with 10 mg/mL lidocaine + 5 µg/mL adrenaline, AstraZeneca, Södertälje, Sweden) was administered to the biopsy area and muscle tissue (133 ± 47 mg excluding muscle tissue for immunohistochemical analysis, see below) was obtained from the mid-belly (10% proximal of the mid-thigh) of the *m. vastus lateralis (VL)* muscle using a modified Bergström technique (6 mm Pelomi-needle, Albertslund, Denmark) with manual suction (i.e., 50 mL syringe connected to the Bergström needle). All human muscle biopsies were obtained under fasted conditions at 09:30 ± 47 minutes across all timepoints and participants, following a standardised protocol to minimise circadian variability and ensure a narrow sampling window relative to diurnal regulation of skeletal muscle metabolism. For immunohistochemical (IHC) analysis, a proportion (≈ 50 mg) of muscle tissue was embedded in optimal cutting temperature (OCT) compound (CellPath, Chemi-Teknik AS, Oslo, Norway) ensuring myofibres were aligned in parallel before freezing in pre-cooled (i.e., above liquid nitrogen, LN_2_) isopentane (Sigma-Aldrich, Oslo, Norway) placed on dry ice and stored at −80 °C. Remaining tissue was rinsed in pre-cooled (4 °C) saline containing 0.9% NaCl (Braun, Melsungen, Germany) and dissected to remove visible fat, connective tissue and blood (vessels). Tissue intended for RNA, DNA and protein analysis was immediately weighed, transferred to nuclease-free Eppendorf tubes (Invitrogen™ RNase-free Microfuge Tubes, Fisher Scientific, Oslo, Norway), snap-frozen in LN_2_ and stored at −80 °C. Remaining tissue used for *in vitro* experiments was submerged in pre-cooled (4°C) transfer media (Hams F-10 medium that includes 1 mM L-glutamine/LG, 0.2% heat inactivated/hiFBS, 100 U/mL penicillin, 100 μg/mL streptomycin/PS and 2.5 µg/mL amphotericin-B/AB, Fisher Scientific) for subsequent primary cell isolations detailed below.

### Assessment of body composition in humans

Height (cm) and body weight (kg) were measured using a digital stadiometer and scale, respectively (Seca, Hamburg, Germany). Whole-body composition and leg lean mass (LLM) were assessed by dual-energy x-ray absorptiometry (DEXA; Lunar iDXA, enCORE software version 18; GE Healthcare, Chicago, USA). Total LLM was defined as all lean mass located distally from the collum femoris and inferior pubic ramus. A triangular segment depicting the pelvic area and a midline separating left and right legs were constructed as previously described ^71^. Specifically, the horizontal line of the triangle segment transversed slightly above the two iliac crests, and the two angled lines crossed the collum femoris, meeting just below the genital area ^71^. An additional line crossing the intercondylar eminence separated the upper and lower leg. All images were analysed single-blinded by three experienced personnel, and the average values were reported. The coefficient of variation (CV) for test-retest reliability was 1%.

### Assessment of muscle cross-sectional area (mCSA) in humans

Muscle cross-sectional area (mCSA) of the *m. rectus femoris (RF)* and *VL* muscles were assessed using panoramic B-mode ultrasound scanning using a linear transducer (L18-5) fixed to an ultrasound system (MACH 30; Hologic SuperSonic Imagine™, Aix-en-Provence, France). During scanning, participants lay supine (knee angle at 0 °C) with their leg muscles relaxed and feet strapped to a stable box to prevent any hip rotation. Three measurements per muscle (*RF* and *VL* each scanned separately) were measured and analysed by following a transverse trajectory line drawn across the mid-thigh (i.e., 50% of the distance between the greater trochanter and distal end of the muscle) using a marker pen, demonstrating a low <2.5% CV across all timepoints (RF baseline 2.4 ± 1.1%, atrophy 1.6 ± 1%, recovery 2.1 ± 1.1%, repeated atrophy 1.8 ± 1.6%, VL baseline 1.7 ± 1.3%, atrophy 1.3 ± 0.7%, 1.8 ± 1% recovery, 1.6 ± 0.7% repeated atrophy). Only acquired images free from visible artefacts/stitching were considered acceptable for analysis. Participants were encouraged to redraw the trajectory line during the immobilization and recovery periods to ensure accurate probe repositioning during subsequent timepoints. A transparent plastic sheet was also used to mark the trajectory and other clear landmarks (e.g., scars, patella and other marks) to provide additional accuracy during rescanning. Finally, previous images were displayed during rescanning to confirm image acquisition accuracy between timepoints. All images were analysed single-blinded using ImageJ2 with Bio-Format plugin (version 2.14.0/1.54f, U. S. National Institutes of Health, Bethesda, Maryland, USA) ^57,58^ at the end of the intervention. One participant was excluded from all mCSA analyses due to inaccurate repositioning and/or visible imaging artefacts.

### Assessment of muscle strength in humans

Voluntary maximal Isometric knee extension and flexion torque (Nm) was measured at 90, 60 and 30° knee flexion using a dynamometer (HUMAC NORM; Computer Sports Medicine inc, Massachusetts, USA). Participants were first familiarised with the protocol one week prior to baseline testing. Prior to strength testing, participants performed a standardised warm-up consisting of 5 mins low intensity cycling, 5 mins stretching and 3 × submaximal efforts (50-80%). Following the warm-up, participants performed 5 × maximal repetitions at each knee flexion angle (in order of 90, 60 and 30°). Verbal encouragement was provided during each repetition. The greatest torque produced was considered their maximum voluntary contraction (MVC). If the difference in torque between the first and second strongest repetition was >5%, participants performed an additional repetition. MVC was acquired on MP150 (Biopac, Goleta, CA, USA) with a sampling frequency of 1000 Hz and analysed using AcqKnowledge software (version 4.4.2, Biopac Systems, Inc., Knivsta, Sweden).

### Animals and study design

Repeated disuse in elderly humans is associated with frailty and increased risk of falls, raising ethical concerns. Therefore, we limited human experiments to young adults and used aged rats as a surrogate model for repeated atrophy in aged muscle. Further, to minimise animal use and adhere to ethical refinement principles, we did not include a direct young rat control group; instead, we integrated previously published young rat data ^66^ with newly generated aged rat data, enabling robust age comparisons while reducing unnecessary duplication in young adult rats. We confirmed that young rats recover muscle mass after disuse, whereas aged rats do not. Ethical approval was obtained, and experimental procedures were conducted with the permissions within a project license granted (PA693D221) under the British Home Office Animals (Scientific Procedures) Act 1986. Aged (23 ± 1 months) male Fischer 344 rats (Charles River) weighing 354 ± 54 g were housed at 20°C and 45% relative humidity on an alternating 12 hr light and 12 hr dark cycle, with food and water available *ad libitum*. Animals were assigned to 1 of 4 groups (*n* = 3 per group) including a control group (sham control) and 3 experimental groups (atrophy, recovery and repeated atrophy). All experimental groups received tetrodotoxin (TTX) to the common peroneal nerve (CPN), preventing contraction of the ankle dorsiflexor muscles. TTX binds to the voltage-gated sodium channel in the nerve cell membranes and so prevents the conduction of action potentials. Therefore, muscles innervated by the blocked CPN cannot be activated to contract. This induces disuse atrophy of the hind limb dorsiflexor muscles i.e., the tibialis anterior (TA) and extensor digitorum longus (EDL), and is advantageous vs. denervation or hindlimb unloading as it allows normal conduction in the tibial nerve to achieve plantarflexion, and therefore animals can undertake locomotion for daily living (**Figure 2A**) ^59–61^. The atrophy and recovery groups received a single infusion of TTX (6-7 d) followed by 9 d of TTX abstinence (e.g., TTX → recovery), whereas the repeated atrophy group received a second TTX infusion (5-6 d) following recovery (e.g., TTX → recovery → repeated TTX) (**Figure 2A**). The sham control group consisted of *n* = 1 for each timepoint (TTX/recovery/repeated TTX) that received the same surgical procedures (detailed below) as the experimental groups in the absence of TTX administration. Pre- and post-intervention animal weights for each group are presented in **supplementary Table 1** and muscle weights are provided in **Supplementary Figure 3.** Rats were socially housed in pairs throughout the experiment in accordance with best welfare practice, which precluded precise measurement of individual nutritional intake. Body weight was recorded at each timepoint, and wet food was provided when required. Body weight remained stable during the atrophy and recovery phases and decreased only after repeated atrophy (Supplementary Table 1), indicating that alterations in recovery was not attributable to reduced nutritional intake. Objective assessment of dorsiflexion in very old, caged rats is challenging; therefore, intervention efficacy and specificity were verified using: (i) transcriptomic signatures that followed the application and withdrawal of TTX, (ii) paired comparisons between treated and contralateral control muscles within each animal, (iii) selective reductions in dorsiflexor muscle mass, with minimal changes in adjacent plantarflexors, and (iv) saline-infused sham controls that confirmed surgical procedures alone did not affect muscle mass or gene expression.

**Figure 2.**
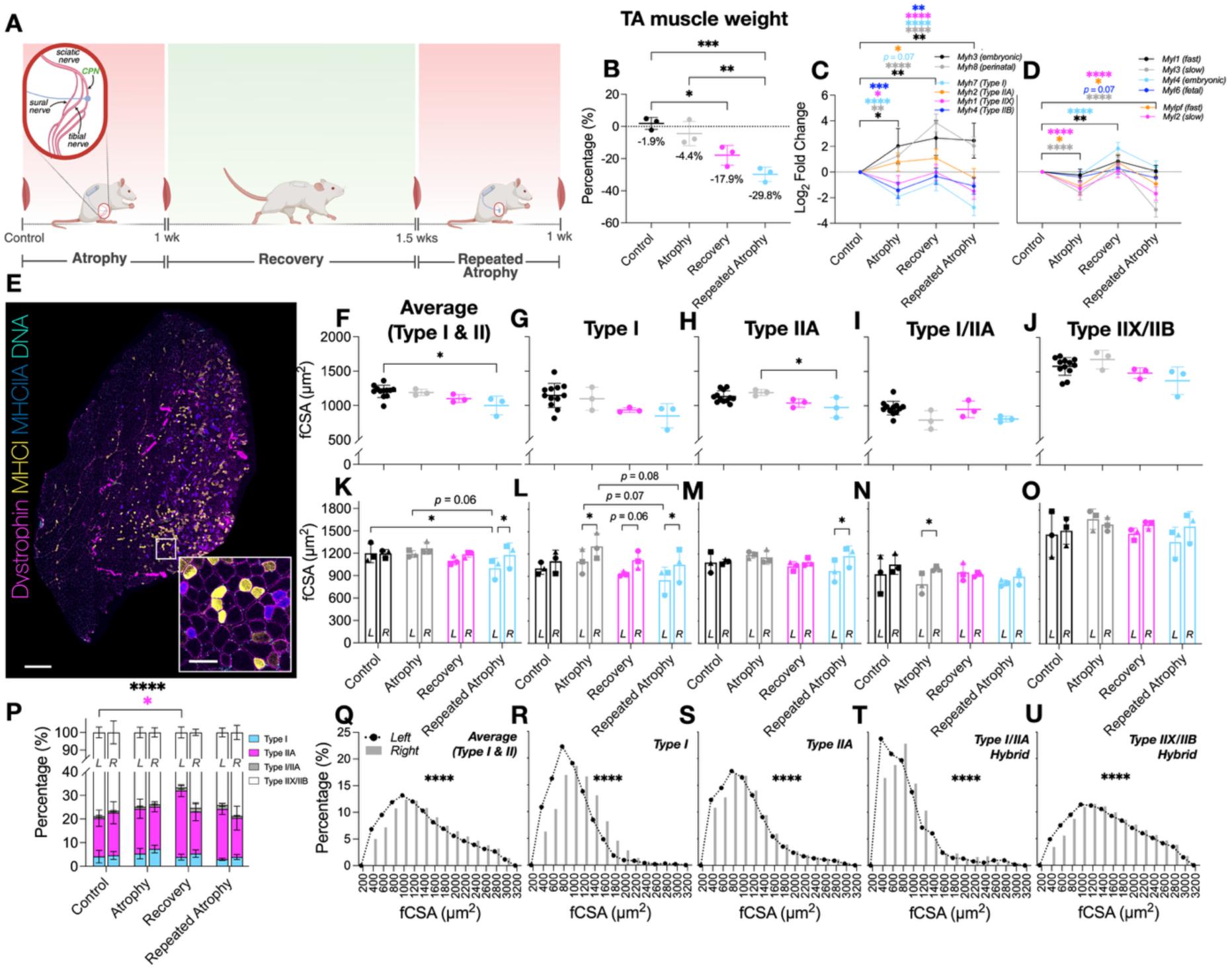
(**A**) Schematic representation of repeated disuse model in aged animals (created with BioRender.com). CPN = common peroneal nerve. (**B**) Reduction in TA muscle weight during atrophy and repeated atrophy with lack of recovery (presented as % change of the left TTX versus right control limb normalised to body weight). (**C**) Myosin heavy chain transcriptional analysis revealed significant changes in developmental (*Myh3/8*) and adult (*Myh7/2/1/*4) isoforms after atrophy and repeated atrophy (**D**) Essential (*Myl1/3/4/6*) and regulatory (*Mylpf/2*) myosin light chains were also differentially expressed after atrophy, recovery and/or repeated atrophy yet most were restored during recovery, despite continued muscle loss (**E**) Representative fluorescence microscopic image of whole TA muscle section labelled with dystrophin (magenta), MyHCI (yellow), MyHCIIA (blue), DNA (cyan). Non-staining represents putative type IIX/IIB myofibres. Tile scan images acquired at 10× magnification (scale bar within scan image = 500 µm, scale bar within magnified inset image = 100 µm). (**F-O**) There were significant reductions in fCSA of predominantly slow fibres after atrophy (pure type I and I/IIA hybrid fibres) and repeated atrophy (average, type I and IIA fibres). Symbol shapes represent individual animals. (**P**) Proportional fibre type analysis revealed greater contribution of type IIA and reduced number of IIX/IIB after recovery versus control in the left surgical limb. (**Q-U**) Fibre size distribution analysis demonstrates a leftward shift and larger frequency of smaller (<1000 µm^2^) fibres in the left TTX versus right control limb. \**p* ≤ 0.05, \*\**p* ≤ 0.01, \*\*\**p* ≤ 0.001. fCSA = myofibre cross-sectional area.

### Animal experimental procedures

Surgical and TTX-administration procedures are described elsewhere ^66, 72^. Briefly, animals were first anaesthetized via inhalation of a gaseous mixture of 3% isoflurane in oxygen for induction and 1–2% for maintenance prior to subcutaneous implantation of a mini osmotic pump (Mini Osmotic Pump 2002; Alzet, Cupertino, CA) in the scapular region. Silicone tubing attached to the mini osmotic pump was then passed under the skin to the site of the left common peroneal nerve (CPN). A second incision was made laterally through the skin, just proximal to the knee joint and through the biceps femoris muscle (posterior compartment of the thigh) to enable access to the CPN. A silicone cuff extending from the silicone tubing was placed around the nerve. All incisions were closed in layers. The mini osmotic pump (Mini Osmotic Pump 2002; Alzet, Cupertino, CA) delivered TTX to the CPN. The osmotic pump successfully delivered 0.5 μl/hr TTX (350 mg/mL in sterile 0.9% saline), continuously blocking ankle dorsiflexion for a 7d period after which the TTX was exhausted, while maintaining normal voluntary plantarflexion via the tibial nerve. The nature of the osmotic pump means that delivery of its contents starts as soon as the pump is implanted; therefore, silencing of the common peroneal nerve was present immediately after surgery. TTX was not administered to the right limb, and it served as a contralateral control for each individual animal. For the repeated disuse atrophy experiments, the rats were anaesthetized briefly after the recovery period. The empty osmotic pump was removed through an interscapular incision and replaced with a new TTX-loaded pump. The welfare and mobility of the rats were checked daily by animal welfare staff. Rats were euthanised with a rising concentration of carbon dioxide in oxygen until cessation of breathing followed by cervical dislocation. All rat tissue collections were performed at a consistent time of day between the hours of 08.00 and 13.00 across all experimental conditions, during the early inactive phase. The scheduling of operating days and subsequent harvesting days introduced no systematic bias for muscle from one group to be harvested consistently earlier or later than muscle from another group across all experimental conditions. Muscles were harvested immediately after euthanasia, weighed, and cut into transverse pieces approximately 2 mm thick. Pieces from the left and right limbs were mounted on the same piece of cork and frozen in pre-cooled isopentane to ensure simultaneous processing of experimental and contralateral control samples. Serial transverse sections from the same block were then used for all downstream analyses, including nucleic acids, protein, and IHC.

### Human muscle derived stem cell (MuSC) experimental procedures

Isolation of human muscle derived stem cells (MuSC) and subsequent cell culture procedures are previously described by our group ^59, 72^. Briefly, pre- and post-atrophy muscle biopsies (*n* = 4 of pre- and post-biopsies) were immediately transferred to a sterilised class II culture hood (Thermo Scientific MSC-advantage) in low serum transfer media (components described above). Visible fat and connective tissue were removed, and remaining muscle tissue was washed 3 × PBS (1× with added PS and AB), before undergoing repeated rounds of scissor mincing in trypsin (0.5%)-EDTA (0.2%) and centrifugation (340 × *g* for 5 min at 24 °C) to promote cell migration. The supernatant, cell pellet and remaining non-homogenised tissue were seeded into separate T25 pre-gelatinised flasks containing 5 mL growth media (GM: Ham’s F-10, 10% hiFBS, 10% heat inactivated new born calf serum/hiNBCS, 2 mM LG, PS and AB) and left unagitated in a humidified incubator (37 °C, 5%, VWR ILCO 180 Premium) for 5-7 d to promote cell growth. GM was changed every 2-3 d thereafter until sufficient cell attachment and growth (<80% confluency within cell colonies) was achieved. MuSC were transferred and sub-cultured in T75 flasks to further increase cell yield and ensure sufficient MuSC numbers for subsequent experiments described below.

To determine the effects of disuse *in vivo* on differentiation capacity *in vitro*, both pre- and post-atrophy MuSC (7 × 10^4^ cells/mL in 2 mL of GM) were seeded onto pre-gelatinised six-well plates and left until ≈ 80% confluency was attained (taking between 24 and 48 hrs). Cells were then washed 3 × PBS (1×) and media was switched to low serum media (DM: same components as GM with lower 2% hiFBS) to permit differentiation over 10 days (d). An additional 1 mL DM was added at 72 hrs and media was switched to DM containing either 0.5 mM nicotinamide riboside (NR, ChromaDex Inc, USA), previously used by out group ^73^ or control (H_2_O) for both pre- and post-atrophy conditions. Cell pellets derived from 4 × wells at 0 hrs, 7 and 10 d (+/- NR supplementation) were snap-frozen and stored at -80 °C for later copurification of DNA (2 ± 0.19_A260/A280_/0.14 ± 0.14_A260/A230_) and RNA (2.05 ± 0.03_A260/A280_/1.86 ± 0.48_A260/A230_) using the AllPrep RNA/DNA Mini Kit (Qiagen). The remaining 2 × wells were treated for subsequent immunocytochemistry analysis described below. There was no difference in the proportion of myoblasts (42 ± 22% versus 45 ± 26%, *p* = 0.62) and fibroblast (58 ± 22% versus 56 ± 26%, *p* = 0.62) in pre- and post-atrophy MuSC, respectively. Thus, MuSC were matched prior to differentiation experiments. All MuSC were between passage 3 and 6 to avoid issues of cellular senescence ^74^. No mycoplasma was identified during primary cell cultures.

### Immuno-histo/cytochemistry, imaging and analysis

Serial sections at either 8 μm (human VL muscle) or 10 μm (mid-belly of the rat TA muscle) thickness were made at −20 °C (CM1860, Leica Biosystems GmbH, Heidelberger, Germany), mounted on microscope slides (Fisherbrand™ Superfrost™ Plus Microscope Slides, Fisher Scientific), air-dried for 30 mins and stored at −80 °C. When labelling muscle tissue samples, samples were first air-dried for 30 mins prior to blocking in 2% bovine serum albumin (BSA, Sigma-Aldrich) in 1× PBS (10 mM phosphate buffer, 3 mM KCl, pH 7.4, Merck, Oslo, Norway) with 0.05% added tween (PBS-T, Tween^®^ 20, VWR, Oslo, Norway) and 1% fat-free dry milk (Sigma-Aldrich) for 1 hr at room temperature (RT). Muscle cross-sections were then counterstained for dystrophin (1:500; ab15277, Abcam, Cambridge, UK), myosin heavy chain (MyHC) type I (1:100; BA-F8, DSHB Supernatant, Schiaffino, S, Iowa, USA) and MyHC type IIA (1:100; SC-71, DSHB Supernatant, Schiaffino, S, Iowa, USA) diluted in 0.05% PBS-T and 1% milk overnight at 4 °C (**Table 2**). After overnight incubation, samples were labelled with goat anti-rabbit (GaR) IgG H+L (1:200; Alexa Fluor™ 488, cat. #A-11008 or Alexa Fluor™ 633, cat. #A-21070, Thermo Fisher Scientific, Waltham, MA, USA), goat anti-mouse (GaM) IgG1 (1:200; Alexa Fluor™ 488, cat. #A-21121) and/or GaM IgG2b (1:200; Alexa Fluor™ 594, cat. #A-21145) diluted in 0.05% PBS-T and 1% milk for 1 hr and RT (**Table 2**). Slides were washed 3 × 5 mins in 0.05% PBS-T between each blocking and AB labelling steps. Finally, coverslips were mounted using ProLong™ Gold Antifade Mountant with 4′,6-diamidino-2-phenylindole/DAPI (cat. #P36935, Thermo Fisher Scientific) and left to air-dry at RT overnight prior to imaging. For imaging, tile scan images (1024×1024 frame size) of whole cross-sections were acquired using a ZEISS LSM 800 (with Airyscan) confocal microscope (ZEISS, Oberkochen, Germany) equipped with a 10×/0.3 objective and subsequently processed using either ZEN 2 (blue edition, version 2.6) or ImageJ2 (version 2.14.0/1.54f, U. S. National Institutes of Health, Bethesda, Maryland, USA) software. Automated analysis of myofibre cross-sectional area (fCSA) and type was conducted using MyoVision 2.0 software ^59^. Issues surrounding muscle tissue processing in the human muscle (e.g., biopsy size, freeze damage) resulted in variable sample numbers for average (type I and II combined; *n* = 8 for baseline/atrophy, *n =* 7 for recovery, *n* = 6 for repeated atrophy) and type I/II (*n* = 7 baseline, *n* = 8 atrophy, *n =* 6 recovery, *n* = 5 repeated atrophy) myofibre analysis. The total number of fibres analysed from human biopsies was 398 ± 331 (average), 149 ± 94 (type I) and 281 ± 77 (type II). The total number of fibres analysed from whole rat muscle was 6297 ± 2067 (average), 280 ± 109 (type I), 1237 ± 546 (type IIA), 69 ± 48 (type I/IIA), 4712 ± 1613 (type IIX/IIB).

**Table 2.**
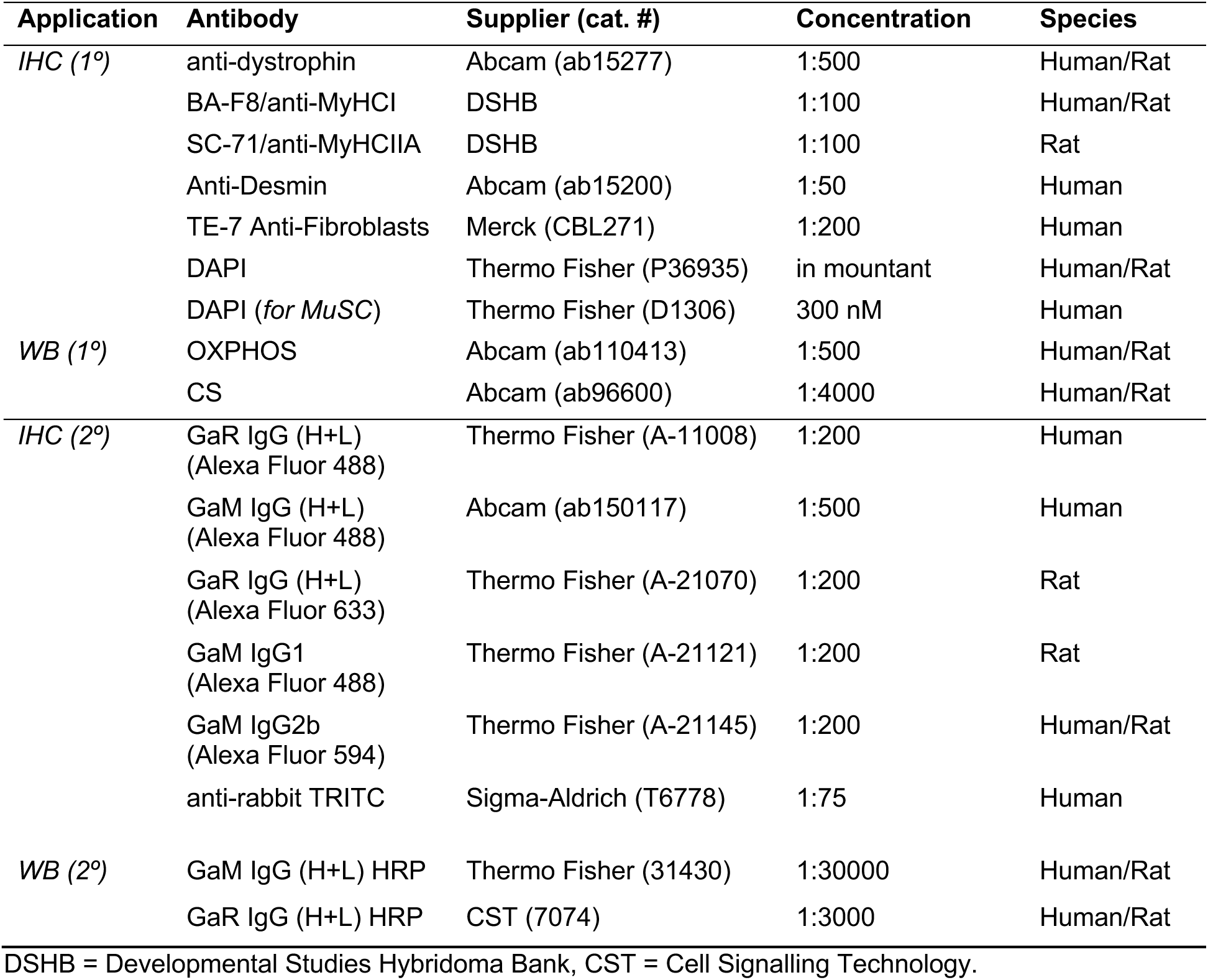
Primary (1°) and secondary (2°) antibodies used for immunohistochemistry (IHC) and Western blot (WB) procedures.

For characterising myogenic (% myoblasts and fibroblasts) proportions and myotube differentiation, MuSC were washed 3 × TBS (1×), fixed in 4% PFA for 15 mins, incubated in blocking/permeabilization solution (5% goat serum/GS and 0.2% Triton X-100 in 1× TBS) for 1 hr at room temperature (RT). Cells were then washed and incubated overnight (4 °C on a rocker) in Ab solution (2% GS and 0.2% Triton X-100 in 1× TBS) containing anti-desmin (1:50; ab15200, Abcam) and TE-7 (1:100; CBL271, Merck) primary Ab for counterstaining myoblasts and fibroblasts, respectively. After overnight incubation, cells were washed 3 × TBS and incubated at RT for 3 hrs (4 °C on a rocker) in secondary Ab solution containing anti-rabbit IgG TRITC (1:75; T6778, Sigma-Aldrich) for myoblasts and GaM Alexa Fluor^®^ 488 (1:500; ab150117, Abcam) for fibroblasts. Finally, cells were washed 3 × TBS, prior to counterstaining nuclei using DAPI (300 nM; D1306, Thermo Fisher Scientific) for 30 mins at RT. DAPI was removed and stained cells were washed and stored in TBS for later image acquisition. The same microscope and acquisition settings used for imaging tissue sections above were used when imaging MuSC (2 × duplicate wells per condition/timepoint) except for acquiring nine (3 × 3) tile scan images. The ImageJ2 cell counter was used to quantify the number of myoblasts and fibroblasts. The polygon tool was used to outline the sarcolemma of all myotubes (cells with ≥3 nuclei) for assessing myotube area (µm^2^).

### RNA extraction from muscle tissue

Skeletal muscle tissue derived from humans (33 ± 6 mg) and animals (27 ± 4 mg) was added to 2 mL homogenizing mix tubes (Fisherbrand™, Fisher Scientific) containing 600 μl Buffer RLT (Qiagen) with added 1% β-Mercaptoethanol (β-ME) before transferring to a tissue homogenizing instrument (MagNA Lyser; Roche) for subsequent homogenization (3 × 45 s at 6000 rpm, placing samples on ice for 5 mins between each disruption). Following homogenization, RNA was purified using a RNeasy^®^ Mini kit (Qiagen) and treated with DNase (RNAse-Free DNase Set, Qiagen) as per the manufacturer’s instructions. RNA quantity and purity for both human (128 ± 175 ng/mg tissue, 1.9 ± 0.1_A260/A280_/2.1 ± 0.2_A260/A230_) and rat (406 ± 215 ng/mg tissue, 2.1 ± 0_A260/A280_/2.2 ± 0.2_A260/A230_) samples were quantified via spectrophotometry (QIAxpert^®^, software version 2.4, Qiagen) and concentrations confirmed via fluorometry (Qubit^®^, Qiagen). RNA integrity prior to sequencing was confirmed via Agilent Bioanalyser 2100 with RIN scores 7.22 + 0.36 for human and 8.29 + 0.62 for rat samples (detailed below).

### qPCR for targeted mRNA analysis of aged rat muscle tissue and human MuSC

Methods for mRNA expression analysis via qPCR by our group have been explained elsewhere ^40, 48, 49, 59^. Briefly, a one-step QuantiFast SYBR Green kit (Qiagen) was used for preparation of PCR reactions. Each 10 μl reaction contained 4.75 μl experimental sample (7.36 ng/μl totalling 35 ng per reaction), 0.075 μl of both forward and reverse primer (100 μM stock) of the genes of interest (primer details are provided in **Table 3**), 0.1 μl of reverse transcriptase and 5 μl of SYBR Green Master Mix (totalling reactions). A thermocycler (Rotorgene 3000Q, Qiagen) was used to amplify target transcripts. Reverse transcription was initiated with a hold at 50 °C for 10 mins (cDNA synthesis) and 5 mins at 95 °C (transcriptase inactivation and initial denaturation), before 40 cycles of; 95 °C for 10 s (denaturation) followed by 60 °C for 30 s (annealing and extension). Only intended targets (via BLAST search) were identified, thus yielding a single peak after melt curve analysis. All relative gene expression was quantified using the comparative Ct (^ΔΔ^Ct) method ^75^. For *Nmrk2* expression analysis of aged animal muscle tissue, C_T_ values (mean of technical duplicates) for each condition (atrophy/recovery/repeated atrophy) were relativised to pooled C_T_ values from the control limb (*n* = 9) and reference gene, *Pol2ra* (19.27 ± 0.41, 2.12% CV). For temporal mRNA analysis of *MRF* and *NMRK* genes in human MuSC, C_T_ values (mean of technical duplicates) from pre- and post-atrophy MuSC at 7 and 10 d (+/- NR) were relativised to own 0 hr C_T_ value and pooled reference gene, *RPL13A* (14.90 ± 0.43, 2.87% CV). To determine the effects of NR supplementation, C_T_ values from 10 d +NR were relativised 10 d -NR and pooled *RPL13A* values for both pre- and post-atrophy timepoints. Primers were designed using Basic Local Alignment Search Tool (BLAST; http://blast.ncbi.nlm.nih.gov/) and Clustal Omega (https://www.ebi.ac.uk/Tools/msa/clustalo/) and purchased from Sigma.

**Table 3:**
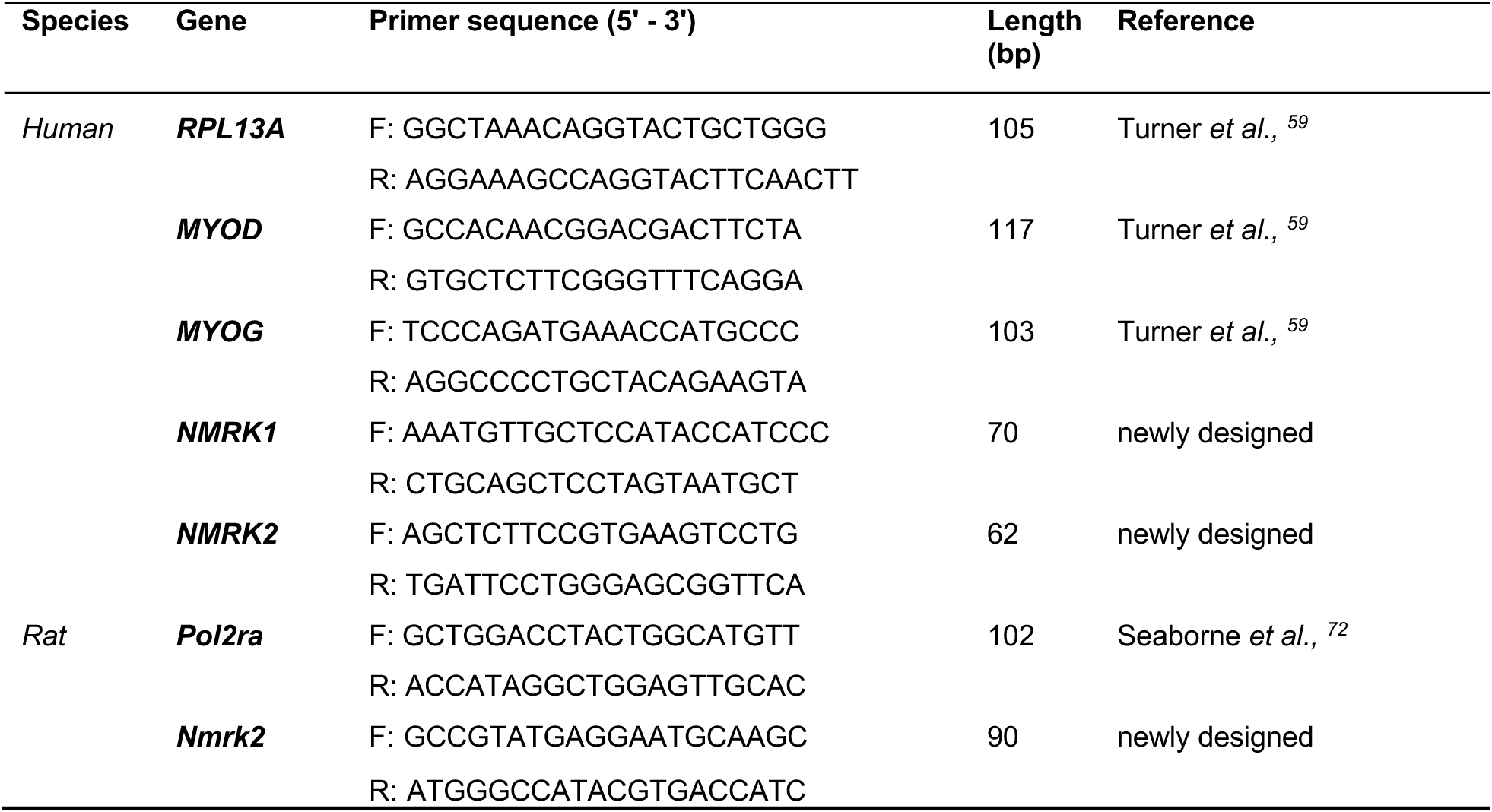
qPCR primers.

### RNA-seq and associated bioinformatic analysis

Total RNA (100 ng) from each sample was used to prepare mRNA libraries using the NEBNext mRNA isolation kit in conjunction with the NEBNext Ultra II Directional RNA Library preparation kit (New England Biolabs, Massachusetts, USA), and samples were randomised before preparation. Fragmentation of isolated mRNA prior to first strand cDNA synthesis was carried out using incubation conditions recommended by the manufacturer for an insert size of 300 bp (94 °C for 10 minutes). 13 cycles of PCR were performed for final library amplification. Resulting libraries were quantified using the Qubit 4.0 spectrophotometer (Life Technologies, California, USA) and average fragment size assessed using the Agilent 4200 Tapestation with D1000 screentape (Agilent Technologies, Waldbronn, Germany). A final sequencing pool was created for each project using equimolar quantities of each sample with compatible indexes. 59 bp paired-end reads were generated for each library using the Illumina NextSeq 2000 in conjunction with the NextSeq 2000 P3 100 cycle kit (Illumina Inc., Cambridge, UK). Resulting FastQ files were imported and processed in Partek Flow^®^ software (version 12.4.0; Partek, Inc. Missouri, USA) to filter contaminants (rDNA/tRNA/mtrDNA) using `bowtie2` (v2.2.5) and align reads to the hg38 *Homo sapiens* (release 39) and rn7 *Rattus norvegicus* genome assemblies using Spliced Transcripts Alignment to a Reference (`STAR`) alignment tool (v2.7.3a). Total number of pre/post-aligned reads was 35/34 million and 49/49 million for human (65 bp average read length) and rat (59 bp average read length) samples, respectively. Phred scores were >33 for both human and rat RNA samples, thus trimming and filtering of reads was not required. Normalised differential gene expression analysis was performed using *DESeq2* median of ratios within the Partek Flow^®^ software. Principal component analysis (PCA), cluster heatmaps and volcano plots were created in R studio.

Venn diagrams were redrawn from the open-source `DeepVenn` web application. Pathway (KEGG and REACTOME) and gene ontology (GO; cellular component/CC, molecular function/MF and biological process/BP) performed via the Database for Annotation, Visualization, and Integrated Discovery (`DAVID`) knowledgebase and associated figures created in R studio. Custom background gene lists of detected genes following *DESeq2* normalization were used for functional annotation analysis. For human analysis, normalised reads for each timepoint (atrophy/recovery/repeated atrophy) were relativised to baseline. For rat samples, all experimental conditions in the left limb (atrophy/recovery/repeated atrophy) were relativised to pooled right contralateral non-surgical limb. Given there were no DEGs detected between sham and pooled contralateral control limb samples, we excluded sham samples from further analysis and pooled contralateral non-surgical limb samples for the control group. Genes with a ≥20% fold change (FC) and false discovery rate (FDR) of ≤5% are considered differentially expressed genes (DEGs). Data are presented as either log_2_ or FC ± confidence intervals (CI). Genes with negative changes presented as signed FC (-1/ratio). To perform temporal clustering of the identified DEGs and DMRs (described below), self-organising map (SOM) profiling was conducted using Partek Genomics suite, as previously described ^38, 40, 46, 59^. SOM analysis plots a standardised mean (z-score normalisation) shifting the mean to a value of 0 and scaling to a standard deviation of 1. Therefore, by subtracting the mean of the group this centres the data around zero and then dividing by the standard deviation, scales the data so the spread is consistent, allowing removal of the absolute differences and focusing on relative changes allowing comparison of gene transcripts and DMRs on a common scale. The mapping then clusters DEGs and DMRs based on similar temporal profiles allowing visualisation of temporal signatures over time relative to the group mean. Also, this allows the temporal signature of the DEGs and DMRs on common genes to be compared on the same scale over the time-course of the intervention. All `package` links and references are available in **Supplementary Table 2.**

### DNA extraction from muscle tissue

Skeletal muscle tissue derived from humans (21 ± 3 mg) and animals (12 ± 2 mg) was added to 2 mL homogenizing mix tubes (Fisherbrand™, Fisher Scientific) containing 180 μl Buffer ATL (Qiagen, Hilden, Germany) before transferring to a tissue homogenizing instrument (MagNA Lyser; Roche, Basel, Switzerland) for subsequent homogenization (3 × 45 s at 6000 rpm, placing samples on ice for 5 mins between each disruption). Following homogenization, 20 μl proteinase K (Qiagen) was added and samples were incubated at 56°C until complete lysis (≈ 30-60 mins) before continuing with DNA purification using a DNeasy^®^ Blood & Tissue kit (Qiagen) as per the manufacturer’s instructions. DNA quantity and quality for both human (251 ± 47 ng/mg tissue, 2 ± 0_A260/A280_/2.1 ± 0.1_A260/A230_) and rat (1427 ± 426 ng/mg tissue, 2 ± 0_A260/A280_/2 ± 0.1_A260/A230_) samples were quantified using spectrophotometry (NanoDrop™ 2000; Thermo Fisher Scientific) and concentrations confirmed via fluorometry (Qubit^®^, Qiagen). DNA integrity was confirmed prior to sequencing via Agilent Bioanalyser 2100 with DIN scores 7.11 ± 0.17 for human and 7.22 ± 0.23 for rat samples.

### Reduced Representation Bisulfite Sequencing (RRBS): Library preparation and sequencing

RRBS libraries were prepared from 100 ng of genomic DNA using the Diagenode Premium RRBS v2 library preparation kit, as per the manufacturer’s protocol. For human samples, two groups of 8 samples, and three groups of 7 samples were prepared after qPCR quantification of adapter-ligated sample and prior to bisulphite conversion. 14 cycles of final library amplification were performed after assessment of bisulphite converted library pool by qPCR. For rat samples RREM (reduced-representation enzymatic methyl-seq) libraries were prepared using 100 ng of genomic DNA in conjunction with the NEBNext Enzymatic Methyl-Seq kit (New England Biolabs, Massachusetts, USA). Briefly, DNA samples were digested using MspI restriction enzyme followed by end-repair and adapter ligation. A two-step enzymatic conversion was then performed by Oxidation of 5mC and 5hmC using TET2 and glucosylation of 5hmC using T4-BGT. Deamination of unmodified cytosines was subsequently performed using APOBEC. Converted samples were then uniquely barcoded and amplified by PCR prior to sequencing. Libraries were sequenced using the Illumina NextSeq 2000 with P3 SBS 100-cycle sequencing kit for human, and with P4 XLEAP 100-cycle sequencing kit for rat, both generating 50 bp paired-end reads.

### RRBS quality control, trimming, alignment and methylation calling

Reduced representation bisulfite sequencing (RRBS) libraries were processed using a standard workflow. Raw base-call files were demultiplexed using Illumina `bcl2fastq /2.19.1`, and unique molecular identifiers (UMIs) were extracted using `umi_tools`. Reads were adapter and quality-trimmed (minimum phred score 30) with `trim_galore` (in RRBS mode, reads less than 15 bp after trimming were discarded. Trimmed reads were aligned to the human reference genome (hg38) or rat genome Rn6 using `bismark/0.22.1` with `bowtie2`, with the PBAT option enabled. PCR duplicates were removed with `deduplicate_bismark` in a UMI-aware manner. CpG methylation calls were obtained using `bismark_methylation_extractor`. All `package` links and references are available in **Supplementary Table 2.**

### RRBS: Identification of differentially methylated positions (DMPs) and regions (DMRs)

Downstream analysis was performed in R. Per-CpG coverage files were imported into `methylKit` with sites of low (<10 reads) or extreme coverage (top 0.1%) removed and outlier samples excluded. For humans, data from each sample was then merged based on only CpGs present in at least 6 samples. Each of our groups contained 7-8 samples after outlier removal, making this threshold a minimum of 75% of samples and giving a total of ≈ 1.8 million total CpGs analysed. For rats, there were 9 samples (3 per experimental condition of atrophy, recovery and repeated atrophy) and 9 samples for the contralateral control limbs, therefore data from each sample were merged based on CpGs present in all samples (100%), giving a total of 29K CpGs analysed. Differential methylation was identified at individual CpGs (DMPs) and then 1-kb tiled regions (DMRs) using `calculateDiffMeth`. The percentage methylation of each CpG was calculated using the `percMethylation` function in `methylKit` and during differential analysis the mean percentage methylation of each CpG was calculated for each group. For DMRs, mean percentage methylation for each experimental group was calculated by averaging the percentage methylation of all CpGs within a given DMR across all samples from the given group. For humans, we performed 6 pairwise comparisons of: 1) Atrophy vs. Baseline, 2) Recovery vs. Baseline, 3) Repeated Atrophy vs. Baseline, 4) Recovery vs. Atrophy, 5) Repeated Atrophy vs. Recovery, 6) Repeated Atrophy vs. atrophy. For rat analysis we performed: 1) Atrophy vs. Control, 2) Recovery vs. Control, 3) Repeated atrophy vs. Control, 4) Recovery vs. Atrophy, 5) Repeated Atrophy vs. Recovery, 6) Repeated atrophy vs. atrophy. For the rats, as with the transcriptome data, control was determined as the pooled right/non-surgical limb (contralateral control limb = 9). Differentially methylated CpGs and regions were annotated to genes and regulatory features for humans using `TxDb.Hsapiens.UCSC.hg38.knownGene`, `org.Hs.eg.db` and `annotatr` and in rats using `TxDb.Rnorvegicus.UCSC.rn6.refGene`, `rg.Rn.eg.db` and `annotatr.` Default settings were used unless otherwise specified. Only DMRs with FDR ≤ 0.05 were considered significant. SOM clustering (described above) was performed on significant DMRs using percentage difference methylation as this allowed methylation values at baseline (human) and control (rat) to enable temporal profiling. All package links and references are available in **Supplementary Table 2.**

### Mitochondrial DNA content

Mitochondrial DNA (mtDNA) content was assessed via real time quantitative polymerase chain reaction (RT-qPCR). PCR primers encoding for nuclear DNA (nDNA) markers; β_2_ microglobulin (*B2M*), hemoglobin subunit beta *(HBB)*, tubulin-specific chaperone A (*Tbca*), glyceraldehyde 3-phosphate dehydrogenase (*Gapdh*), and mtDNA markers; mitochondrial 16S rRNA (*MT-RNR2*), D-loop region (*DLOOP*), cytochrome B (*CYTB*), mitochondrially encoded 16S rRNA (*mt-Rnr2*) and 12S ribosomal RNA (*mt-Rnr1),* were used to determine nuclear and mitochondrial DNA copy number, respectively. Primer information is presented in **Table 4**. For qPCR, 20 ng DNA (9.6 μl at 2.08 ng/μl in nuclease-free H_2_O) was added to 10 μl SYBR^®^ Green Mastermix (Qiagen) and 0.2 μl primers (100 μM stock, 1 μM final concentration) to ensure 20 μl reaction volume. DNA was then amplified using a Rotor-Gene Q 5-Plex HRM thermocycler (Qiagen) with supporting software (version 2.3.1). Thermal cycle settings consisted of 3 mins at 95 °C (initial denaturation), followed by 40 cycles of 10 s at 95 °C, 30 s at 62 °C (annealing) and final extension of 10 s at 95 °C. Melt curve analysis was performed at 0.5 °C increments from 65 to 95 °C. Mitochondrial DNA content was reported as the ratio of mtDNA (*16S, CYTB* and/or *DLOOP* for human; *mt-Rnr2* and/or *mt-Rnr1* for rat) to pooled nDNA (*B2M/HBB* for human; *Tbca/Gapdh* for rat) determined using the ^ΔΔ^Ct method ^75, 76^.

**Table 4.**
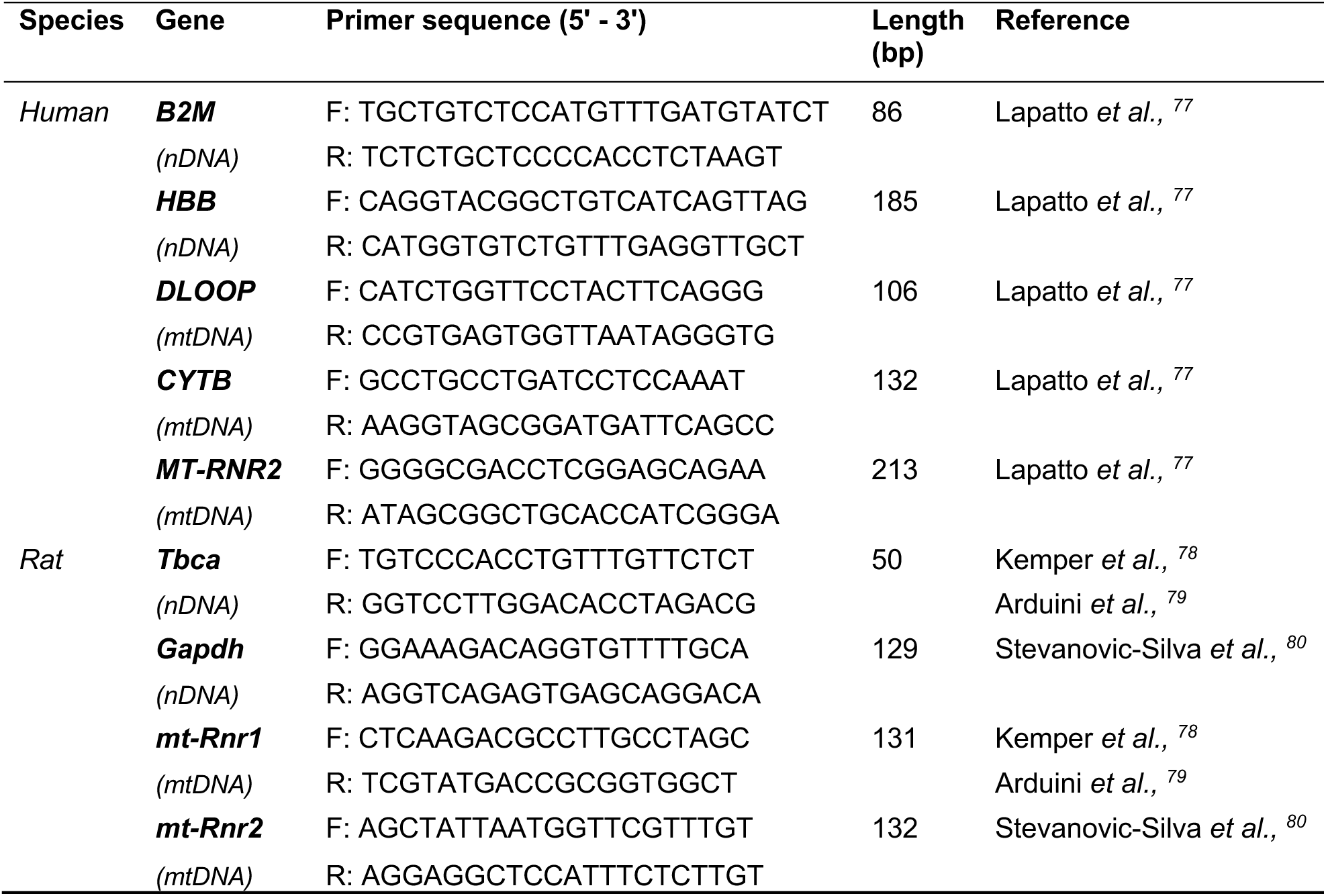
Primers used for encoding for nuclear DNA (nDNA) and mitochondrial DNA (mtDNA) markers for assessment of mtDNA content.

### Protein extraction and Western blot

Skeletal muscle tissue derived from humans (21 ± 6 mg) and rats (38 ± 14 mg) was added to 2 mL homogenizing mix tubes (Fisherbrand™, Fisher Scientific) containing either 1 mL RIPA buffer (Thermo Fisher Scientific) with added protease and phosphatase inhibitors (Halt™ inhibitor cocktail, Thermo Fisher Scientific) for human samples or sucrose lysis buffer (250 mM sucrose, 50 mM Tris pH 7.5, 1 mM EDTA, 1 mM EGTA, 1% Triton™ X-100, Sigma-Aldrich) with added inhibitors (Thermo Scientific™ Pierce Mini Tablets, Thermo Fisher Scientific) for rat samples. Tubes were then transferred to a MagNA Lyser instrument (Roche) for subsequent homogenization (3 × 45 s at 6000 rpm, placing samples on ice for 5 mins between each disruption). Following homogenization, protein concentration was quantified for both human (73 ± 13 µg/mg) and rat (84 ± 18 µg/mg) tissue via the bicinchoninic acid (BCA) method (Pierce™ BCA protein assay kit, Thermo Fisher Scientific) and a FLUOstar Omega microplate reader (BMG LABTECH, Ortenberg, Germany) with supporting software (Omega version 1.3). For Western blotting, protein aliquots were first diluted in 4× Laemmli Sample Buffer (Bio-Rad) with added 0.5 M Dithiothreitol (DTT, Bio-Rad) and heated at either 45°C (oxidative phosphorylation, OXPHOS) or 70°C (citrate synthase, CS) for 10 mins. Following denaturation, 20 or 25 µg for all protein samples (in 15 µL for human and 20 µL for rat) and 5 µL protein marker (10-250 kDa, Precision Plus Protein™ All Blue Prestained Protein Standards, Bio-Rad) were loaded into pre-cast gels (12 or 15 well, 4–20% Mini-PROTEAN^®^ TGX Stain-Free™, Bio-Rad) before undergoing protein separation via electrophoresis (200 volts/V for 30 mins). The current was monitored (A) pre- and post-electrophoresis. Following electrophoresis, gels were activated under UV light ChemiDoc MP Imaging System (Bio-Rad) and proteins transferred to PVDF nitrocellulose membranes (0.2 µm, Bio-Rad) using transfer stacks (Bio-Rad) and a Trans-Blot Turbo Transfer System (Bio-Rad). Following protein transfer, membranes were blocked (5% milk in 1% PBS-T for 1 hr at RT) and subsequently labelled with primary antibodies against mitochondrial OXPHOS complexes (CI-NDUFB8, CII-SDHB, CIII-UQCRC2, CIV-MTCO1, CV-ATP5A) (1:500, cat. #ab110413, Abcam) and CS (1:4000, cat. #ab96600, Abcam). Primary Ab were diluted in 1% milk in 1% PBS-T and incubated on a roller overnight at 4 °C. After overnight incubation, OXPHOS and CS proteins were labelled with secondary antibodies, GaM IgG (H+L) HRP (1:30000, cat. #31430, Thermo Fisher Scientific) and GaR IgG (H+L) HRP (1:3000, cat. #7074, Cell Signaling Technology), respectively. Membranes were washed (3 × 5 mins in 1% PBS-T) between blocking and antibody labelling steps. Finally, membranes were incubated in enhanced chemiluminescence (ECL) HRP substrate (SuperSignal™ West Dura Extended Duration Substrate, Thermo Scientific) for 5 mins at RT to enable visualisation of membrane-bound proteins via chemiluminescence using a ChemiDoc MP system (Bio-Rad) and analysis of band intensities using supporting software (Image Lab 6.1, Bio-Rad). For human samples, all timepoints for each participant were loaded on the same gel and repeated to ensure duplicate samples across two separate gels. Band intensities were relativised to own baseline sample for subsequent timepoints (atrophy/recovery/repeated atrophy) and the average across the 2 gels was reported. Total protein normalisation was performed using Bio-Rad stain-free technology. Lane-specific normalisation factors were derived from UV-activated stain-free images and applied uniformly to all target proteins. Loading consistency was then assessed by expressing relative total protein as the percentage difference from each human participant’s baseline sample and, for animals, from each rat’s right contralateral control limb. Mean percentage differences across samples were 19% for OXPHOS and 20% for citrate synthase in humans, and 11% for OXPHOS and 18% for citrate synthase in rats. Representative total protein gel images are provided in the relevant figures. One human recovery sample was excluded from all analysis due to low detection of protein. Thus, there was *n* = 8 (rather than *n* = 9) samples for recovery timepoint. Similarly for aged animal samples, the left/surgical and right/non-surgical limbs for each animal and condition (excluding repeated atrophy as the protein lysates were low quality and quantity) were loaded on the same gel across two separate gels. Band intensities of left/surgical limb were relativised to thier own right/non-surgical limb for each condition (sham/atrophy/recovery). Control samples consisting of a mixture of all human or rat samples were loaded in duplicate within each gel to ensure consistent sample loading. There was no CII-SDHB band detected in 1 out of the 2 gels for both human and rat samples. Therefore, data is derived from only one gel for this protein.

### NAD^+^ measurements

NAD^+^ levels were measured using an enzymatic cycling assay ^81^. Briefly, 11 ± 1 mg (human) and 8 ± 2 mg (rat) muscle were lysed in 400 µl 0.6 M perchloric acid (HClO_4_) and homogenized using steel beads and a TissueLyser II (Qiagen). Following homogenisation, samples were centrifuged for 5 min at 13,000 *g.* The supernatant was transferred to new tubes and neutralized in 100 mM Na_2_HPO_4_ (1:300 for human, 1:800 for rat). For the assay, 100 µl of diluted extract was pipetted into a white 96-well plate after which 100 µl of reaction mix was added. The reaction mix contained: 100 mM Na_2_HPO_4_, 10 μM flavinmononucleotide, 2 % ethanol, 90 U/mL alcohol dehydrogenase, 130 mU/mL diaphorase, 2.5 μg/mL resazurin, and 10 mM nicotinamide. A Hidex Sense Microplate Reader (Hidex, Finland) was used to measure the fluorescence of accumulated resorfurin (Ex 544 nm/ Em 580 nm) for 30 min immediately after the reaction mix was added. Samples were run in duplicates and NAD^+^ content was calculated from the increase in fluorescence of a dilution series of NAD^+^. Values were normalized to protein concentration by dissolving the resultant pellets from the previous centrifugation step in 0.2 M NaOH via heating to 95°C. Protein concentrations were determined using the bicinchoninic acid (BCA) assay (Pierce™ BCA protein assay kit, Thermo Fisher Scientific).

### Statistical Analysis

A Shapiro-Wilk’s test of normality was first performed for all physiological (e.g., body weight, SkM mass, strength, mCSA and fCSA), protein abundance, mtDNA content and NAD^+^ measurements. A repeated (for human) or between (for aged animals) measures one-way analysis of variance (ANOVA) with Tukeýs HSD post hoc detected significant (*p* ≤ 0.05) differences whenever data was normally distributed. Whenever data was non-normally distributed, a Friedman (for human) or Krustal-Wallis (for aged animals) test with follow-up Dunńs post hoc was carried out. For human myofibre (fCSA and fibre type) analysis, a mixed-effect model was performed due to differences in sample number across timepoints. Fibre cross-sectional area (fCSA) frequency distributions were compared between left and right limbs using the Kolmogorov–Smirnov test on raw fibre-level data. Tests were performed per animal and fibre type, and p-values were combined across animals using Fisher’s method. For animal body weight (individual conditions), myofibre, mtDNA and NAD^+^ content analyses, a two-way ANOVA (within subject factor: limb, between subject factor: condition/timepoint) with Tukeýs HSD post hoc was performed. For mean animal body weight across all conditions, an unpaired t-test was performed. All statistical analysis was conducted in GraphPad Prism software for macOS (version 10.4.1, Massachusetts, USA) and associated figures were created using GraphPad or R studio for macOS (version 4.4.3). All data are presented as mean ± standard deviation (SD) unless otherwise stated in the figure legends. RNA-seq and RRBS data statistical analyses are described above. To combine RNA-seq and RRBS data, differentially expressed genes (DEGs) and differentially methylated regions (DMRs) were first identified using the approaches detailed in the RNA-seq analysis section and RRBS analysis section. These datasets were then integrated using self-organising map (SOM) profiling. This approach enabled temporal clustering of DEGs and DMRs on a common scale, allowing visualisation of coordinated inversely related transcriptional and epigenetic signatures across atrophy, recovery, and repeated atrophy.

## Results

### Physiological, muscle size, strength, muscle quality, myofibre structural, fibre-type, and myosin isoform expression responses to atrophy, recovery, and repeated atrophy in young adult humans

To characterise the physiological, muscle size, muscle strength, muscle quality, myofibre structural characteristics, fibre-type composition, and myosin isoform expression changes across atrophy, recovery, and repeated atrophy in young adult skeletal muscle, we first assessed leg lean mass, quadriceps anatomical cross-sectional area, knee extension strength, muscle quality (strength normalised to cross-sectional area), histological fibre cross-sectional area and fibre-type distribution, and the expression of key myosin light chain (MYL) and myosin heavy chain (MYH) genes. The first period of disuse-induced atrophy (hereafter referred to as “atrophy”) resulted in a significant -3.6% reduction in total leg lean mass of the immobilised limb (*p* = 0.005, **Figure 1B**). Upon returning to normal habitual activity (i.e., “recovery”), mass was fully restored to pre-atrophy baseline levels (*p* = *N.S* recovery vs. baseline). The second period of disuse (i.e., “repeated atrophy”) led to a similar significant -3.2% loss compared to previous recovery (*p* = 0.004, **Figure 1B**). Further assessment of *vastus lateralis (VL)* and *rectus femoris (RF)* anatomical muscle cross-sectional area (mCSA) in the immobilised limb revealed a -9% reduction in *VL* mCSA after initial atrophy (*p* = 0.0003), that was fully restored during recovery (recovery vs. baseline, *p* = *N.S,* **Figure 1C***). Vastus lateralis* mCSA was also significantly reduced after repeated atrophy relative to both baseline (−10.6%, *p* = 0.0003) and previous recovery (−8.9%, *p* = 0.002, **Figure 1C**). *Rectus Femoris* mCSA was only significantly reduced after repeated atrophy relative to recovery (−3.5%, *p* = 0.02, **Figure 1D**). There were no significant changes in the non-immobilised limb for total lean leg mass (**Supplementary Figure 1A-F)** or *VL* and *RF* mCSA (*p* = *N.S*, **Supplementary Figure 1 G-J**). There was however a significant increase in lower leg lean mass of the non-immobilised limb during both periods of disuse periods possibly due to a greater reliance on plantarflexor and dorsflexor muscles to support balance and locomotion during immobilisation of the opposite limb (**Supplementary Figure 1F**).

Concomitant with the loss of leg lean mass and quadriceps muscle size was a significant reduction in knee extension strength (assessed via MVC) at 90° (−14.3%, *p* = 0.02) and 60° (−12.5%, *p*= 0.02) knee flexion following initial atrophy that was also fully restored during recovery (*p* = *N.S*) **(Figure 1E & F)**. Repeated atrophy induced a similar loss of strength at 90° (−13.8%, *p* = 0.02) and 60° (−14.3%, *p* = 0.002) following recovery (**Figure 1E & F**). Despite this loss of muscle strength at 60° and 90°, there were no significant changes in knee extension MVC at 30° knee flexion (*p* = *N.S*, **Figure 1G**). Concurrent reductions in size and strength also resulted in impaired muscle quality (e.g., specific force by normalizing MVC to mCSA) (**Supplementary Figure 2A-I**). Specifically, MVC at 90° knee flexion when normalised to *RF* mCSA was significantly reduced after atrophy (−12.8%, *p* = 0.05, **Supplementary Figure 2D),** where repeated atrophy led to the greatest loss of specific force at 60° when normalised to *VL* (−7.1%, *p* = 0.03), *RF* (−12.2%, *p* = 0.04) and *VL+RF* mCSA combined (−8.2%, *p* = 0.02, **Supplementary Figure 2B & E & H** respectively). Finally, repeated atrophy led to significantly reduced knee flexion torque at 90° (*p* = 0.03, **Supplementary Figure 2J**). Mean MVC at 60° and 30° knee flexion also demonstrated the same trend but did not reach statistical significance (*p* = *N.S,* **Supplementary Figure 2 K & L**).

At the myofibre level, IHC analysis revealed an average reduction in fibre type-specific cross-sectional area (fCSA) after atrophy which approached significance when pooling fast and slow fibres (**Figure 1H-K**). However, due to limited fibre number in the cross-sections from fewer samples in specific conditions, we also used the open-source “FibeRtypeR” ^82^ web application to predict fibre type proportions using RNA-seq data, with this analysis suggesting a significantly greater proportion of type II fibres at baseline (60% type II vs. 40% type I, *p* = 0.01), atrophy (67% type II vs. 33% type I, *p* < 0.001) and repeated atrophy (63% type II vs. 37% type I, *p* = 0.003) with no significant difference after recovery (61% type II vs. 39% type I, *p* = 0.05) (**Figure 1L**). Overall, demonstrating a reduction in the number type I fibres and an increase in type-II fibres with both atrophy and repeated atrophy and a return to baseline proportions with recovery (**Figure 1L**). From these gene lists, we next interrogated all essential and regulatory fast (*MYL1, MYLPF*), slow (*MYL2, MYL3, MYL6B*) and fetal myosin light chains (*MYL6*) as well as adult (*MYH7, MYH2, MYH1*) and developmental (*MYH3*, *MYH8*) myosin heavy chains (**Figure 1M & N)**. On average, all 6 myosin light chains decreased after atrophy, were fully restored after recovery and then reduced again after repeated atrophy. The slow *MYL2* and *MYL6B* genes were significantly downregulated after atrophy (**Figure 1M)**. A similar temporal profile was observed for slow(er) adult *MYH7* (type I) and *MYH2* (type IIA) genes that both decreased after atrophy and repeated atrophy whereas fast *MYH1* (type IIX) increased after both periods of atrophy (**Figure 1N**). Overall, suggesting a predominant downregulation of slow MYL and MYHĆs after repeated atrophy and an increase in type IIX myosin, thus suggestive of a shift towards a faster phenotype in response to disuse. Finally, expression of embryonic *MYH3* and perinatal *MYH8* both significantly increased after repeated atrophy (**Figure 1N**). Having established coordinated changes in muscle size, muscle strength, muscle quality, myofibre structural characteristics, fibre-type composition, and MYH and MYL gene expression across atrophy, recovery, and repeated atrophy in young adult humans, we next sought to determine whether similar or divergent patterns emerge in aged muscle. However, because repeated atrophy in elderly humans is associated with frailty, reduced balance, and increased fall risk, it is not ethically acceptable to expose older adults to sequential periods of disuse. Therefore, aged animals were used as a surrogate model to examine how ageing influences recovery capacity and susceptibility to repeated atrophy.

### Muscle mass, myofibre structure, fibre-type and myosin isoform expression responses to atrophy, recovery, and repeated atrophy in aged rats

To determine how ageing alters susceptibility to atrophy and repeated atrophy, and whether it impairs the recovery of muscle mass, myofibre structural characteristics, fibre-type composition, and myosin isoform expression, we evaluated muscle weights, histological fibre cross-sectional area and fibre-type distribution, and MYH and MYL gene expression across atrophy, recovery, and repeated atrophy conditions in aged rats. In aged rats, there was no significant reduction in body weight in controls after atrophy or recovery (*p* = *N.S*) (**Supplementary Table 1**). A significant reduction in body weight after repeated atrophy (−18.1%, *p* = 0.02, **Supplementary Table 1**). Therefore, muscle weights were normalised to respective body weights for each condition throughout. TTX administration to the common peroneal nerve (CPN) (**Figure 2A**) resulted in an average reduction of TA muscle weight during initial atrophy (−4.4%, *p* = 0.6) that was not significant, and a significant loss after repeated TTX-induced atrophy (−29.8%, *p* = 0.002, **Figure 2B**). There was also a significant reduction in EDL muscle weight following repeated atrophy (−31.1%, *p* = 0.01). Our previous study demonstrated that young adult rats recovered muscle mass and fibre CSA following TTX-induced atrophy ^66^. Below, we present a physiological and transcriptomic comparison between aged rats and these previously published young rat data to enable robust age-related insights while adhering to ethical refinement principles. Despite TTX-cessation and a return to normal dorsiflexion in aged rats during the recovery period, TA muscle loss continued to occur after recovery (−17.9%, *p* = 0.01, **Figure 2B**), despite shorter TTX exposure and longer active recovery than young adult rats ^66^. There was no significant reduction in plantarflexor (soleus, plantaris and gastrocnemius) muscle weights, thus emphasising precise selective silencing of the dorsiflexor muscles using TTX to the common peroneal nerve (*p = N.S*, **Supplementary Figure 3A-D**). At the transcript level, there was increased expression of developmental myosin heavy chain (*Myh3/8)* genes across all conditions (**Figure 2C**), whereas adult fast (*Myh1/4*), slow (*Myh7*) myosin heavy chains, as well as essential (*Myl3*) and regulatory (*Mylpf/2*) myosin light chains significantly decreased after both periods of disuse (**Figure 2D**). Interestingly, these returned to control levels upon recovery despite continued muscle loss during TTX cessation (**Figures 2C & D**) suggesting that *Myh1/4/7* and *Myl2/3/pf* genes were sensitive to the TTX-disuse stimulus. Conversely, *Myh2*, *Myl1* and *Myl4* gene expression significantly increased following TTX cessation only. Overall, persistent elevation of developmental *Myh3* and *Myh8* mRNA expression coupled with significant reductions in adult fast and slow *Myh* and *Myl* isoforms occurred after both periods of atrophy. Coinciding with reductions in muscle weight and *Myh* and *Myl* expression, IHC analysis (**Figure 2E**) showed a significant loss in average (type I & II combined, **Figures 2F & K**), type I (**Figures 2G & L**) and type IIA (**Figures 2H & M**) myofibre cross-sectional area (fCSA) after repeated atrophy. There was also a significant reduction in pure type I (**Figure 2G & L**) and hybrid I/IIA (**Figure 2I & N**) fCSA after initial TTX-induced atrophy, thus supporting the transcriptional expression data that slow fibres were sensitive to TTX-induced disuse. When relativising left TTX to right control limb, there were no significant changes in fCSA across all fibre types (**Supplementary Figures 3 F-J**). Finally, there was a greater contribution of type IIA and reduced proportion of IIX/IIB fibres after recovery (**Figure 2P**), where fCSA distributions were significantly left-shifted in the left limb, indicating a higher frequency of smaller fibres following TTX-exposure for each fibre type (*p* < 0.0001) (**Figures 2Q-U**).

Having established that aged muscle exhibits impaired recovery capacity, persistent myofibre atrophy, marked reductions in slow and fast myosin isoform expression, and altered fibre-type composition across atrophy, recovery, and repeated atrophy, we also observed that the majority of MYH and MYL genes returned towards control levels during the recovery period, despite continued loss of muscle mass. This dissociation between physiological recovery and transcriptional normalisation suggested that aged muscle retains sensitivity to TTX-induced disuse but is unable to translate transcriptional recovery into structural or functional restoration. To define the molecular basis of this divergent physiological–transcriptional relationship, and to establish a reference molecular profile for subsequent age comparisons, we next examined genome-wide transcriptomic responses to atrophy, recovery, and repeated atrophy in young adult human skeletal muscle, enabling direct comparison with the aged-rat transcriptome profiles that follow.

### Transcriptomic responses to atrophy, recovery, and repeated atrophy in young adult human skeletal muscle

To determine whether the physiological and structural responses observed in young adults are underpinned by coordinated alterations in gene expression, and to establish a molecular reference for comparison with the aged-rat transcriptome, we next performed genome-wide RNA-seq analysis in young human skeletal muscle across atrophy, recovery, and repeated atrophy. In young adult humans, the largest changes in gene expression occurred after initial atrophy, where most genes were downregulated after both atrophy and repeated atrophy (**Figures 3A-F**). PCA analysis revealed a larger overlap between baseline and recovery as well as between atrophy and repeated atrophy timepoints, suggesting a closer transcriptomic profile between both periods of physical activity (baseline and recovery) and inactivity (atrophy and repeated atrophy) (**Figure 3B**). Among the 15,087 genes detected, there were 4-5-fold more DEGs after initial atrophy (381 DEGs) than after repeated atrophy (103 DEGs vs. baseline, 82 DEGs vs. recovery) with predominant downregulation of genes after both atrophy (63%) and repeated atrophy (77% vs. baseline, 85% vs. recovery) (**Figures 3C-F**). Following recovery from initial atrophy, more genes were upregulated (58%) (**Figure 3E**). Venn diagram analysis demonstrated a large proportion of overlapping downregulated (45% of total repeated atrophy DEGs vs. baseline and recovery) or upregulated (53%) genes after both periods of atrophy (**Figure 3G**). During recovery, there were no detected DEGs relative to baseline, with 536 DEGs following recovery from initial atrophy which were all unique from atrophy and repeated atrophy comparisons (**Figures 3E & G**).

**Figure 3.**
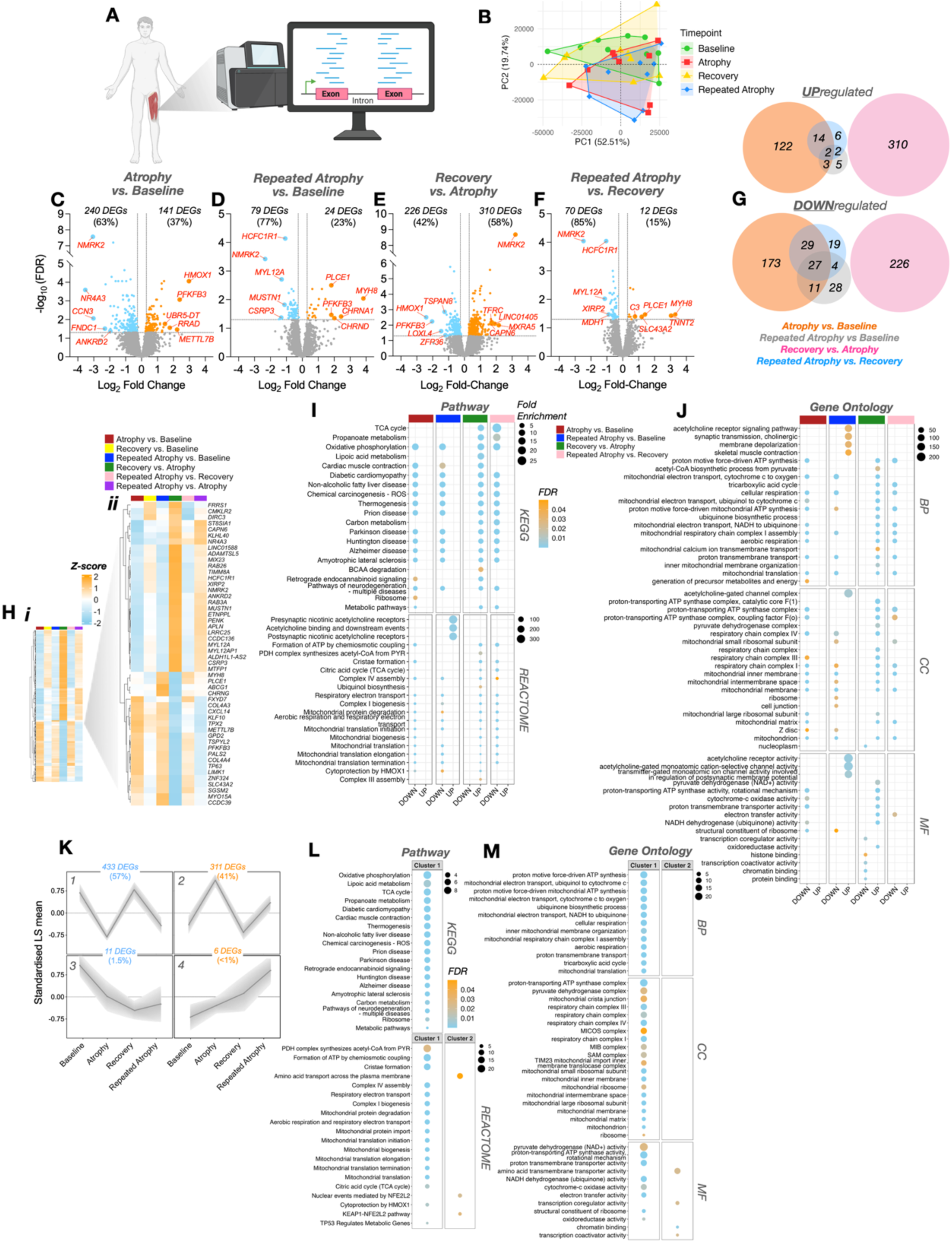
(**A**) Schematic depicting the transcriptomic response to repeated disuse atrophy in young adult human skeletal muscle (created with BioRender.com). (**B**) Principal component analysis (PCA) by time point. (**C-F**) Volcano plots depicting the number of differentially expressed genes (DEGs) that were either DOWN- (blue) or UP-regulated (orange) for each pairwise comparison. No genes met the FDR ≤ 0.05 and ±1.2-fold change thresholds for the repeated atrophy vs. atrophy or recovery vs baseline comparison. Twelve and 10 genes met an unadjusted p value ≤ 0.001 for these comparisons respectively. Labelled genes within volcano plots represent the top 5 DEGs according to greatest FC in either direction of change (FDR ≤ 0.05). (**G**) Venn diagrams depicting the number of unique and common DEGs across all timepoints for both DOWN- (bottom) and UP-regulated (top) DEGs. (**H**) Heatmap of all DEGs in at least one comparison (**i**). Magnified heatmap displaying the top 25 DOWN- (blue) or UP-regulated (orange) genes according to average fold-change across all timepoints (**ii**). Over-representation analysis (ORA) of the top 20 enriched (**J**) pathway (KEGG, top 4 panels; REACTOME, bottom 4 panels) and (**K**) gene ontology (GO) terms for both DOWN- and UP-regulated DEGs across all comparisons (“Atrophy vs. Baseline”, “Repeated Atrophy vs. Baseline”, “Recovery vs. Atrophy”, “Repeated Atrophy vs. Recovery”). (**K**) Self-Organizing Maps (SOM) composed of 761 genes that were significantly differentially expressed in at least one comparison, illustrating gene cluster temporal profiles across the time-course of atrophy, recovery and repeated atrophy. Cluster 1 & 3 (blue) represent genes that were DOWN-regulated following the first period of atrophy. Cluster 2 & 4 (orange) depict UP-regulated genes after initial atrophy. (**L**) Pathway and (**M**) GO analysis revealed enriched terms for clusters 1 and 2 only, in which most genes were either DOWN- (Cluster 1) or UP-regulated (Cluster 2) after atrophy, recovered upon reloading and were less susceptible to changes after repeated atrophy. Cluster 3 represents downregulated genes after atrophy, with retained reduction during recovery and repeated atrophy. Cluster 4 represents upregulated genes after atrophy, retained during recovery, followed by greater increases in expression after repeated atrophy. BP = biological process, CC = cellular component, MF = molecular function.

Interrogation of the most differentially expressed genes across comparisons revealed that *NMRK2,* an NAD^+^ biosynthesis gene, was among the top-most downregulated DEGs after atrophy (−8.4 FC, FDR most significant), ranked second to the nuclear receptor, *NR4A3* (−11.5 FC, **Figures 3C & H**). *NMRK2* was also the most downregulated gene after repeated atrophy versus baseline (−5.0 FC) and when compared with recovery (−5.6 FC), as well as the most upregulated DEG after recovery versus atrophy (9.2 FC, **Figures 3C & H).** Interestingly, after repeated atrophy only, the most upregulated genes were developmental myosin heavy chain *MYH8* (14.2 FC vs. baseline, 9.8 FC vs. recovery), tropomyosin-binding subunit *TNNT2* (8.3 FC vs. recovery) and NMJ nicotinic acetylcholine receptor (nAChR) subunit genes, *CHRNA1* (5.4 FC versus baseline) and *CHRND* (4.0 FC versus baseline) (**Figure 3D & F)**. Another AchR subunit gene, *CHRNG* (2.86 FC versus baseline), was one of the most upregulated by average FC (across all timepoints) only after repeated atrophy (**Figure 3H**).

Such transcriptional perturbations across the transcriptome corresponded to significant downregulation of enriched pathway and gene ontology terms related to aerobic metabolism, respiration (e.g., NAD^+^ / NADH metabolism, TCA cycle, ETC, ATP synthesis) and mitochondrial structure, function and turnover after both periods of disuse, with these genes returning to baseline levels after recovery (**Figures 3I & J** and **Figure 3K, cluster 1).** Furthermore, enrichment of upregulated genes (including AchR subunits and *MYH8 and TNNT2* highlighted above) were associated with muscle-specific terms including acetylcholine receptor signalling, synaptic transmission, membrane depolarisation and skeletal muscle contraction (**Figure 3I and J**).

### Temporal profiling of transcriptional responses (SOM analysis) in young adult human skeletal muscle

To build upon the global transcriptional alterations identified above, and to determine whether repeated disuse modifies the temporal coordination of gene expression rather than merely its magnitude, we next applied self-organising map (SOM) profiling to cluster transcripts according to their trajectories across atrophy, recovery, and repeated atrophy. This approach enabled us to distinguish genes that exhibited transcriptional protection during repeated disuse from those that retained or amplified their responses after atrophy into recovery and across successive atrophic episodes. Temporal gene clustering analysis (SOM) revealed most genes decreased (57%, cluster 1) and increased (41%, cluster 2) after atrophy, returned to baseline levels after recovery, and were less susceptible to changes after repeated disuse (**Figure 3K**). This represents a transcriptional attenuation, resulting in a smaller magnitude of change to repeated atrophy stimuli for both downregulated (cluster 1) and upregulated (cluster 2) genes. Based on SOM standardised means, there was a 40% smaller magnitude in downregulation of genes in cluster 1 and an 72% smaller magnitude in upregulation of genes in cluster 2 after repeated atrophy compared with initial atrophy. Enriched pathway and GO terms associated with the response of these downregulated genes to both periods of disuse (cluster 1) were the same aerobic metabolism, respiration and mitochondrial-related terms identified across global downregulated genes described above (i.e. NAD^+^ / NADH dehydrogenase activity, TCA cycle, ETC, ATP synthesis and mitochondrial structure, assembly and protein turnover pathway genes, **Figures 3L & M**). Genes that were upregulated after atrophy and increased less after repeated atrophy (cluster 2, **Figure 3K**), were enriched in terms related to amino acid transport, transcription coregulator/coactivator activity and chromatin binding (**Figure 3L & M**). In contrast, genes associated with these terms/pathways were also downregulated during recovery from atrophy (**Figure 3J**). Therefore, the fewer number of total DEGs coupled with less susceptible alterations to repeated versus initial atrophy suggests a transcriptionally protective memory of earlier atrophy. SOM cluster temporal analysis also identified 11 downregulated genes (*NR4A1, NR4A3, PRKAG2, RP9, CIITA, CORO2B, ENSG00000242299, RPL35, SNRPA1, UBE3D, RPL29P11*) after atrophy that remained downregulated during recovery, despite restoration of muscle mass (cluster 3, **Figure 3K**). This included nuclear receptor genes, *NR4A1* and *NR4A3,* as well as *PRKAG2* (AMPK gamma2), where *NR4A3* was among the top ranked downregulated genes within enriched metabolic processes. Ribosomal genes (*RPL35*, *RPL29P11*) also exhibited downregulated enrichment at the pathway level after atrophy (**Figure 3J**), indicating a transcriptional ‘memory’ of the initial disuse episode through the retention of suppressed expression even after the recovery period. Finally, 6 genes (*TNNT2, MYH3, DCLK1, ZBED6CL, CDC73, WDR44*) were upregulated after atrophy, remained elevated during recovery, with even larger increases in expression after repeated atrophy (cluster 4, **Figure 3K**). This included *TNNT2* (top ranked upregulated gene as discussed above), embryonic myosin heavy chain (*MYH3*), *DLK1* associated with regeneration of muscle following injury, and *ZBED6CL* also linked with muscle development. Therefore, whilst most genes demonstrated transcriptional protection after repeated atrophy, a small number of genes related to muscle development and regeneration (together with the AchR subunit genes discussed above) demonstrated amplified responses to repeated atrophy in adult human muscle. The retained upregulation and larger increases after repeated disuse also suggests a memory of earlier atrophy at the transcriptional level in these genes.

Taken together, the global DEG analysis and SOM temporal profiling demonstrate that young adult skeletal muscle exhibits a coordinated transcriptional protection against repeated disuse, characterised by attenuated suppression of NAD^+^ and oxidative metabolism, mitochondrial, and structural gene networks, with only a small subset of developmental and neuromuscular genes retaining or amplifying their responses across successive atrophic episodes. To determine whether ageing disrupts this protective molecular programme, and to identify transcriptional signatures underlying the impaired physiological recovery observed in aged muscle, we next examined genome-wide RNA-seq profiles in muscle from aged rats across atrophy, recovery, and repeated atrophy.

### Transcriptomic responses to atrophy, recovery, and repeated atrophy in aged rat skeletal muscle

Together, the differential gene expression analysis and SOM profiling in young adult muscle revealed a coordinated transcriptional protection to repeated disuse, characterised by attenuated suppression of NAD^+^ and oxidative metabolism, mitochondrial, and structural gene networks. To determine whether ageing alters this temporal and pathway-level resilience, and to identify molecular signatures underpinning the impaired physiological recovery observed in aged muscle, we next performed genome-wide RNA-seq analysis in muscle from aged rats across atrophy, recovery, and repeated atrophy. Alongside the loss of TA muscle mass during atrophy, recovery and repeated atrophy in aged animals, there were considerably large transcriptomic perturbations after atrophy and recovery with the greatest changes detected after repeated atrophy (**Figures 4A-I**). PCA analysis showed similar clustering of TTX-induced atrophy and repeated atrophy, and distinct clustering after TTX cessation-induced recovery (**Figures 4B & C**). There was also considerable overlap of the sham surgical and pooled non-surgical controls (**Figure 4B**). Given there were also no DEGs detected between sham and pooled contralateral control limb samples, suggesting the surgery had no significant impact on gene transcription, we excluded sham samples from further analysis and pooled contralateral (right) non-surgical limb samples for the control group. Among the 11,284 genes detected post-normalization, there was a similar number of DEGs in response to atrophy (2,106 DEGs) and recovery (2,008 DEGs) with 2-fold more DEGs detected after repeated atrophy (3,937 DEGs) vs. controls. The proportion of downregulated DEGs was 56% after atrophy (**Figure 4D**), 53% after recovery and 56% (vs. control) or 58% (vs. recovery) after repeated atrophy (**Figure 4F-H)**. However, despite a similar number of total DEGs after atrophy and recovery, the majority of downregulated (90%) and upregulated (74%) DEGs were unique after TTX cessation (**Figure 4J**), as evidenced by the distinct PCA clustering of recovery samples (**Figure 4C**). Also, whilst the 2-fold greater number of DEGs after repeated atrophy versus initial atrophy may be partially explained by the lack of recovery during the no TTX period, when relativised to previous recovery, the number of DEGs after repeated atrophy (3,198 DEGs) was still 1.5-fold more than initial atrophy (2,052 DEGs) (**Figure 4H**), that corresponded to a -11.9% reduction in muscle mass after repeated atrophy versus recovery (**Figure 2B**). There were 79 DEGs after repeated versus initial atrophy in aged animals with a larger number of downregulated (84%) versus upregulated (16%) genes (**Figure 4I)**. Most of these were shared across both periods of atrophy (2,461 DEGs/75% downregulated, 3,935 DEGs/61% upregulated), but several downregulated (1,294 DEGs/25%) and upregulated (1,550/39%) DEGs that were unique to repeated atrophy (**Figure 4J**).

**Figure 4.**
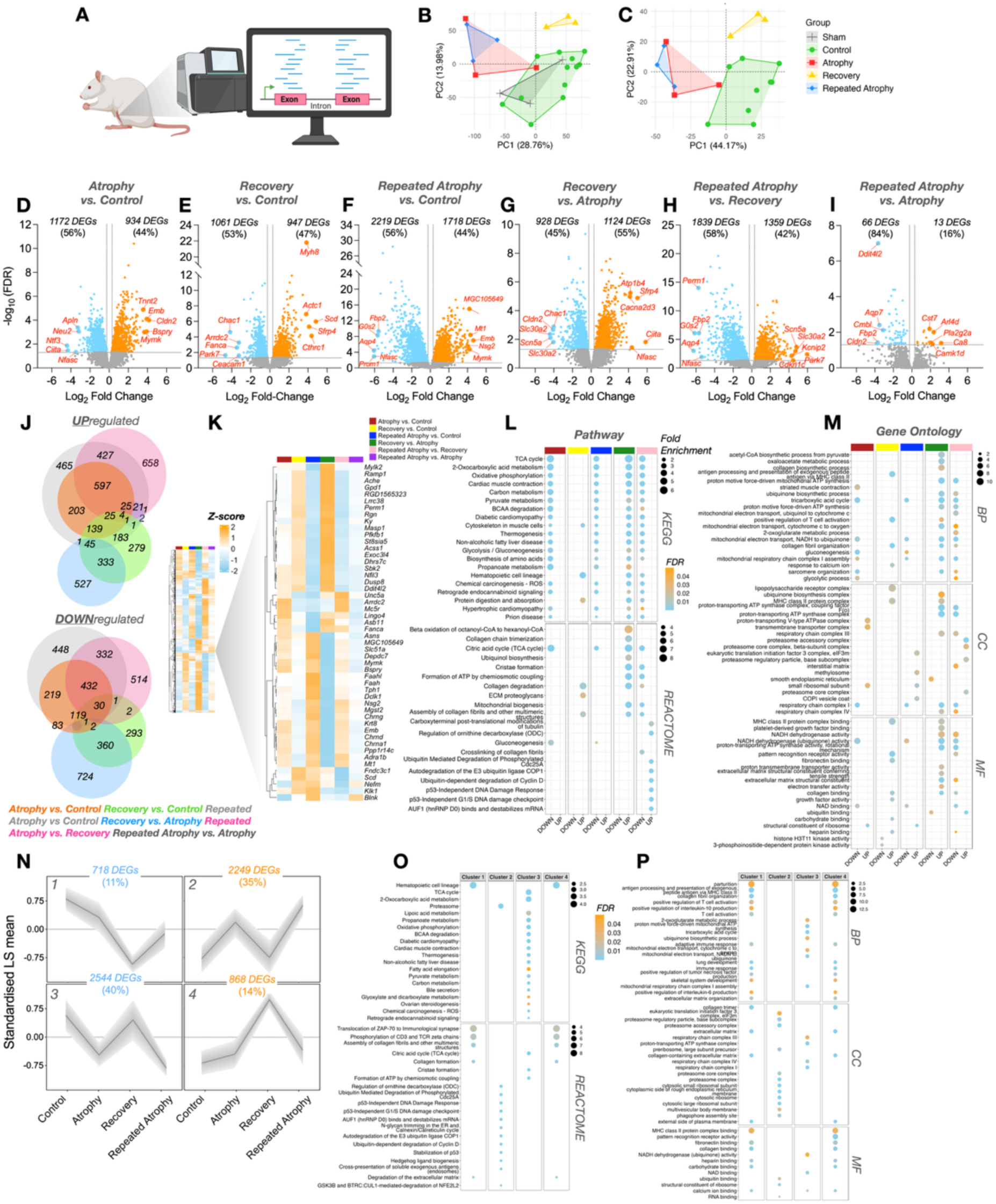
(**A**) Schematic depicting transcriptomic (via RNA-seq) analysis of repeated TTX-induced disuse atrophy in aged animals (created using BioRender.com). Principal component analysis (PCA) including (**B**) and excluding (**C**) sham surgery group. (**D-I**) Volcano plots depicting the number of differentially expressed genes (DEGs) that were either DOWN-(blue) or UP-regulated (orange) for each pairwise comparison. Labelled genes within volcano plots represent the top 5 DEGs according to greatest fold-change with FDR ≤ 0.05. There were no DEGs detected in sham versus pooled contralateral control limb, suggesting the surgery had no significant impact on gene transcription. The sham group was therefore excluded for all downstream analyses and pooled contralateral limb was used as the control. (**J**) Heatmap of all DEGs in at least one comparison (***i***). Magnified heatmap displaying the top 50 DOWN-(blue) and top 50 UP-regulated (orange) genes (by gene symbol) by average fold-change across all pairwise comparisons (***ii***). (**K**) Venn diagrams depicting the number of unique and common DEGs across all conditions for both UP- (top) and DOWN-regulated (bottom) DEGs. Over-representation analysis (ORA) of the top 20 enriched (**L**) pathway (KEGG, top 4 panels; REACTOME, bottom 4 panels) and (**M**) gene ontology (GO) terms for both DOWN- and UP-regulated DEGs across all comparisons (“Atrophy vs. Control”, ”Recovery vs. Control”, “Repeated Atrophy vs. Control”, “Recovery vs. Atrophy”, “Repeated Atrophy vs. Recovery”). (**N**) Self-Organizing Maps (SOM) composed of a total 6379 genes that were DEGs in at least one comparison, illustrating gene cluster temporal profiles across the time-course of repeated disuse atrophy. Profiles 1 & 3 represent genes that were DOWN-regulated following the first period of atrophy whereas profiles 2 & 4 depict UP-regulated genes after initial atrophy. (**L**) Pathway and (**M**) GO analysis revealed enriched terms across all cluster profiles where most genes were either UP- (cluster 2) or DOWN-regulated (cluster 3) after atrophy, recovered upon TTX cessation, followed by larger changes (in the same direction of initial atrophy) after repeated disuse atrophy. BP = biological process, CC = cellular component, MF = molecular function.

Regarding gene-specific alterations, the top downregulated DEGs included genes associated with NMJ function and stability (*Ache, Unc5a, Ntf3, Nfasc*), mitochondria and metabolism (*Park7, Perm1, Nfil3, Gpd1, Acss1, Mc5r, Dhrs7c, G0s2, Fbp2*), muscle structure and contraction (*Lrrc38, Ky, Mylk2, Rgn*) as well as DNA damage (*Fanca, Ddit4l2*) (**Figure 4K).** Interestingly, the myokine, *Apln*, important for angiogenesis, muscle regeneration and ATP synthesis was among the top 5 downregulated genes after initial TTX-induced atrophy (−9.7 FC, **Figure 4D**) (that also decreased after repeated atrophy) and was within the top 25 downregulated genes in young adult humans after atrophy (**Figure 3H**). Concerning the top upregulated genes, the muscle specific fuso-gene, *Mymk* significantly increased after both atrophy (12.8 FC, **Figure 4D**) and repeated atrophy (18.6 FC, **Figure 4F**), implying satellite cell fusion in attempt to repair and/or offset TTX-induced muscle wasting. Also, among the top-ranked upregulated genes after initial atrophy was tropomyosin-binding, *Tnnt2* (11.8 FC, **Figure 4D)**. Embryonic *Myh8* was one of the most upregulated DEGs after recovery (14.3 FC, **Figure 4E**) and was among the top-ranked upregulated DEGs in young adult human muscle after repeated atrophy (**Figures 3D & F**). Further, NMJ AChR subunit genes, *Chrnd,* and *Chrna1* were both ranked among the top upregulated genes in aged animal muscle after both atrophy (*Chrnd* = 5.8 FC*, Chrna1* = 7 FC) and to a greater extent repeated atrophy (*Chrnd* = 12.5 FC*; Chrna1* = 16 FC) whilst *Chrng* significantly increased only after repeated atrophy (**Figure 4K**). Such AChR genes were also similarly upregulated in young adult human muscle following disuse (**Figure 3D & H**). Collectively, genes *Tnnt2, Myh8, Chrna1, Chrnd* and *Chrng*, increased in response to disuse-induced muscle wasting, irrespective of age, species and atrophy model utilised.

Enrichment analysis of downregulated genes revealed similar pathways and GO terms after both atrophy and repeated atrophy in aged muscle. Specifically, terms related to metabolic processes and aerobic respiration (e.g., NAD binding / NADH dehydrogenase activity, TCA cycle, ETC and ATP synthesis) as well as muscle-specific terms (e.g., muscle contraction, cytoskeleton in muscle cells and sarcomere organisation) (**Figure 4L & M**). Such terms were also enriched in downregulated DEGs after atrophy and repeated atrophy in young adult human muscle. In contrast, most of these terms were enriched in upregulated DEGs during recovery from atrophy, suggesting they recovered after the initial disuse, despite continued muscle wasting after TTX cessation. This suggests that genes were TTX-sensitive during disuse rather than simply reflecting the continued muscle wasting observed during recovery and a possible attempt to restore muscle during return to activity. This was also observed in young adult human muscle, where expression of genes within these pathways, alongside skeletal muscle size and strength, returned to baseline levels during recovery. Upregulated pathway enrichment that occurred after both atrophy and repeated atrophy was evident within ribosomal structure and subunit genes (**Figure 4L**). Interestingly, genes associated with proteasomal core and accessory complex, ECM turnover, ubiquitin-mediated proteolysis and DNA damage pathways only increased after repeated atrophy (**Figure 4L-M**). While DEG-level analyses in aged muscle revealed extensive perturbations, including exaggerated suppression of oxidative metabolism, NAD⁺/NADH, and structural gene networks after repeated atrophy, these comparisons do not capture how transcriptional responses evolve over time relative to the initial atrophic episode. Given that SOM profiling in young adult muscle identified clear temporal signatures of transcriptional protection after repeated disuse, we next applied the same SOM approach to aged rat muscle to determine whether ageing disrupts this pattern of coordinated temporal regulation. This analysis enabled us to identify gene clusters that define the impaired recovery and heightened susceptibility of aged muscle across atrophy, recovery, and repeated atrophy.

### Temporal profiling of transcriptional responses (SOM analysis) in aged rat skeletal muscle

To determine whether the temporal protection observed in young adult muscle is lost with ageing, and to establish how gene-expression trajectories relate to the impaired physiological recovery in aged rats, we next applied SOM temporal clustering to the aged-rat transcriptome across atrophy, recovery, and repeated atrophy. Temporal clustering analysis by SOM profiling revealed that 75% of genes (40% downregulated, cluster 3; 35% upregulated, cluster 2) after atrophy, were largely restored to control levels after TTX cessation during recovery, followed by larger magnitude changes after repeated disuse (**Figure 4N**). This indicates that most genes recovered after initial atrophy despite continued muscle loss in aged muscle yet showed exaggerated responses after repeated atrophy. Specifically in cluster 3, downregulated genes after atrophy that returned to control levels during recovery (despite continued loss of muscle mass), followed by greater reductions during repeated atrophy, were enriched in: aerobic metabolism, TCA cycle, NAD binding/NADH dehydrogenase activity, ETC and ATP synthesis in the mitochondria as well as in muscle structure and function (cluster 3) (**Figures 4O & P**). Based on SOM standardised means, on average there was a 95% larger magnitude of decreased expression of genes within cluster 3 after repeated atrophy compared with initial atrophy. Therefore, genes in these metabolic pathways demonstrated, on average, a 95% greater downregulation of transcripts in response to repeated atrophy, whereas young adult human muscle demonstrated a less susceptible, attenuated transcriptional response in genes within the same pathways following repeated atrophy. This suggests a negative, detrimental and exaggerated transcriptional response to repeated atrophy in aged muscle. Importantly, these genes still recovered to control levels during recovery, despite continued muscle wasting in aged animals. Therefore, the amplified response to repeated atrophy in aged muscle was not simply due to the lack of recovery but rather an exaggerated response to repeated atrophy having experienced earlier TTX-induced disuse atrophy. Genes within cluster 2 (**Figure 4N**) that were upregulated after atrophy, returned to control levels during recovery and were upregulated to a greater extent during repeated atrophy, were enriched in proteosome, ECM turnover, and DNA damage pathways in aged muscle. Based on SOM standardised means, there was an average 141% larger magnitude in upregulation of genes in cluster 2 after repeated atrophy compared to the initial atrophy. These genes/terms were unique to aged muscle, also demonstrating recovery of gene expression despite continued muscle wasting during the recovery period, followed by exaggerated transcriptional and physiological responses to repeated atrophy. A small proportion of genes also decreased (11%, cluster 1) or increased (14%, cluster 4) to a lesser extent during atrophy with the largest change occurring during recovery, followed by a return to atrophy levels during repeated atrophy (**Figure 4N**). Genes in both clusters 1 & 4 were enriched in inflammatory pathways, ECM turnover, collagen assembly and matrix organisation (**Figure 4O & P**), suggestive that changes in inflammatory and ECM genes were associated with continued muscle wasting in aged muscle, despite TTX cessation.

Together, these DEG-level and temporal transcriptional analyses demonstrate that aged skeletal muscle exhibits a marked susceptibility to repeated disuse, characterised by greater suppression of oxidative metabolism, mitochondrial, NAD⁺/NADH, and structural gene networks, despite the recovery of these pathways during the TTX cessation period. These findings, coupled with the continued physiological decline observed in aged rats during recovery, suggest that ageing disrupts the capacity to translate transcriptomic restoration into functional or structural recovery. To contextualise this age-related susceptibility within species, and to determine whether these exaggerated molecular responses diverge from those observed in younger muscle exposed to the same disuse paradigm, we next compared the aged-rat findings with previously published young adult rat data generated using the same TTX-induced atrophy and recovery model.

### Physiological and transcriptomic responses to TTX-induced atrophy and recovery in young adult rat skeletal muscle

In both young adult human and aged rat muscle, initial and repeated atrophy were characterised by marked downregulation of genes associated with aerobic respiration and mitochondrial function, with these pathways recovering upon return to habitual activity in both species despite divergent physiological outcomes. Given the full physiological recovery observed in young adults and the persistent muscle loss in aged rats, we next examined previously published data from young adult rats exposed to the same TTX-induced atrophy and recovery model to contextualise the age-related susceptibility observed herein. This within-species comparison enables robust interpretation of age effects while adhering to ethical refinement principles, as it avoids unnecessary duplication of animal experiments. Notably, our earlier work demonstrated that young adult rats recovered muscle mass and fibre cross-sectional area following a slightly longer period of TTX exposure and a shorter active recovery interval than that used in the present aged cohort ^66^. Specifically, young adult rats received 2 wks of TTX exposure to the common peroneal nerve (versus 1 wk in aged rats in the present study), resulting in an average -51% and -69% loss of TA muscle mass and fCSA, respectively (**Figures 5A, B & C**). Interestingly, 1 wk active recovery (versus 1.5 wks in aged animals) recovered TA mass and fCSA by 53 and 63%, respectively compared to the prior 2-week atrophy period (**Figures 5A & C**). This physiological response contrasts with aged animals herein, where TA muscle weight and fCSA continued to decrease during a longer active recovery, despite less atrophy observed following shorter TTX exposure (**Figures 5B-C**). Together with similar recovery of genes within the same pathways in young adult human muscle and aged rat muscle, this suggests that the lack of recovery in aged animals is dependent on age and not insufficient time to recover. In support of this, overlapping transcriptome datasets from young and aged animals revealed a considerable degree of overlap of DEGs after atrophy (77% upregulated and 72% downregulated DEGs) with a much smaller overlap after recovery (34% upregulated and 26% downregulated) (**Figures 5D-H**).

**Figure 5.**
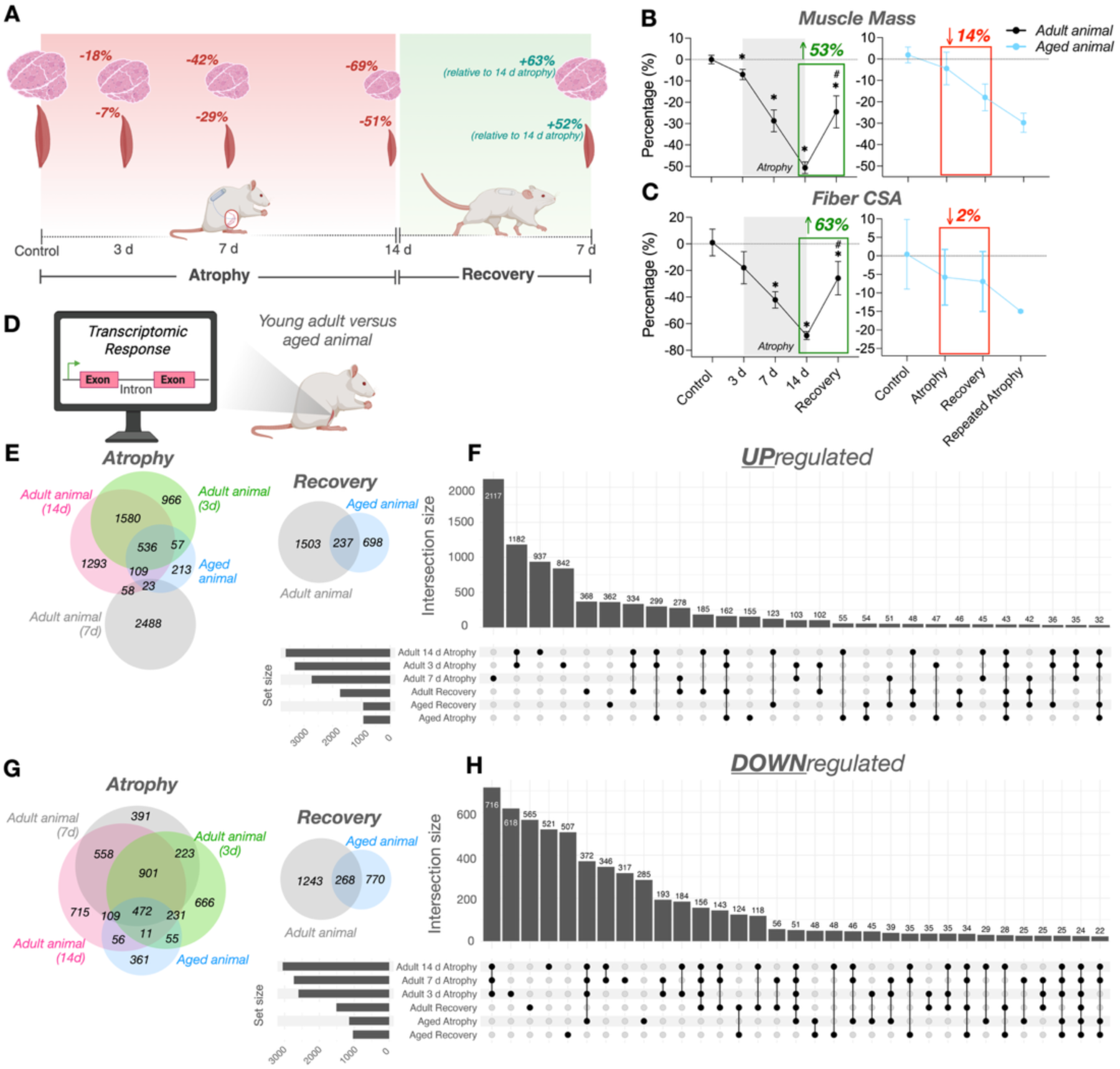
(**A**) Schematic of study design (created using BioRender.com) of TTX-induced atrophy (3, 7 and 14 days/d) and subsequent recovery (7 d) in young adult rats (Fisher et al.,) ^66^. There was a loss of TA muscle mass (**B**) and fCSA (**C**) at 3, 7 and 14 d atrophy (−7 to -18%, -29 to -42% and -51 to -69%, respectively). In contrast to aged animals, muscle mass and fCSA was restored by 53 and 63%, respectively following 7 d recovery in young adult animals (**B & C** redrawn from Fisher et al., ^66^). (**D**) Overlap of transcriptome in young adult and aged rat muscle revealed many common UP- (**E & F**) and DOWN-regulated (**G & H)** DEGs after atrophy and a larger proportion of unique genes after recovery.

### Integrative methylome-transcriptome analysis identifies epigenetic regulation of metabolic, mitochondrial, and neuromuscular gene networks after repeated atrophy

To determine whether the transcriptional signatures identified above were accompanied by coordinated epigenetic alterations, we next integrated RNA-seq and RRBS datasets by mapping differentially expressed genes and differentially methylated regions onto shared temporal trajectories using self-organising map (SOM) profiling. This approach enabled us to identify genes that exhibited inverse SOM profiles of DNA methylation and gene expression across atrophy, recovery, and repeated atrophy, thereby defining the differentially methylated, differentially expressed genes (DmDEGs) regulated by repeated disuse. Repeated periods of disuse atrophy and recovery led to differentially methylated DNA regions (DMRs) that were inversely associated with the DEGs identified in both young (**Figures 6A-D**) and aged (**Figures 6E-H**) skeletal muscle. These genes are termed **D**ifferentially **m**ethylated **DEGs** (DmDEGs). In young adult humans, 71 DMRs on 57 genes (from cluster 1 above, **Figure 3K**) were identified as DmDEGs with an inverse relationship of hypermethylation and decreased expression. These DmDEGs were hypermethylated and downregulated after atrophy and repeated atrophy, with both DNA methylation and gene expression returning to baseline levels after recovery where skeletal muscle size was restored (cluster 1, **Figures 6B*i* & C*i***).

**Figure 6.**
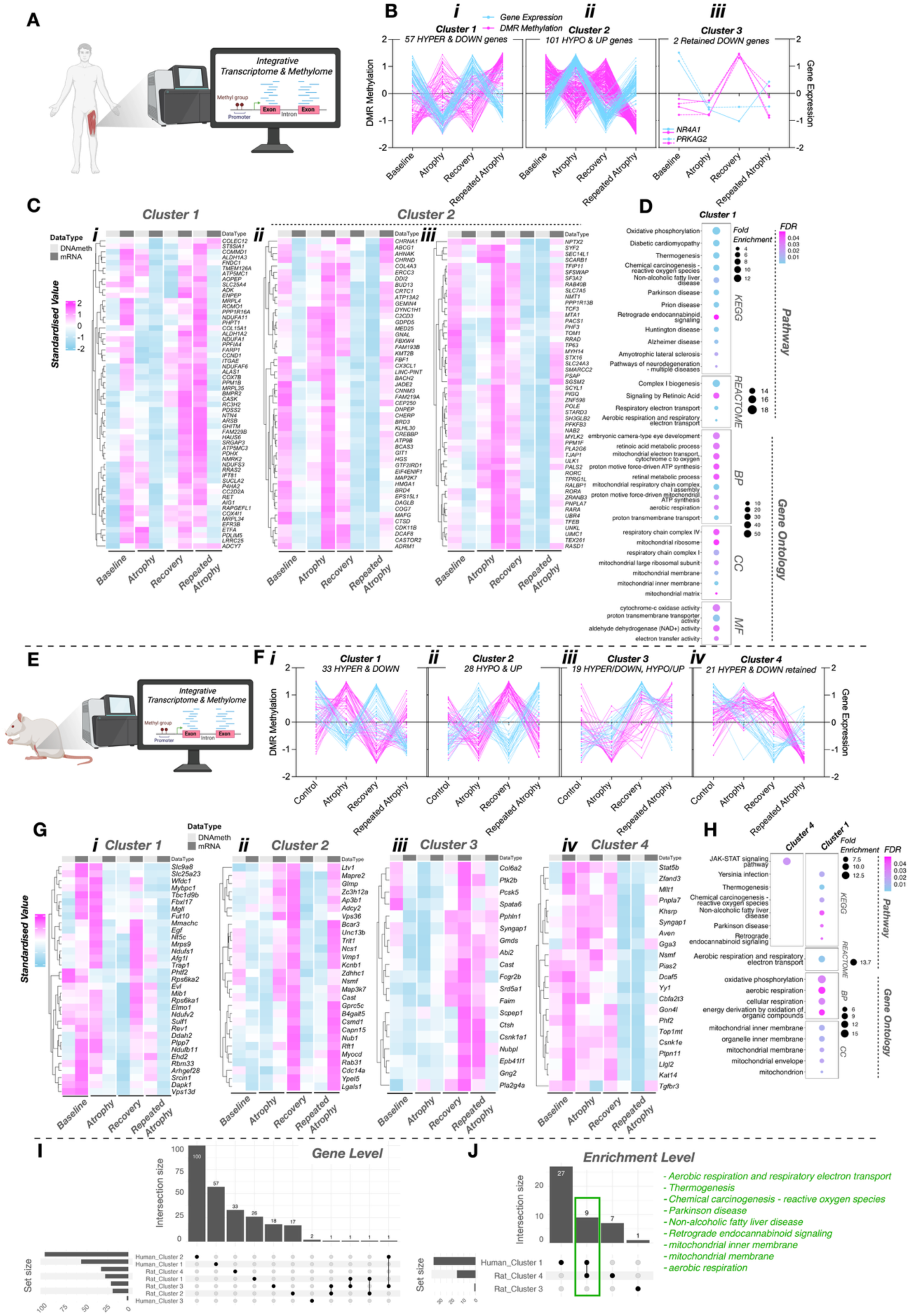
Integrative DNA methylome and transcriptome analyses in young adult human (**A-D**) identified 57 hypermethylated and downregulated genes after both periods of disuse that were returned to baseline levels after recovery (cluster 1, **Bi & Ci**); 101 hypomethylated and upregulated genes after both periods of disuse that also returned to baseline levels after recovery (cluster 2, **Bii & Cii**); 2 downregulated/hypermethylated genes after initial atrophy that were retained during recovery (cluster 3, **Biii & Ciii**). There were significant pathway and GO terms for cluster 1 in young human muscle of hypermethylated/downregulated genes after atrophy, restored during recovery, and hypermethylated/downregulated after repeated atrophy enriched in aerobic metabolism/respiration, NAD^+^ activity, ETC, ATP synthesis, mitochondrial and ribosome terms (**D**). Integrative analyses in aged rat muscle (**E-G**) revealed 33 genes (**Fi & Gi,** cluster 1) hypermethylated/downregulated after atrophy, hypomethylated/upregulated upon TTX cessation during recovery, and hypermethylated/downregulated after repeated atrophy; 28 unique genes (**Fii & Gii,** cluster 2) were hypomethylated/upregulated after atrophy, hypermethylated/restored after recovery, and hypomethylated/upregulated again after repeated atrophy (**Fiii & Giii,** cluster 2); 19 unique genes (cluster 3) that were hypermethylated/downregulated after atrophy, hypermethylated yet upregulated during recovery, and hypomethylated/upregulated after repeated atrophy; 21 unique genes (**Fiv and Giv,** cluster 4) were hypermethylated/downregulated after atrophy, with retained hypermethylation and a continued decrease in gene expression during recovery followed by subsequent hypomethylation and increased gene expression after repeated atrophy. Significant enriched pathway and GO terms (**H**) for cluster 1 were related to aerobic metabolism/respiration, electron transport and specifically mitochondrial structure (for cluster 1) and JAK-STAT enrichment (in cluster 4). Average DMR methylation is depicted in aged animal heatmaps whenever there is >1 DMR for a given gene (**G**). Average mRNA expression is also presented where there were multiple transcript variants (**G**). UpSet plot analysis of young adult human and aged rat muscle gene clusters revealed only a few epigenetically regulated genes that overlapped between both models (**I**). However, there were 9 overlapping enriched pathway / GO terms between human and animal muscle (**J**). Note that ‘DMR Methylation’ is standardised percentage DMR methylation derived from SOM profiling analysis and ‘Gene Expression’ is representative of standardised LS means from SOM profiling analysis (**B & F**). BP = biological process, CC = cellular component, MF = molecular function.

ORA analysis identified these DmDEGs as enriched in NAD⁺ metabolism, the electron transport chain (ETC), ATP synthesis, mitochondrial structural pathways, and mitochondrial ribosome components, demonstrating that the pathway terms enriched within cluster 1 of the transcriptomic analysis (downregulated after atrophy, returning to baseline during recovery, and showing reduced susceptibility during repeated atrophy; Figure 3K) were also subject to epigenetic regulation in response to both disuse atrophy and repeated atrophy. Notably, *NMRK2*, a key gene involved in NAD⁺ biosynthesis and one of the most strongly downregulated genes after both periods of disuse in young adult humans, was included among these epigenetically regulated loci. *NMRK2* exhibited hypermethylation and reduced expression during both atrophy periods, followed by hypomethylation and increased transcription towards baseline levels during recovery. In addition to these hypermethylated and downregulated genes, there were 189 DMRs on 101 genes that were hypomethylated and upregulated after both atrophy and repeated atrophy (cluster 2, **Figures 6B*ii* & C*ii & Ciii***). There was no significantly enriched pathway or GO terms for this cluster, thus suggestive that most of these genes likely have divergent molecular, cellular and biological roles. However, several interesting hypomethylated and upregulated genes emerged from this cluster. Specifically, the AChR genes, *CHRND* and *CHRNA1* (that were upregulated after atrophy and/or repeated atrophy across both human and rat muscle transcriptomes, **Figure 3D-H**), demonstrated the largest hypomethylation and increased expression after repeated atrophy (**Figures 6B*ii* and C*ii***). These genes were among the few DmDEGs in young human muscle that demonstrated increased susceptibility and exaggerated responses to repeated atrophy. Finally, *NR4A1* and *PRKAG2* (AMPK gamma2) were downregulated after atrophy despite no detectable change in DNA methylation at that stage, followed by hypermethylation and sustained suppression of transcription during recovery and repeated atrophy (cluster 3, **Figure 6Biii**). This pattern suggests that the retained reduction in expression during recovery is likely driven by the emergence of hypermethylation at these loci, even though skeletal muscle size and function were restored upon returning to habitual activity. The full DNA methylome dataset for repeated atrophy in young adult human muscle is provided in the supplementary material (**Supplementary Figure 4**).

In aged rat muscle, integrative analysis of DNA methylome and transcriptome (**Figure 6E**) revealed 41 DMRs on 33 unique genes were hypermethylated and downregulated after atrophy (cluster 1, **Figure 6Fi & Gi**), hypomethylated and upregulated upon TTX cessation during the recovery period, and finally hypermethylated and downregulated after repeated atrophy. As with the young human muscle, these genes were associated with aerobic metabolism/respiration, electron transport and mitochondrial components (**Figure 6H**). Integrative analysis also revealed 40 DMRs on 28 unique genes (cluster 2, **Figure 6Fii and 6Gii**) that were hypomethylated and upregulated after atrophy, hypermethylated after recovery, and hypomethylated and upregulated again after repeated atrophy. ORA analysis revealed no significantly enriched pathways or GO terms for this cluster. There were also 26 DMRs on 19 unique genes (cluster 3, **Figure 6Fiii & Giii**) that were hypermethylated and downregulated after atrophy, hypermethylated yet upregulated during recovery, and hypomethylated and upregulated after repeated atrophy, with no significantly enriched pathways or GO terms. Interestingly, 43 DMRs on 21 unique genes (cluster 4, **Figure 6F*iv* *& Giv)*** demonstrated hypermethylation and downregulation after atrophy, with retained hypermethylation and a continued decrease in gene expression during recovery, followed by subsequent hypomethylation and increased gene expression after repeated atrophy. These genes were enriched for JAK–STAT signalling pathways (cluster 4, **Figure 6H**) and exhibited retained epigenetic and transcriptional signatures following the initial atrophy period that persisted into recovery, yet became oppositely hypomethylated and upregulated after repeated atrophy.

Despite the differential physiological response to repeated atrophy and recovery, both young adult humans (cluster 1, **Figures 6B*i*, C*i***, **D)** and aged animals (cluster 1, **Figure 6F*i***, **G*i*, H**) demonstrated associated DNA methylation and gene transcription of genes enriched within the same pathway and GO terms related to aerobic metabolism/respiration, electron transport and mitochondria (**Figures 6D & H**). However, the majority of these DmDEGs were specific to young human or aged rat muscle (**Figure 6I**), despite enrichment of the same metabolic pathways across species and models of atrophy (**Figure 6J**). Further cross-species comparison of the shared transcriptomic and epigenetic responses to repeated disuse in young adult human and age animal muscle via UpSet plots (**Supplementary Figure 5**), identified several common up (**Supplementary Figure 5A**) and downregulated DEGs (**Supplementary Figure 5B**) as well as hypo- (**Supplementary Figure 5C**) and hyper-methylated DMRs (**Supplementary Figure 5D**). It was also apparent that out of the 761 DEGs used for the SOM temporal profiling in human muscle, 54% (414 out of 761) of the genes were also differentially expressed in the aged rat muscle (**Supplementary Figure 5E**). Pathway (**Supplementary Figure 5F**) and GO (**Supplementary Figure 5G**) analyses suggested these shared genes were also related to oxidative metabolism, NAD^+^, TCA cycle, ETC, ATP synthesis and mitochondrial function. Among these shared DEGs, there were 406 DMRs on 189 shared DEGs in young adult human muscle (**Supplementary Figure 5H)** that were associated with oxidative metabolism, ETC, ATP synthesis and mitochondria (**Supplementary Figure 5I & J**). In aged animals, these there were 56 DMRs on 16 shared DEGs (**Supplementary Figure 5K**) that were related to mitochondria and RNA splicing (**Supplementary Figure 5L**). Overall, these data demonstrate epigenetic regulation of shared genes associated with oxidative metabolism across species in response to atrophy, recovery and repeated atrophy. The full DNA methylome data to repeated atrophy in aged rat muscle is also available as supplementary material (**Supplementary Figure 6**).

Together, these integrative analyses demonstrate that repeated atrophy imprints coordinated epigenetic and transcriptional alterations across metabolic, mitochondrial, and neuromuscular pathways, with several DmDEGs showing inverse SOM profiles across atrophy, recovery, and repeated atrophy. Because NAD⁺ metabolism genes featured prominently among these epigenetically regulated clusters, we next examined whether repeated disuse disrupts NAD⁺ biosynthesis and salvage pathways in young adult humans and aged rats.

### Regulation of NAD⁺ biosynthesis pathways following repeated atrophy

Genes involved in NAD⁺/NADH metabolism were strongly represented among the DmDEGs identified through the integrative SOM analysis, showing downregulation and hypermethylation after atrophy and repeated atrophy in both young adult human (**Figure 7A-D)** and aged rat muscle (**Figure 7E-I**). Moreover, the NAD^+^ biosynthetic gene, *NMRK2,* was the among the top downregulated genes after both periods of atrophy with inversely associated DNA across the time course of atrophy, recovery and repeated atrophy. Genes enriched in these pathways were also restored during the recovery period. Targeted mRNA analysis (via qPCR) of *Nmrk2* in aged rat muscle revealed a non-significant reduction in *Nmrk2* expression after atrophy, followed by a significant increase during recovery and a larger significant reduction after repeated atrophy (**Figure 7E & G**). Given the predominant role of *NMRK2/Nmrk2* in synthesising NAD^+^ coupled with over-representation of NAD/NADH metabolism related enrichment terms for downregulated genes during both periods of atrophy, we next interrogated gene expression of other NAD^+^ biosynthesis genes and whether such alterations corresponded with a reduction in NAD^+^ content in both young adult (**Figure 7A-D**) and aged muscle (**Figure 7E-I**). In young adult humans, while most of these NAD^+^ genes were downregulated after both periods of atrophy, *NMRK2* was the only significantly downregulated DEG out of the 18 genes analysed (**Figures 7A-C**). Furthermore, *NMRK2* was the most upregulated gene across all DEGs after recovery from atrophy in humans (**Figure 3E**) and was among the identified DmDEGs that was downregulated less after repeated versus initial atrophy. Given *NMRK2* was the only significantly downregulated NAD^+^ gene, this resulted in no significant changes in NAD^+^ content (**Figure 7D**). In contrast, half of the 14 NAD^+^ genes assessed in aged animals were significantly differentially expressed where most genes were downregulated after atrophy, either unchanged/restored or increased after recovery, followed by greater downregulation after repeated atrophy (**Figure 7E-G**), thus resulting in significantly reduced absolute NAD^+^ content in the left (L) TTX versus right (R) control limb after repeated atrophy (**Figure 7H & I**). Collectively, these findings demonstrate that repeated atrophy suppresses multiple components of the NAD⁺ biosynthesis and salvage pathways, with NMRK2 emerging as a consistently downregulated gene across both species and analytical layers. Given that NMRK2 activity contributes directly to NAD⁺ regeneration and that its suppression occurred across atrophic episodes, we next sought to determine whether NMRK2-related deficits extend to human derived muscle stem cell function, and whether nicotinamide riboside (NR) supplementation could modulate NMRK1/2 expression and myogenic capacity in cells derived before and after atrophy.

**Figure 7.**
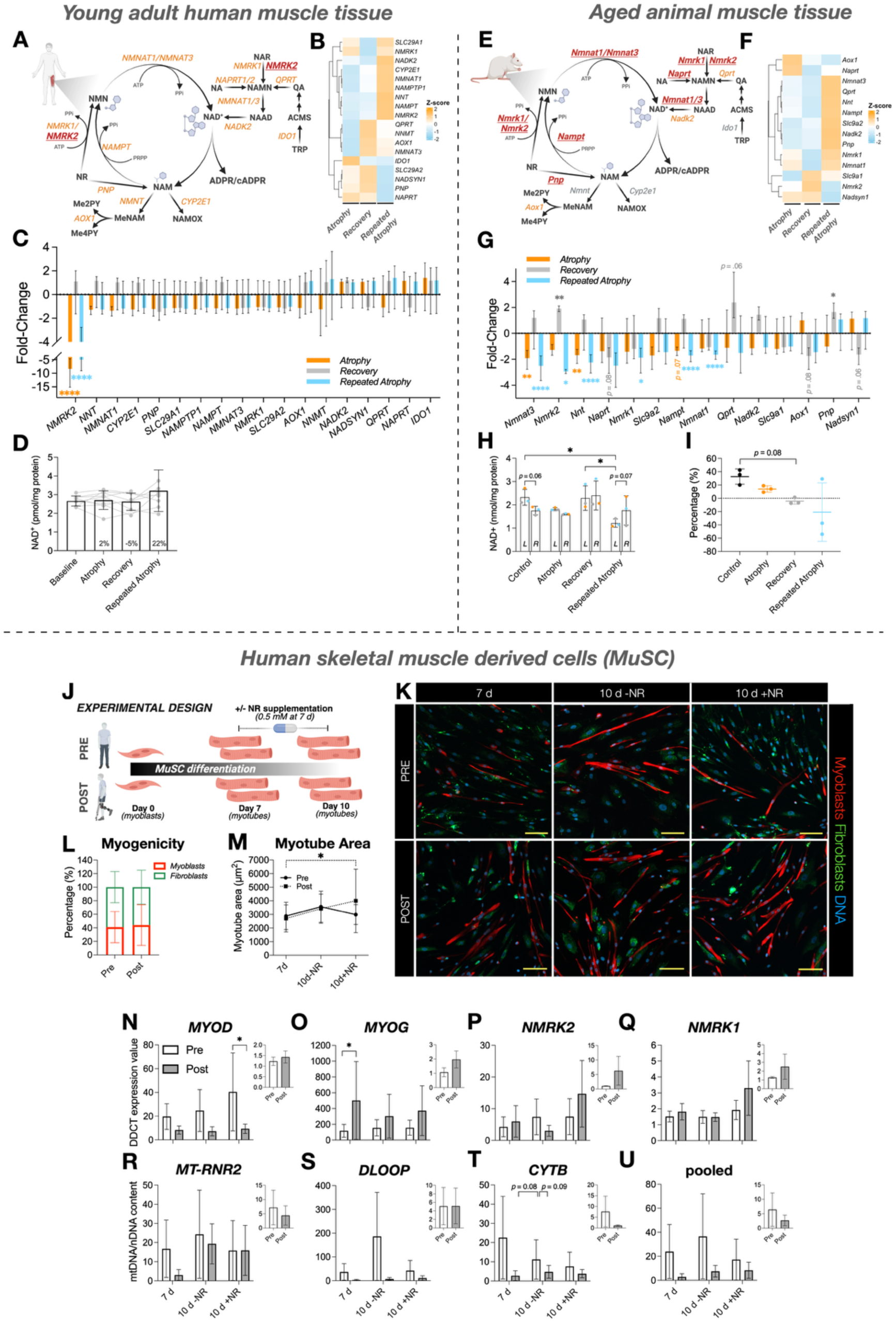
mRNA expression of NAD^+^ biosynthesis genes and corresponding NAD^+^ content in young adult human (**A-D**) and aged rat (**E-I**) skeletal muscle. (**A & E**) Schematic representation of Preiss-Handler and salvage NAD^+^ biosynthetic pathways (redrawn from Damgaard & Treebak, 2023^83^ using BioRender.com). Key metabolites are depicted as black text. Genes encoding NAD^+^ biosynthesis enzymes are shown in either orange (if non significantly expressed) or red text (significantly downregulated after atrophy or repeated atrophy). In young adult humans, *NMRK2* was the only significantly downregulated DEG out of the 18 NAD^+^ genes (**B & C**), which did not lead to significant alterations in NAD^+^ content (**D**). In aged animals, most of the 14 NAD+ genes assessed were significantly differentially expressed after atrophy and repeated atrophy and were restored upon recovery (**F & G**), resulting in significantly reduced absolute (**H**) NAD^+^ content in the left (L) TTX versus right (R) control limb after repeated atrophy. Relative NAD^+^ content (**I**). Nicotinamide riboside (NR) supplementation (0.5 mM) in human pre- and post-atrophy MuSC (*n* = 4) (J-K). (**K**) Fluorescence microscopic tile scan (3 × 3) images of MuSC labelled for myoblasts (desmin, red), fibroblasts (TE-7, green) and DNA (DAPI, blue), acquired at 10× magnification (scale bar = 200 µm). (**L**) Myoblasts (≈ 40%) and fibroblast (≈ 60%) proportions were similar between pre- and post-atrophy MuSC. NR supplementation resulted in significantly increased myotube area at 10 versus 7 d (*p* = 0.03) in post-atrophy MuSC only (**M**). qPCR analysis revealed greater *MYOD* expression in pre-atrophy 10 d NR treated MuSC (**N**) coupled with greater *MYOG* expression at 7 d post-atrophy with no effect of NR treatment (**O**). Despite greater average *NMRK2* (**P**) and *NMRK1* (**Q**) expression in 10 d NR supplemented post-atrophy MuSC, this was not significantly different from pre-atrophy or non-treated MuSC. (**R-U**) There was a trend towards lower mtDNA content in post-versus pre-atrophy MuSC at each timepoint/condition, as well as lower mtDNA in NR treated pre-atrophy MuSC only, albeit not statistically significant. Larger main figures relativised to 0 h, small inset figures represent 10 d NR treated relativised to 10 d non-treated MuSC. \**p* ≤ 0.05, \*\**p* ≤ 0.01, \*\*\**p* ≤ 0.001, \*\*\*\**p* ≤ 0.0001. Myogenicity (**L**) and myotube area (**M**) data presented as mean ± SD. Gene expression (**N-Q**) and mtDNA (**R-U**) data presented as mean ± standard error of the mean (SEM).

### Nicotinamide riboside (NR) enhances myogenic responses in human post-atrophy MuSCs

Because NMRK2 was among the most strongly downregulated genes after both atrophy periods in humans, and given its established activation by the vitamin B₃ derivative nicotinamide riboside (NR), we next examined whether NR supplementation could modulate NMRK1/2 expression and myogenic potential in muscle stem cells isolated from pre- and post-atrophy human biopsies (Figure 7J–M). Although Nmrk2 expression and NAD⁺ content were significantly reduced after repeated disuse in aged animals, these omics-driven discoveries were made after tissue allocation in the animal experiments, and muscle stem cells were therefore not available for parallel studies in aged rat muscle. Similar proportions of myoblast (desmin⁺) and fibroblast (TE7⁺) populations were observed between pre- and post-atrophy MuSCs, thereby negating differences in cell composition as a potential confounder (Figure 7L). Similar proportions of myoblast (desmin+) to fibroblasts (TE7+) were observed between pre- and post-atrophy MuSC, thus negating possible effects of cell populations (**Figure 7L**). Despite limited myotube formation in the MuSC, likely due to a higher proportion of fibroblasts (≈ 60%) versus myoblasts (≈ 60%) (**Figures 7K & L**), post-atrophy MuSC were more responsive to NR supplementation, where myotube size significantly increased in NR treated post-atrophy cells when compared to 7d differentiation (*p* = 0.03, **Figure 7M**). This was supported by lower *MYOD (early differentiation)* and greater *MYOG (fusion/differentiation)* gene expression in NR-supplemented MuSC, coupled with greater (albeit non-significant *NMRK1* and *NMRK2* expression (**Figures 7N-Q**). Interestingly, mtDNA content demonstrated lower mean (albeit non-significant) mtDNA in post-versus pre-atrophy MuSC (**Figures 7R-U**). Collectively, post-atrophy MuSC were more responsive to NR supplementation that altered *NMRK* and *MYOG* expression and increased myotube size.

Together, these findings indicate that post-atrophy muscle stem cells retain the capacity to respond to nicotinamide riboside through modulation of NMRK1/2 expression and myotube size, suggesting that NAD⁺ salvage influences myogenic potential following disuse. To determine whether these cellular effects correspond to broader alterations in mitochondrial pathways in vivo, and given the strong enrichment of mitochondrial and aerobic metabolism genes within the DmDEG clusters identified above, we next examined mitochondrial gene expression, protein abundance, and mtDNA content across atrophy, recovery, and repeated atrophy in both young adult human and aged rat skeletal muscle.

### Mitochondrial gene expression, protein abundance, and DNA content across atrophy, recovery, and repeated atrophy in young adult human and aged rat muscle

In addition to the NAD⁺ metabolism pathways described above, genes involved in mitochondrial structure, function, assembly, and turnover were strongly represented among the differentially methylated and differentially expressed clusters in both species. Specifically, genes involved in aerobic respiration (and thus ETC and oxidative phosphorylation) and mitochondrial structural components were hypermethylated and downregulated after both periods of atrophy, with the majority of these genes returning to baseline levels in both human and rat muscle despite restoration of muscle size in humans and continued wasting in aged rats. Furthermore, these genes demonstrated further reductions in transcription in aged rat muscle, whereas these genes were less susceptible to reductions in young adult human muscle following repeated disuse. Given these data, we next assessed gene expression of mitochondrial fission, fusion, transport and biogenesis genes-specific genes, as well as mRNA and protein abundance of ETC protein complexes I-V. Finally, we assessed mtDNA copy number and citrate synthase protein abundance as markers of mitochondrial content/density. In young adult human muscle (**Figure 8A**), the majority of the 66 genes profiled across ETC complexes I-V were significantly suppressed after both periods of atrophy where most genes either returned to baseline levels or increased after recovery (**Figure 8B**). This resulted in significant changes occurring after both atrophy and repeated atrophy in 24 (out of 66) ETC genes; within complex I (7 out of 32 genes), III (3 out of 8 genes), IV (8 out of 13 genes), and V (6 out of 9 genes) with non-significant changes in CII (**Figure 8B**). Concerning genes related to mitochondrial fission and mitophagy (*FIS1, DRP1*), fusion (*MFN1/2, OPA1*), transport (*SLC25A51, SLC25A4)* and biogenesis (*PPARGC1A, TFAM*), 8 out of 9 of these genes non-significantly decreased after atrophy and repeated atrophy, all of which returned to baseline levels or non-significantly increased after recovery (**Figure 8C**). There was a significant reduction in *PPIF*, a mitochondrial membrane gene controlling calcium overload and fatigue resistance in muscle (**Figure 8C**). Given the large transcriptional perturbations in ETC-specific genes across all 5 complexes in both atrophy periods, we next assessed ETC (I-V) protein abundance. The reductions in gene expression did not correspond to significant changes in ETC protein complex abundance (**Figure 8D**).

**Figure 8.**
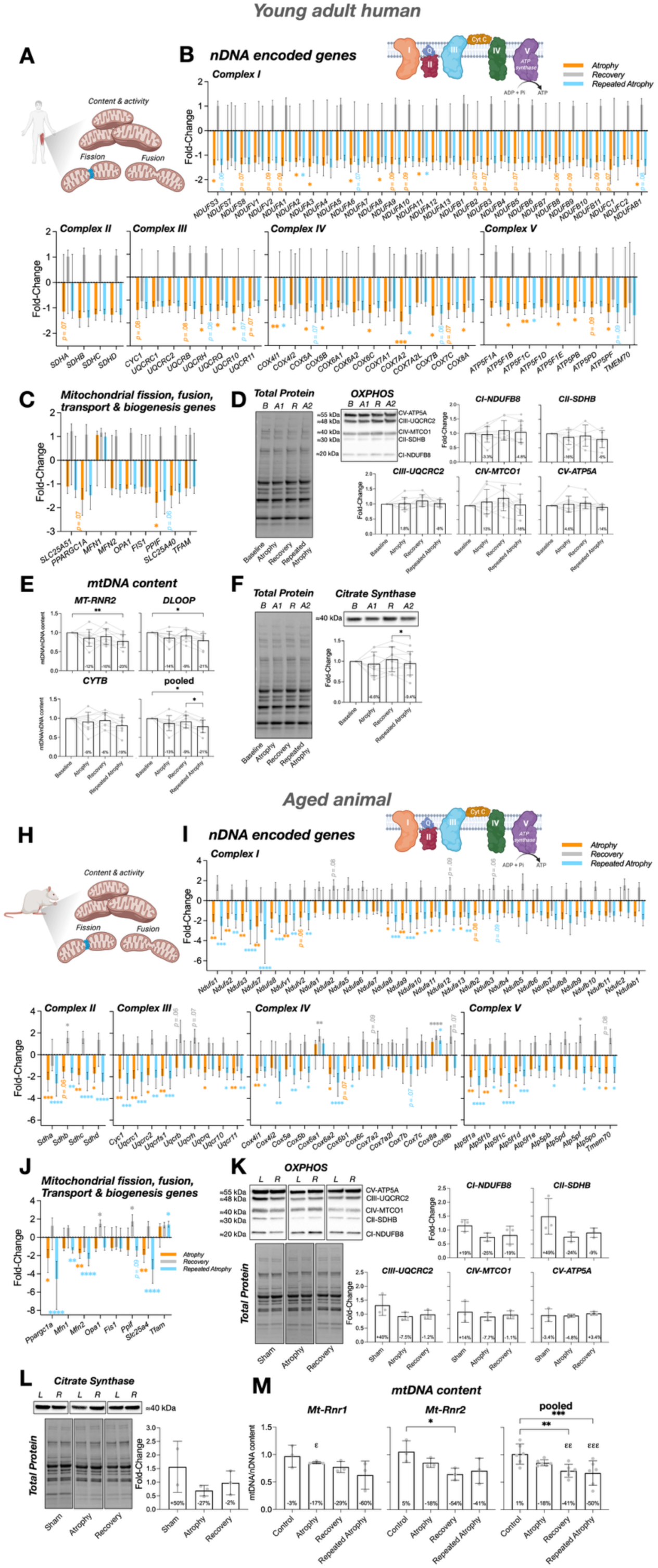
Mitochondrial response to repeated periods of disuse in young adult human muscle (**A-G**) and aged rat muscle (**H-M**) skeletal muscle. (**A & H**) Schematic depicting a specific focus on mitochondrial content as well as mitochondrial activity, fusion and fission processes (created using BioRender.com). (**B & I**) Temporal transcriptional response of ETC complex I-V genes where most genes significantly decreased after atrophy and/or repeated atrophy and recovered upon returning to normal activity in both adult humans (**B**) and aged animals (**I**). A similar transcriptional response occurred for mitochondrial fission, fusion, transport and biogenesis-related genes across species (**C & J**). There were no significant changes in ETC complex (I-V) protein abundance in both human (**D**) and rat (**K**) muscle. However, mtDNA content and citrate synthase protein significantly decreased after repeated atrophy in young adult humans (**E & F**). Despite no significant changes in citrate synthase across the sham/atrophy/repeated atrophy conditions analysed in aged animals (**L**), there was a reduction in mtDNA content in response to atrophy, recovery and repeated atrophy with the largest loss of -50% after repeated atrophy (**M**). Gene expression data presented as mean (signed) FC ± absolute 95% CI (absolute upper and lower limits). \**p* ≤ 0.05, \*\**p* ≤ 0.01, \*\*\**p* ≤ 0.001, \*\*\*\**p* ≤ 0.0001.

Interestingly however, mtDNA content (**Figure 8E**) and citrate synthase protein abundance (**Figure 8F**) significantly decreased after repeated atrophy only in young adult humans. Collectively, these data suggest that the large transcriptional perturbations of genes enriched in mitochondrial-specific pathways and GO terms culminated in impaired mitochondrial content and density following repeated disuse only in young adults despite restoration of these parameters and skeletal muscle during recovery.

In aged animals, most genes across all ETC complexes were differentially expressed (**Figure 8H & I**). Specifically, 57% (38 out of 67) of genes encoding complex’s I (13 out of 30), II (4 out of 4), III (7 out of 9), IV (6 out of 14) and V (8 out of 10 genes) were significantly downregulated after atrophy and/or repeated atrophy (**Figure 8H & I**). Similarly to young adult humans, the majority of these genes recovered during TTX cessation despite continued loss wasting during return to activity. Furthermore, *Sdhb, Cox6a1, Cox8a and Atp5pf* were significantly upregulated during recovery (**Figure 8I**). In contrast to young adult humans, aged rat muscle demonstrated a greater average reduction across all 67 genes following repeated atrophy (−1.72 FC) versus the initial atrophy period (−1.52 FC), despite a similar average FC between recovery and control (0.1 FC). Mitochondrial fusion (*Mfn1/*2), transport (*Slc25a4)* and biogenesis (*Ppargc1a*) genes also significantly decreased after atrophy and/or repeated atrophy whereas *Tfam* significantly increased after repeated atrophy (**figure 8J**). Most genes recovered during TTX cessation, and *Opa1* and *Ppif* genes both significantly increased during recovery (**Figure 8J**). As with the ETC genes described above, there was a greater average reduction across all genes (excluding *Tfam* which increased) after repeated atrophy (−2.23 FC) compared to previous atrophy (−1.56 FC) relating to a 43% larger reduction after repeated atrophy compared to initial atrophy, where most genes recovered upon returning to activity (average 0.72 FC) (**Figure 8I & J**). At the protein level, there were no significant changes in protein abundance of ETC complexes I-V nor citrate synthase across the conditions analysed (e.g., sham, atrophy and recovery respectively, **Figure 8L)**. Finally, there was a reduction in mtDNA content during atrophy (−18%), recovery (−41%) and the greatest reduction after repeated atrophy (−50%) when *Mt-Rnr1* and *Mt-Rnr2* genes were combined (**Figure 8M).**

Together, these findings demonstrate that repeated atrophy imposes substantial and age-dependent disruption of mitochondrial gene expression, protein abundance, and mtDNA content, with young adult muscle showing partial resilience and aged muscle exhibiting exaggerated susceptibility. These data complete our multi-layered assessment of physiological, transcriptional, epigenetic, metabolic, and mitochondrial responses to repeated disuse.

## Discussion

### Summary of main findings

Skeletal muscle exhibits a molecular memory of repeated disuse-induced atrophy. In young adult humans, repeated immobilization induced a comparable loss of skeletal muscle with full recovery between periods of disuse. Consistent with this, young adult rats also recovered muscle mass and fibre CSA after initial disuse, whereas aged rats did not, reinforcing the age-dependent divergence in recovery capacity. Repeated disuse triggered a protective molecular resilience where transcriptional perturbations of genes associated with oxidative metabolism and mitochondria (structure and function) were attenuated during recurrent atrophy. In contrast, aged muscle displayed a detrimental molecular susceptibility, characterised by further suppression of genes within similar pathways following repeated disuse, despite transcriptional recovery and continued muscle wasting upon returning to normal habitual activity. Proteasomal, ECM, and DNA damage associated genes were uniquely elevated following repeated atrophy in aged muscle. Methylome analyses revealed conserved epigenetic regulation of aerobic metabolism and mitochondrial genes across species, along with locus-specific memory genes including *NR4A1* and acetylcholine receptor subunit genes. Notably, nicotinamide riboside (NR) supplementation in human primary MuSC partially restored *NMRK1/2* expression and improved myotube size in MuSC derived post-atrophy. Importantly, repeated disuse in elderly humans is associated with frailty and increased fall risk, raising ethical concerns. Therefore, our design limited human experiments to young adults and employed aged rats as a surrogate model for repeated atrophy. In addition, to comply with the 3Rs (Reduction, Refinement, Replacement), we avoided unnecessary duplication by not including a direct young rat control group; instead, we integrated previously published young rat data ^66^ with newly generated aged rat data, enabling robust age comparisons while minimising animal use.

### Molecular memory of repeated disuse-induced skeletal muscle atrophy

Skeletal muscle can retain an epigenetic ‘memory’ of prior growth, as demonstrated by persistent hypomethylation during detraining in humans and mice ^33, 36, 37^, and amplified gene expression upon retraining ^33, 34, 36, 37^. Our findings now extend this concept to a clinically relevant negative stimulus. Specifically, disuse-induced atrophy which commonly occurs after injury, hospitalization, bed rest, surgery, or spaceflight. Crucially, we identified an age-dependent molecular memory: a *protective* transcriptional attenuation in young muscle, where repeated disuse elicited blunted transcriptional responses, and a *detrimental* memory in aged muscle, where repeated atrophy exaggerated suppression of NAD^+^, aerobic metabolism and mitochondrial gene pathways.

### Attenuated transcriptional memory in young adult humans and a detrimental exaggerated transcriptional memory in aged animal muscle

In young adult human muscle, genes suppressed during initial atrophy largely recovered and were less perturbed by repeated disuse, indicating a transcriptional protection or resilience. These responses predominantly involved oxidative metabolism and mitochondrial pathways, including TCA cycle, NAD^+^/NADH, ETC, ATP synthesis. In contrast, aged animal muscle showed the opposite pattern, in which these same pathways were further suppressed during repeated atrophy despite transcriptional recovery after the initial disuse, accompanied by unique activation of proteasomal, ECM, and DNA damage gene pathways after repeated atrophy. These signatures are consistent ageing-associated skeletal muscle deficits in metabolic regulation, proteostasis, NMJ dysfunction (discussed below) and mitochondrial function ^10, 19, 84^. Importantly, expression of these gene pathways recovered even though muscle mass continued to decline in aged muscle, confirming that the exaggerated transcriptional and physiological responses reflected a detrimental molecular memory rather than incomplete recovery of these molecular pathways.

### Mitochondrial vulnerability and loss of mtDNA with repeated disuse atrophy

Repeated disuse caused significant loss of mtDNA that was not evident after the initial atrophy period in either young adults or aged animals, with the most pronounced decline observed in aged muscle. These findings align previous bed rest studies demonstrating broad downregulation of mitochondria and reduced content/respiration during unloading ^24, 64, 85^. Our transcriptomic data also revealed widespread suppression of electron transport chain and mitochondrial genes during atrophy and repeated atrophy, followed by recovery upon reloading/ return to activity. Although aged muscle exhibited the greatest mitochondrial vulnerability, young muscle also exhibited a marked loss of mtDNA loss after repeated atrophy, despite the transcriptional protection discussed above. This suggests that repeated disuse increases susceptibility to mitochondrial loss, raising the question as to whether further atrophic episodes of disuse atrophy could drive even young muscle toward the detrimental molecular program observed in aged muscle.

### A similar epigenetic program in both species represses aerobic metabolism genes

Integrative methylome and transcriptome analyses revealed differentially methylated, differentially expressed genes (DmDEGs) enriched in oxidative metabolism and mitochondrial structure/turnover in both humans and rats. These genes were hypermethylated and downregulated during atrophy and repeated atrophy, returning toward baseline on recovery. This extends prior observations of disuse-linked methylation (e.g., nNOS hypermethylation ^65^, *Chrna1* hypomethylation/upregulation ^66^) and contrasts with the hypomethylating effects of exercise training, higher physical activity and aerobic fitness associated with a ‘younger’ skeletal muscle methylome ^36, 37, 39, 48, 57, 58, 59, 60, 63^.

### Neuromuscular signalling pathways in response to repeated atrophy: Epigenetic priming of the AChR subunit genes

Repeated disuse increased expression of acetylcholine receptor subunits, *CHRNA1, CHRND* in young human muscle, alongside hypomethylation at these loci, and similarly elevated *Chrna1, Chrnd, and Chrng* expression in aged rat muscle. Although potentially compensatory, such extra-junctional expression, especially of the fetal-type gamma subunit (*Chrng*), is typically linked to NMJ dysfunction rather than stabilization ^10^. In young muscle, this response emerged only after repeated atrophy, coinciding with the strongest hypomethylation, suggesting an epigenetic memory within AChR genes. Aged muscle demonstrated over a doubling in the upregulation of *Chrna1* (16-fold) and *Chrnd* (12.5-fold) after repeated disuse compared with initial disuse (*Chrna1* 7-fold*, Chrnd*-5.8 fold), and only significant elevation of *Chrng* after repeated atrophy. Whilst we did not detect hypomethylation of these genes in aged rat muscle, prior studies have also reported *Chrna1* hypomethylation in young adult rats utilised the same model of disuse ^66^. Therefore, despite general transcriptional resilience, young muscle still exhibited exaggerated AChR gene responses to repeated atrophy. Further, the exaggerated response in aging muscle is consistent with aging studies associating poor recovery with NMJ instability and ER stress ^10, 26, 86^, and imply that repeated stress, even in young muscle, may epigenetically prime these genes, heightening vulnerability to future disuse.

### NR supplementation partially restored NMRK2 levels and increased myotube size in human MuSC derived post-atrophy

*NMRK2*, a key NAD⁺ salvage enzyme, was among most downregulated genes after both periods of atrophy in young adult humans and was significantly downregulated in aged rat muscle after repeated atrophy only, with broader NAD^+^/NADH metabolism pathway repression across species, most pronounced in aged muscle. In primary human MuSC derived post-atrophy, nicotinamide riboside (NR) partially restored *NMRK2* and improved *MYOG* expression and myotube size, suggesting NAD⁺ salvage as a potential target. However, whilst this provides initial *in-vitro* evidence, preclinical work shows intravenous NR elevates muscle NAD⁺, while oral NR trials report mixed efficacy ^83, 87^. Rigorous clinical studies in older, immobilization-prone populations are essential before considering NR as a viable therapeutic intervention.

### ECM remodelling during recovery and repeated atrophy in aged muscle

Only aged muscle upregulated genes linked to extracellular matrix (ECM), collagen organization, collagen binding and sarcomere structure during recovery from initial disuse. This suggests structural remodelling as a compensatory response, which may stabilize tissue but also increase stiffness ^88, 89^, a hallmark of muscle aging, and therefore perhaps impair functional recovery. Previous work has shown that repeated cycles of atrophy in rats can lead to structural remodeling of the extracellular matrix (ECM), which may influence muscle resilience and susceptibility to future disuse ^90^. In the present study, ECM turnover, proteasomal pathways, and ubiquitin-mediated proteolysis were exclusively enriched after repeated atrophy in aged muscle, reinforcing the concept of compromised resilience to repeated disuse with age.

### The NR4A genes are important metabolic regulators of repeated disuse atrophy in young human muscle

*NR4A1*, a key energy metabolism gene, was hypermethylated and remained downregulated even after full recovery in young muscle, while *NR4A3* showed the largest reduction of all altered genes (ranked 1^st^ out of 384). Temporal profiling confirmed both genes retained suppression post-atrophy, indicating transcriptional memory. NR4A genes are highly responsive to exercise and inactivity ^91^; *NR4A1* is oppositely hypomethylated and upregulated after high-intensity exercise ^46^, and *NR4A3* overexpression in rats enhances mitochondrial content, ATP production, and type-IIA & IIX muscle fibre protein content ^92, 93, 94^. Knockdown of *NR4A3* impairs glucose metabolism and protein synthesis in skeletal muscle ^95^. Collectively, these findings nominate *NR4A* genes as candidate targets to improve recovery and protect against future disuse atrophy.

### Strengths and Limitations

This study provides a comprehensive physiological and molecular analysis of repeated disuse atrophy across young adult human and aged rat skeletal muscle. However, several limitations should be acknowledged. 1) Interpretation of molecular protection vs. atrophy outcomes: While we observed transcriptional attenuation in young human muscle following repeated disuse, this did not translate into attenuated muscle mass loss. The term “protective molecular memory” refers specifically to reduced transcriptional responsiveness in metabolic pathways, not an attenuated loss in skeletal muscle size due to repeated atrophy. Further, whilst there was a predominance of molecular attenuation in young human muscle to repeated atrophy, there were also more susceptible losses, such as further mtDNA content and larger increases in AChR subunit gene expression (discussed above). However, this raises the interesting question as to whether further atrophic episodes of disuse atrophy could drive even young muscle toward the detrimental program observed in aged muscle. 2) Coss-species comparisons and protocol differences: Although both models were analysed independently before integration, differences in species, disuse protocols (immobilization vs. TTX), recovery durations, and age must be considered when interpreting cross-species comparisons. Although the human and rat models differ in species, age, and the method of disuse induction (immobilisation versus TTX), both exhibited highly conserved molecular signatures in key pathways, including aerobic metabolism, mitochondrial function, and NAD⁺ metabolism; these similarities were unexpected and suggest that molecular memory may represent a shared biological phenomenon across species. Nevertheless, these cross-species comparisons should be interpreted cautiously and considered exploratory, as biological variability between species, strains, and age groups may influence interpretation. Further, young and aged rats compared were of different strains (Wistar vs. Fischer 344), which may influence physiological and molecular responses to TTX-induced disuse independently of age. This should be considered when interpreting cross-strain comparisons. Nevertheless, transcriptomic analysis revealed substantial overlap in downregulated genes during atrophy in both young adult and aged rats (Figure 5), suggesting that the core molecular response to disuse is largely conserved across strains. The greatest divergence occurred during recovery, where young rats regained muscle mass, whereas aged rats continued to lose muscle mass despite transcriptional recovery. This indicates that age, rather than strain, is the primary determinant of impaired recovery following repeated disuse. The study also used male animals, which may restrict generalisability and preclude assessment of sex-specific responses. This choice was driven by practical and welfare considerations: animals were imported at approximately 18 months of age and housed for ∼5-6 months before experimentation, and male rats were selected because they could be socially housed in groups of two or three during this acclimation period, whereas female rats were less likely to form stable group pairings after import. In addition to this sex-related constraint, a relatively small group size of aged rats was used. To address this, the unilateral TTX model was employed because it offers a methodological advantage: the contralateral limb remains fully innervated and untreated, allowing each animal to serve as its own internal control. This paired design substantially increases statistical power and reduces between-animal variability, enabling robust detection of physiological and molecular changes with fewer animals and aligning with the principles of Reduction and Refinement. This was particularly important given the four experimental timepoints and the use of very old Fischer 344 rats, where availability was inherently limited. Although bilateral hindlimb unloading remains a valuable model of disuse, it affects both limbs simultaneously and therefore typically requires larger group sizes to achieve comparable statistical sensitivity. The unilateral approach used here was therefore selected to maximise statistical robustness while minimising total animal use. Finally, although protocol differences exist between young and aged rat studies, the observation that young rats recovered muscle mass after a slightly longer TTX exposure and shorter recovery period, whereas aged rats failed to recover despite a shorter exposure and extended recovery, strongly suggests that age is the primary determinant of impaired recovery. The extreme age of the Fischer 344 rats (∼23 months) likely further contributed to this phenotype. 3) Recovery protocol and activity monitoring: In aged rats, molecular pathways recovered during TTX cessation despite continued muscle wasting, suggesting that the transcriptional response was TTX-atrophy dependent rather than a consequence of incomplete recovery. Further, we present a comparison of the aged rats with young adult rats ^66^ that recover half of their muscle mass following a slightly longer period of TTX-atrophy (and thus greater muscle wasting than aged animals herein) in a shorter recovery period. However, whilst rat activity was confirmed by the animals being able to perform dorsiflexion, activity levels during recovery was not directly measured. Future studies should incorporate objective tracking of habitual activity (for example, respirometry or automated behavioural monitoring) together with detailed nutritional intake measures to fully disentangle behavioural influences from molecular and physiological recovery. Further, during human immobilization considering wearables (accelerometers) or bed-rest protocols to better characterize recovery dynamics and disentangle molecular recovery from functional outcomes. It is also worth noting that recovery dynamics after unloading have been previously characterised in young versus old rats ^96^, where older rats exhibit impaired but measurable recovery. In contrast, the very old Fischer 344 rats used in our study (∼23 months) continued to lose muscle mass during recovery, suggesting that advanced age may exacerbate susceptibility to repeated disuse. 4) Methodological constraints and data integration: The integration of multi-omics data across species and models introduces complexity. While individual analyses are robust, interpretation of age-related effects must consider differences. For example, due to limited fibre number in the cross-sections from human samples we also inferred fibre proportions using MYH and MYL gene expression data. Further, ETC protein data were not available for repeated atrophy in aged rats, and ETC protein changes were not statistically significant despite transcriptomic, NAD⁺ content and mtDNA evidence of impairment. While our integration strategy used SOM profiling to cluster DEGs and DMRs temporally, advanced computational multi-omic frameworks such as BETA ^43^ could provide deeper mechanistic insights. Implementing such approaches was beyond the scope of the manuscript but represents an important avenue for future work. 5) Translational scope and *in vitro* validation: NR supplementation partially restored *NMRK2* expression and myotube size in post-atrophy human MuSCs, suggesting a potential therapeutic avenue with translation to humans. However, these findings are limited to *in vitro* experiments in young adult cells. Further studies are needed to validate NAD⁺ salvage interventions in aged muscle and *in vivo* models of repeated disuse. Further, these *in vitro* experiments were performed in differentiating MuSCs and developing myotubes rather than fully mature myotubes, and MuSC isolation from aged rat muscle was not possible as these experiments were driven by omics discovery after cell isolation had been completed, which precluded parallel aged rat studies. Nevertheless, the data provide preliminary insight into NAD⁺ salvage and NMRK2 responsiveness, including partial recovery of NMRK2 (and NMRK1) expression, modulation of MYOD and MYOG, and improved myotube size with NR supplementation. Future studies should address targeted manipulation of NMRK2 in aged rat myotubes or aged rat tissue *in-vivo* under atrophic conditions to provide important mechanistic insight into NR’s role in protecting against disuse-induced atrophy. 6) Epigenetic interpretation and functional relevance: While DNA methylation changes were inversely associated with gene expression in key metabolic pathways, functional consequences of these epigenetic modifications remain to be fully elucidated. To complement the descriptive multi-omics dataset, we incorporated targeted validation of mitochondrial function, including gene expression profiling, NAD⁺ content, mtDNA copy number, citrate synthase activity, and ETC protein abundance. While most measures aligned with the omics findings, some species-specific differences were evident, such as unchanged NAD⁺ content in humans despite reductions in aged rats, and unchanged ETC protein abundance despite consistent transcriptional suppression of mitochondrial pathways. These discrepancies likely reflect assay limitations and biological variability. Nevertheless, the strong concordance between omics and targeted analyses at the pathway level reinforces the robustness of the findings. Future work should explore whether these methylation signatures directly modulate transcriptional memory or serve as biomarkers of prior disuse. In addition, structural correlates were not assessed in detail. A limitation of the study is the absence of additional histological analyses of denervation markers (e.g., NCAM, eMyHC, central nucleation) and extracellular matrix (ECM) organisation in the repeated atrophy rat model, which would have provided further corroboration of the omics data. These analyses were not feasible within the scope of the current study, as all available tissue was utilised for immunohistochemistry, RNA, DNA, and protein isolation, primary cell isolation, and NAD⁺ content assays. Future studies should incorporate detailed histological assessments of neuromuscular junction integrity and ECM remodelling alongside multi-omic profiling to strengthen mechanistic interpretation.

### Future mechanistic and clinical directions

We propose mechanistic and translational studies that address testable hypothesis, such as: (i) Does enhancing NAD⁺ salvage attenuate the maladaptive transcriptional response to repeated disuse in aged muscle? ii) Does exercise ‘preconditioning’ ^97, 98^ or training reprogram methylation signatures to prevent negative molecular memory in the nuclear ^39, 57^ and mitochondrial genome ^63^? (iii) does targeted activation of *NR4A* family and *NMRK2* genes during atrophy or recovery prevent or offset disuse-induced atrophy and/or promote recovery from atrophy?

## Conclusion

Disuse atrophy can be remembered at the molecular level in skeletal muscle. It is predominantly protective in young skeletal muscle yet detrimental in aged skeletal muscle. This memory is partially epigenetically encoded at the single locus level and converges in aerobic metabolism, mitochondrial and NAD⁺ pathways, and nominates molecular targets to preserve muscle mass and function after recurrent immobilization or disuse.

## Acknowledgements

This research was funded by the Research Council of Norway (RCN) (314157) awarded to principal investigator Adam P. Sharples, post-doc fellow Daniel C. Turner and co-investigators Sue Bodine, Jonathan C. Jarvis, David C. Hughes, Daniel J. Owens, Truls Raastad and Olivier Seynnes. We thank Olivier Seynnes for his critical expertise and input into the funding application, training & supervision of post-doctoral researcher for the design, testing and analysis of the human experiments and associated physiological data acquisition and analysis (muscle size/CSA & strength). We thank Jostein Hallèn, Siri T. Dalen, Hege N. Ødemark and Emil Løvstakken (NIH) for assistance with muscle biopsies and tissue processing; Josephine A. Eyrich (UCPH) for assisting with the measurements of NAD^+^ content; Rune Enger, as the medical lead for the human biopsies; Enovis/DJO LLC for kindly donating the knee braces and to; ChromaDex, Inc., a subsidiary of Niagen Bioscience for providing nicotinamide riboside for *in vitro* studies. Finally, sincere gratitude to all participants who volunteered to participate in this study that included two periods of muscle wasting.

## Author contributions

Adam P. Sharples conceived the project, secured funding as principal investigator, and led the human experimental design, ethical approvals, data/finance management, dissemination, and oversaw analysis of all human and rat experiments, and supervised postdoctoral fellow Daniel C. Turner. Daniel C. Turner conducted most of the experimental work, including recruitment, human testing, biopsy processing, wet lab work, data collection, analysis, visualization, dissemination and manuscript drafting. Jonathan C. Jarvis supervised and undertook the animal intervention studies with Hazel Sutherland and provided vital input into the funding application and expertise in experimental design, analysis and interpretation. Sue C. Bodine, David C. Hughes, Daniel J Owens, and Truls Raastad were co-investigators in the funding application and heavily involved in study design, biopsy sampling, interpretation of results, and manuscript preparation. Stian F. Christiansen undertook experimental work for the mtDNA analysis in MuSC. Jonas T. Treebak and Emilie D. Kellezi undertook NAD^+^ content experiments. Eva Wozniak, Charles Mein and James Boot undertook RNA-seq and RRBS/RREM wet lab experiments and pre-processing of RNA-seq, RRBS/RREM and bioinformatics. Max Ullrich assisted in omics analysis and visualization. All authors contributed significantly to the final manuscript.

## Data availability

Raw and pre-processed RNA-seq and bisulfite sequencing data will be released via the Gene Expression Omnibus (GEO) repository (Human RNA-seq: GSE310344; Human Bisulfite-seq: GSE310743; Rat RNA-seq: GSE310848; Rat Bisulfite-seq: GSE310849). Any other processed data are available upon reasonable request to the corresponding authors.

## Software and code availability

All analysis code used for preprocessing, normalization, differential expression/methylation, enrichment and visualization of the data is freely available/open source with links and references detailed in **Supplementary Table 2.** Any unique code is available on reasonable request to the corresponding authors.

