## Supplementary Table 1 for "Repeated disuse atrophy imprints a molecular memory in skeletal muscle: transcriptional resilience in young adults and susceptibility in aged muscle"

**Supplementary Table 1.** Pre-surgical and post-intervention animal body weights (g). Values presented as mean ± standard deviation (SD).

| **Condition** | **Pre** | **Post** |
|---|---|---|
| Sham/Control *(no TTX)* | 374 ± 37 | 325 ± 102 |
| Atrophy *(6-7 d TTX)* | 378 ± 60 | 357 ± 36 |
| Recovery *(6-7 d TTX + 9 d recovery)* | 434 ± 31 | 392 ± 28 |
| Repeated Atrophy *(6-7 d TTX + 9 d recovery + 5-6 d TTX)* | 420 ± 9 | 344 ± 7* |
| Mean | 402 | 354** |
| SD | 43 | 54 |

TTX, tetrodotoxin. * depicts significant reductions in post- versus pre-body weight in repeated atrophy and mean of all conditions. *$p \leq 0.05$, **$p \leq 0.01$.
