## Supplementary Table 2 for "Repeated disuse atrophy imprints a molecular memory in skeletal muscle: transcriptional resilience in young adults and susceptibility in aged muscle"

**Supplementary Table 2.** Software and Code Availability

| Package name | Version | Link | Reference |
|---|---|---|---|
| openxlsx | v4.2.8 | https://CRAN.R-project.org/package=openxlsx | Schauberger P, Walker A (2025). _openxlsx: Read, Write and Edit xlsx Files_. |
| readxl | v1.4.5 | https://CRAN.R-project.org/package=readxl | Wickham H, Bryan J (2025). _readxl: Read Excel Files_. |
| stringr | v1.5.2 | https://CRAN.R-project.org/package=stringr | Wickham H (2025). _stringr: Simple, Consistent Wrappers for Common String Operations_. |
| dplyr | v1.1.4 | https://CRAN.R-project.org/package=dplyr | Wickham H, François R, Henry L, Müller K, Vaughan D (2023). _dplyr: A Grammar of Data Manipulation_. |
| annotatr (used for RNA-seq) | v1.32.0 | https://academic.oup.com/bioinformatics/article/33/15/2381/3092365 | Cavalcante RG, Sartor MA. annotatr: genomic regions in context. Bioinformatics. (2017) 33(15):2381-2383. doi:10.1093/bioinformatics/btx183 |
| STAR | v2.7.3a | https://doi.org/10.1093/bioinformatics/bts635 | Dobin A, *et al.* STAR: ultrafast universal RNA-seq aligner. *Bioinformatics* **29**, 15-21 (2013) |
| DeepVenn | Open source-Most recent | https://www.deepvenn.com/ | Hulsen T. DeepVenn - a web application for the creation of area-proportional Venn diagrams using the deep learning framework Tensorflow.js. *ArXiv* **abs/2210.04597**, (2022). |
| DAVID | Open source-Most recent | https://davidbioinformatics.nih.gov/ | Huang DW, Sherman BT, Lempicki RA. Systematic and integrative analysis of large gene lists using DAVID bioinformatics resources. *Nature Protocols* **4**, 44-57 (2009). |
| cowplot | V1.2.0 | https://CRAN.R-project.org/package=cowplot | Wilke C (2025). _cowplot: Streamlined Plot Theme and Plot Annotations for 'ggplot2'_. |
| org.Hs.eg.db | v3.20.0 | https://doi.org/10.18129/B9.bioc.org.Hs.eg.db | Carlson M (2024). _org.Hs.eg.db: Genome wide annotation for Human_. R package version 3.20.0. |
| org.Rn.eg.db | v3.21.0 | https://doi.org/doi:10.18129/B9.bioc.org.Rn.eg.db | Carlson M (2024). _org.Rn.eg.db: Genome wide annotation for Rat_. R package version 3.20.0. |
| ggplot2 | v4.0.0 | https://ggplot2.tidyverse.org | H. Wickham. ggplot2: Elegant Graphics for Data Analysis. Springer-Verlag New York, 2016. |
| ComplexHeatmap | v2.22.0 | http://bioconductor.org/packages/ComplexHeatmap/ | 1. Gu, Z. Complex Heatmap Visualization. iMeta 2022.<br>2. Gu, Z. Complex heatmaps reveal patterns and correlations in multidimensional genomic data. Bioinformatics 2016. |
| circlize | v0.4.16 | https://cran.r-project.org/package=circlize | Gu, Z. circlize implements and enhances circular visualization in R. Bioinformatics 2014. |
| ComplexUpset | v1.3.6 | http://doi.org/10.5281/zenodo.3700590 | Alexander Lex, Nils Gehlenborg, Hendrik Strobelt, Romain Vuillemot, Hanspeter Pfister, UpSet: Visualization of Intersecting Sets, IEEE Transactions on Visualization and Computer Graphics (InfoVis '14), vol. 20, no.12, pp. 1983–1992, 2014. |
| enrichplot | v1.26.6 | https://doi.org/10.18129/B9.bioc.enrichplot | Yu G (2025). _enrichplot: Visualization of Functional Enrichment Result_. doi:10.18129/B9.bioc.enrichplot |
| clusterProfiler | v4.14.6 | https://yulab-smu.top/contribution-knowledge-mining/ | T Wu, E Hu, S Xu, M Chen, P Guo, Z Dai, T Feng, L Zhou, W Tang, L Zhan, X Fu, S Liu, X Bo, and G Yu. clusterProfiler 4.0: A universal enrichment tool for interpreting omics data. The Innovation. 2021, 2(3):100141 |

| | | | |
|---|---|---|---|
| DOSE | v4.0.1 | https://yulab-smu.top/contribution-knowledge-mining/ | Guangchuang Yu, Li-Gen Wang, Guang-Rong Yan, Qing-Yu He. DOSE: an R/Bioconductor package for Disease Ontology Semantic and Enrichment analysis. Bioinformatics. 2015, 31(4):608-609 |
| ReactomePA | v1.50.0 | https://yulab-smu.top/contribution-knowledge-mining/ | Guangchuang Yu, Qing-Yu He. ReactomePA: an R/Bioconductor package for reactome pathway analysis and visualization. Molecular BioSystems. 2016, 12(2):477-479 |
| enrichplot | v1.26.6 | https://yulab-smu.top/contribution-knowledge-mining/ | Guangchuang Yu, Li-Gen Wang, Guang-Rong Yan, Qing-Yu He. DOSE: an R/Bioconductor package for Disease Ontology Semantic and Enrichment analysis. Bioinformatics. 2015, 31(4):608-609 |
| TxDb.Rnorvegicus.UCSC.rn6.refGene | v3.4.6 | https://doi.org/doi:10.18129/B9.bioc.TxDb.Rnorvegicus.UCSC.rn6.refGene | Team BC, Maintainer BP (2019). _TxDb.Rnorvegicus.UCSC.rn6.refGene: Annotation package for TxDb object(s)_. |
| TxDb.Hsapiens.UCSC.hg38.knownGene | v3.20.0 | 10.18129/B9.bioc.TxDb.Hsapiens.UCSC.hg38.knownGene | Team BC, Maintainer BP (2024). _TxDb.Hsapiens.UCSC.hg38.knownGene: Annotation package for TxDb object(s)_. |
| annotatr (used for methylation) | v1.34.0 | https://doi.org/10.18129/B9.bioc.annotatr | Cavalcante RG, Sartor MA (2017). "annotatr: genomic regions in context." Bioinformatics. R package version 1.34.0. |
| bcl2fastq /2.19.1 | v2.19.1 | https://support.illumina.com/sequencing/sequencing_software/bcl2fastq-conversion-software.html | https://support.illumina.com/content/dam/illumina-support/documents/downloads/software/bcl2fastq/bcl2fastq-2-19-1-release-notes-1000000035330-00.pdf |
| umi_tools | v1.1.4 (human) v1.1.6 (rat) | https://doi.org/10.1101/gr.209601.116 | Smith T, Heger A, Sudbery I. UMI-tools: modeling sequencing errors in Unique Molecular Identifiers to improve quantification accuracy. Genome Res 27, 491-499 (2017). |
| trim_galore | v0.6.10 | https://doi.org/10.5281/zenodo.5127898 | Felix Krueger, Frankie James, Phil Ewels, Ebrahim Afyounian, Michael Weinstein, Benjamin Schuster-Boeckler, Gert Hulselmans, & sclamons. (2023). FelixKrueger/TrimGalore: v0.6.10 - add default decompression path (0.6.10). Zenodo. https://doi.org/10.5281/zenodo.7598955 |
| bismark/0.22.1 - requires- bowtie2; includes- deduplicate_bismark; bismark_methylation_extractor | v0.22.1 | https://doi.org/10.1093/bioinformatics/btr167 | Krueger F, Andrews SR. Bismark: a flexible aligner and methylation caller for Bisulfite-Seq applications. Bioinformatics 27, 1571-1572 (2011). |
| bowtie2 | v2.4.1 | https://github.com/BenLangmead/bowtie2 | Langmead B, Salzberg S. Fast gapped-read alignment with Bowtie 2. *Nature Methods*. 2012, 9:357-359. |
| methylKit -includes calculateDiffMeth; percMethylation | v1.32.1 | http://doi.org/10.18129/B9.bioc.methylKit | Altuna Akalin, Matthias Kormaksson, Sheng Li, Francine E Garrett-Bakelman, Maria E Figueroa, Ari Melnick and Christopher E Mason. methylKit: a comprehensive R package for the analysis of genome-wide DNA methylation profiles. Genome Biology 13:R87 (2012). |
