## Supplementary Figure 1 for "Repeated disuse atrophy imprints a molecular memory in skeletal muscle: transcriptional resilience in young adults and susceptibility in aged muscle"

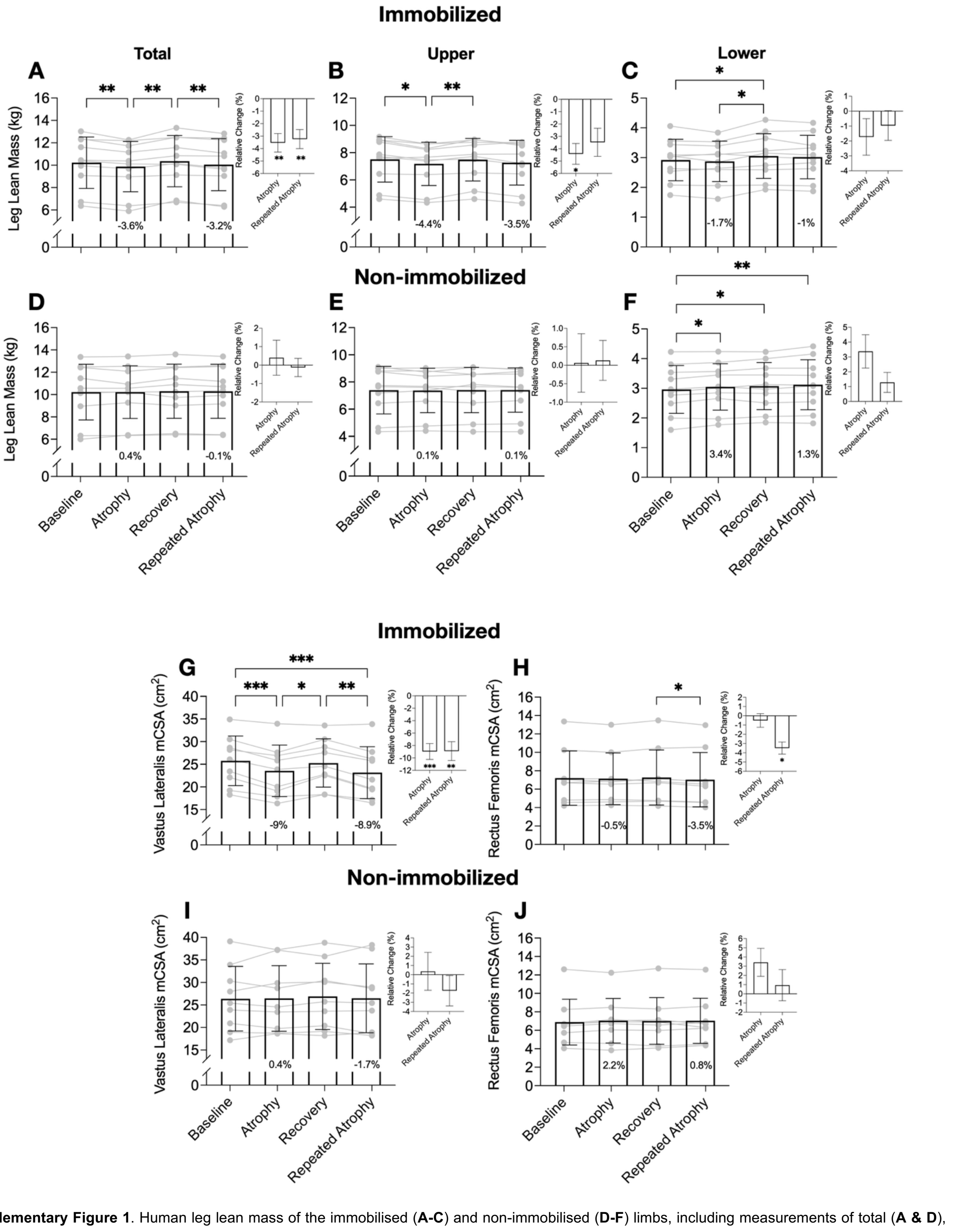


**Supplementary Figure 1**. Human leg lean mass of the immobilised (**A-C**) and non-immobilised (**D-F**) limbs, including measurements of total (**A & D**), upper (**B & E**) and lower (**C & F**) leg lean mass. Bar graphs depict relative change (%) from the previous timepoint (i.e., atrophy versus baseline, repeated atrophy versus recovery). *N* = 10. **p* ≤ 0.05, ***p* ≤ 0.01. Human mCSA of the immobilised (**G-H**) and non-immobilised (**I-J**) limbs, for *VL* (**G & I**) and *RF* (**H & J**) muscles. Bar graphs depict relative change (%) from the previous time point (i.e., atrophy versus baseline, repeated atrophy versus recovery). *N* = 9. **p* ≤ 0.05, ***p* ≤ 0.01, ****p* ≤ 0.001.
