## Supplementary Figure 2 for "Repeated disuse atrophy imprints a molecular memory in skeletal muscle: transcriptional resilience in young adults and susceptibility in aged muscle"

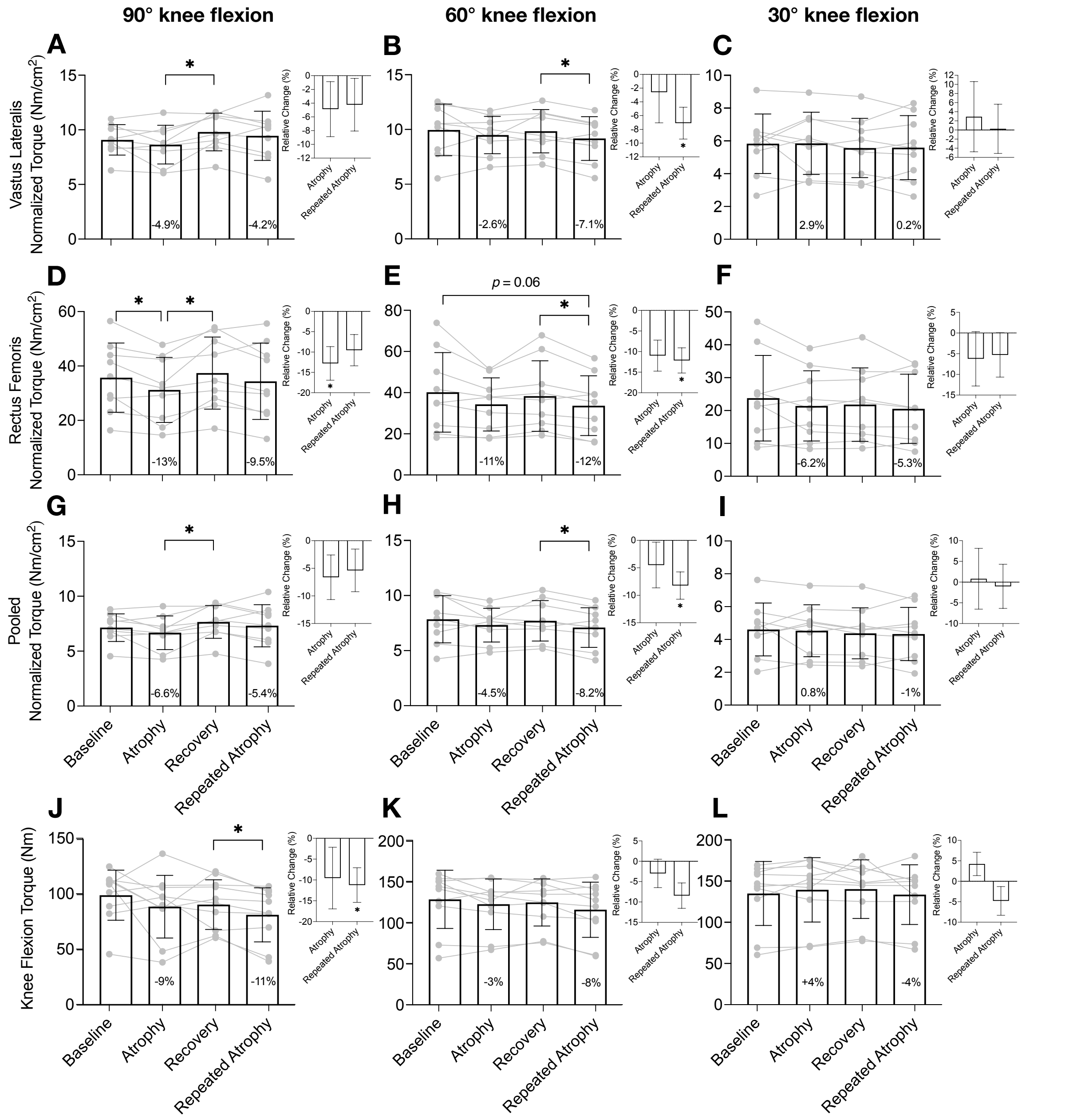


**Supplementary Figure 2**. Human specific knee extension strength (**A-I**). Knee extension torque normalised to mCSA of the *VL* (**A-C**), *RF* (**D-F**) and sum of *VL* + *RF* (**G-I**) at 90° (**A/D/G**), 60° (**B/E/H**) and 30° (**C/F/I**) knee flexion. Knee flexion torque at 90° (**J**), 60° (**K**) and 30° (**L**) knee flexion. Bar graphs depict relative change (%) from the previous time point (i.e., atrophy versus baseline, repeated atrophy versus recovery). *N* = 10. *$p \leq 0.05$.
