## Supplementary Figure 3 for "Repeated disuse atrophy imprints a molecular memory in skeletal muscle: transcriptional resilience in young adults and susceptibility in aged muscle"

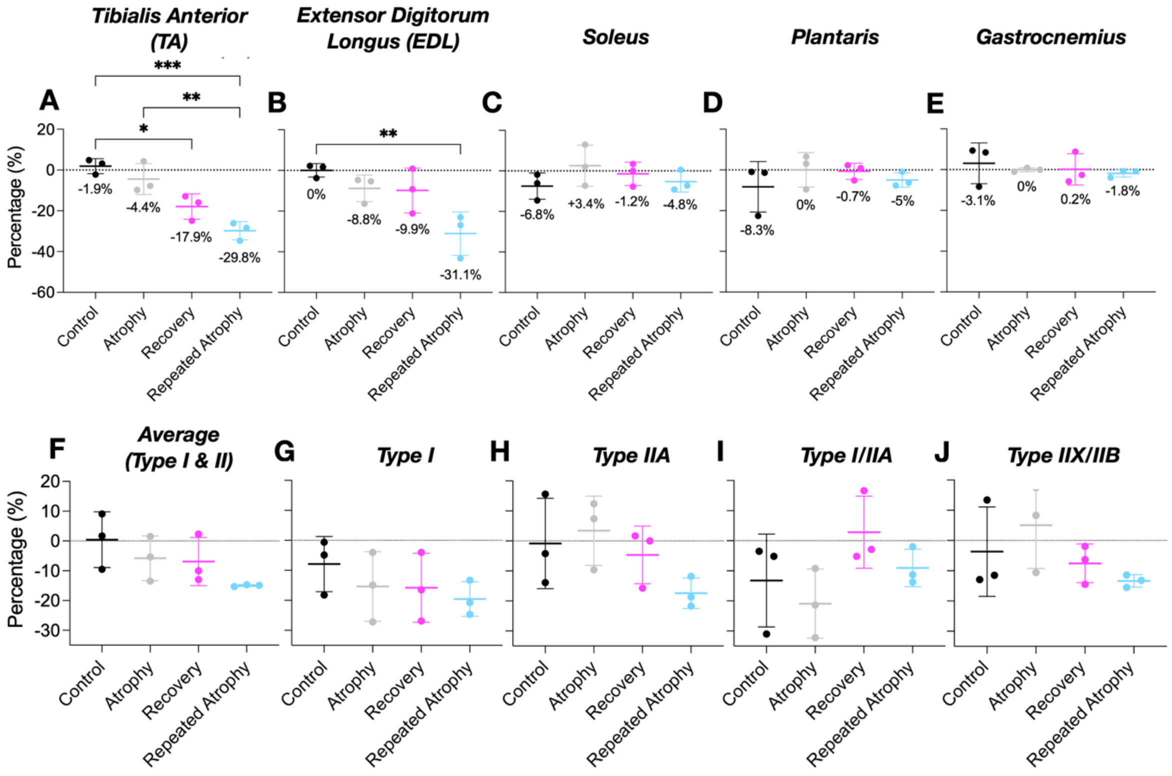


**Supplementary Figure 3**. Muscle weights of *TA* (**A**), *EDL* (**B**), *soleus* (**C**), *plantaris* (**D**) and *gastrocnemius* (**E**) muscles. Muscle weights presented as % change of the left surgical versus right non-surgical control limb. **p* ≤ 0.05, ***p* ≤ 0.01, ****p* ≤ 0.001. Relative fCSA (left/surgical % of right/non-surgical limb normalized to body mass) for each fibre type (**F**) - average type I & II, (**G**) - type I, (**H**) -type I IA, (**I**) -Type I/IIA and (**J**) - Type IIX/IIB. Control consists of pooled right contralateral control limbs across all conditions (*n* = 12). **p* ≤ 0.05.
