## Supplementary Figure 4 for "Repeated disuse atrophy imprints a molecular memory in skeletal muscle: transcriptional resilience in young adults and susceptibility in aged muscle"

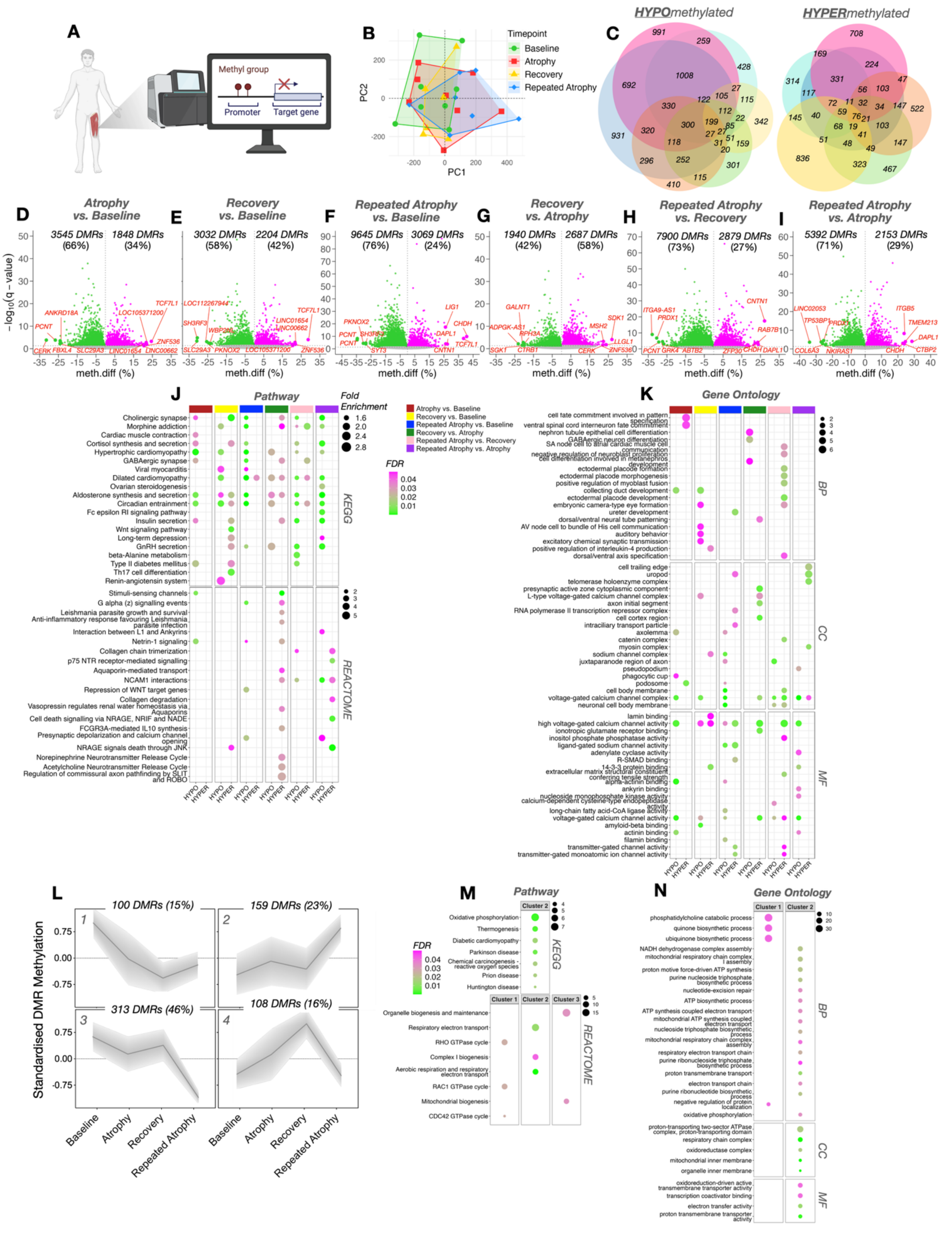

**Supplementary Figure 4.** (**A**) Schematic depicting the genome-wide DNA methylation (via RRBS) response to repeated disuse in adult human skeletal muscle (created with BioRender.com). (**B**) Principal component analysis (PCA) including clustering by time point. (**C**) Venn diagrams depicting the number of unique and common genes with associated HYPO and HYPER differential methylated regions (DMRs). (**D-I**) Volcano plots depicting the number of DMRs that were either HYPO- (green) or HYPER-methylated (magenta). Gene symbols within volcano plots represent the top 5 DMRs associated with those genes according to greatest methylation difference (meth.diff %) with FDR < 0.05. Over-representation analysis (ORA) of the top 20 enriched (**J**) pathway (KEGG, top 4 panels; REACTOME, bottom 4 panels) and (**K**) gene ontology (GO) terms for both HYPO- and HYPER-methylated DMRs across all comparisons ("Atrophy vs. Control", "Recovery vs. Control", "Repeated Atrophy vs. Control", "Recovery vs. Atrophy", "Repeated Atrophy vs. Recovery"). (**L**) Self-Organizing Maps (SOM) gene clustering analysis of 608 DMRs by gene symbol (that were significantly differentially methylated in at least one comparison) revealed most DMRs were either HYPER- (Cluster 2) or HYPO-methylated (Cluster 3) after atrophy, return to baseline levels following recovery with larger changes (in the same direction) after repeated atrophy. Pathway (**M**) and (**N**) gene ontology (GO) enrichment analysis of these SOM cluster profiles revealed most enriched terms were related to oxidative and energy metabolism and mitochondrial biogenesis / function. BP = biological process, CC = cellular component, MF = molecular function.
