## Supplementary Figure 5 for "Repeated disuse atrophy imprints a molecular memory in skeletal muscle: transcriptional resilience in young adults and susceptibility in aged muscle"

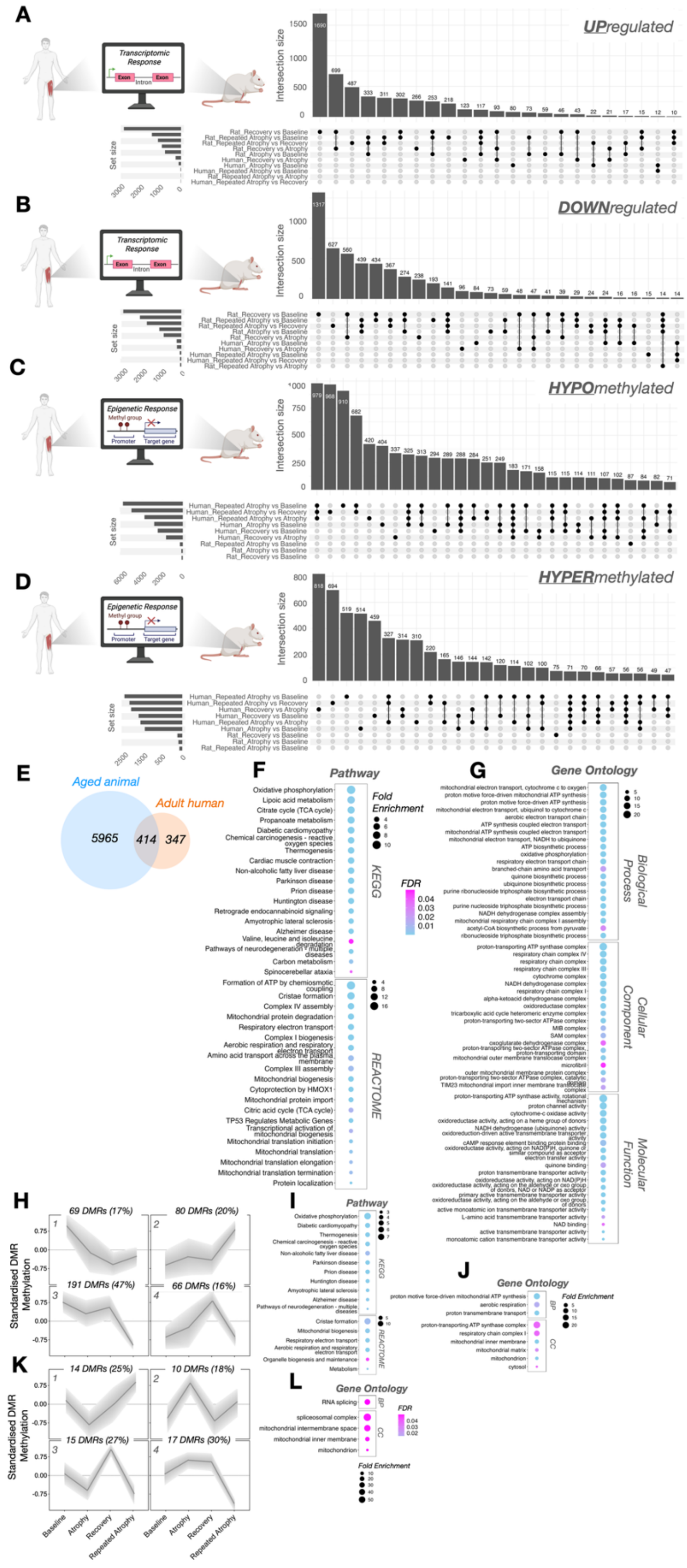


**Supplementary Figure 5.** Comparison of the transcriptomic (**A & B**) and epigenetic (**C & D**) responses to repeated disuse atrophy in young adult human and age rat skeletal muscle across all six pairwise comparisons. Identifying several common and unique UP- (**A**) and DOWN-regulated (**B**) DEGs as well as HYPO- (**C**) and HYPER-methylated (**D**) DMRs within each pairwise comparison. (**E**) Venn diagram analysis of all genes that were significantly differentially expressed in at least 1 of the pairwise comparisons for both young adult human (762 DEGs) and aged animal (6379 DEGs) identified 414 shared DEGs across both models of repeated disuse. Pathway (**F**) and GO (**G**) analyses demonstrated these genes were related to oxidative metabolism, energy metabolism and mitochondrial function. (**H**) SOM profile DNA methylation analysis identified 406 DMRs on 189 out of 414 DEGs in adult human muscle, with genes related to oxidative metabolism and mitochondrial function (**I & J**). (**K**) SOM profile DNA methylation analysis in aged animals identified 56 DMRs on 16 /DEGs with genes also related mitochondria as well as RNA splicing (**L**).
