## Supplementary Figure 6 for "Repeated disuse atrophy imprints a molecular memory in skeletal muscle: transcriptional resilience in young adults and susceptibility in aged muscle"

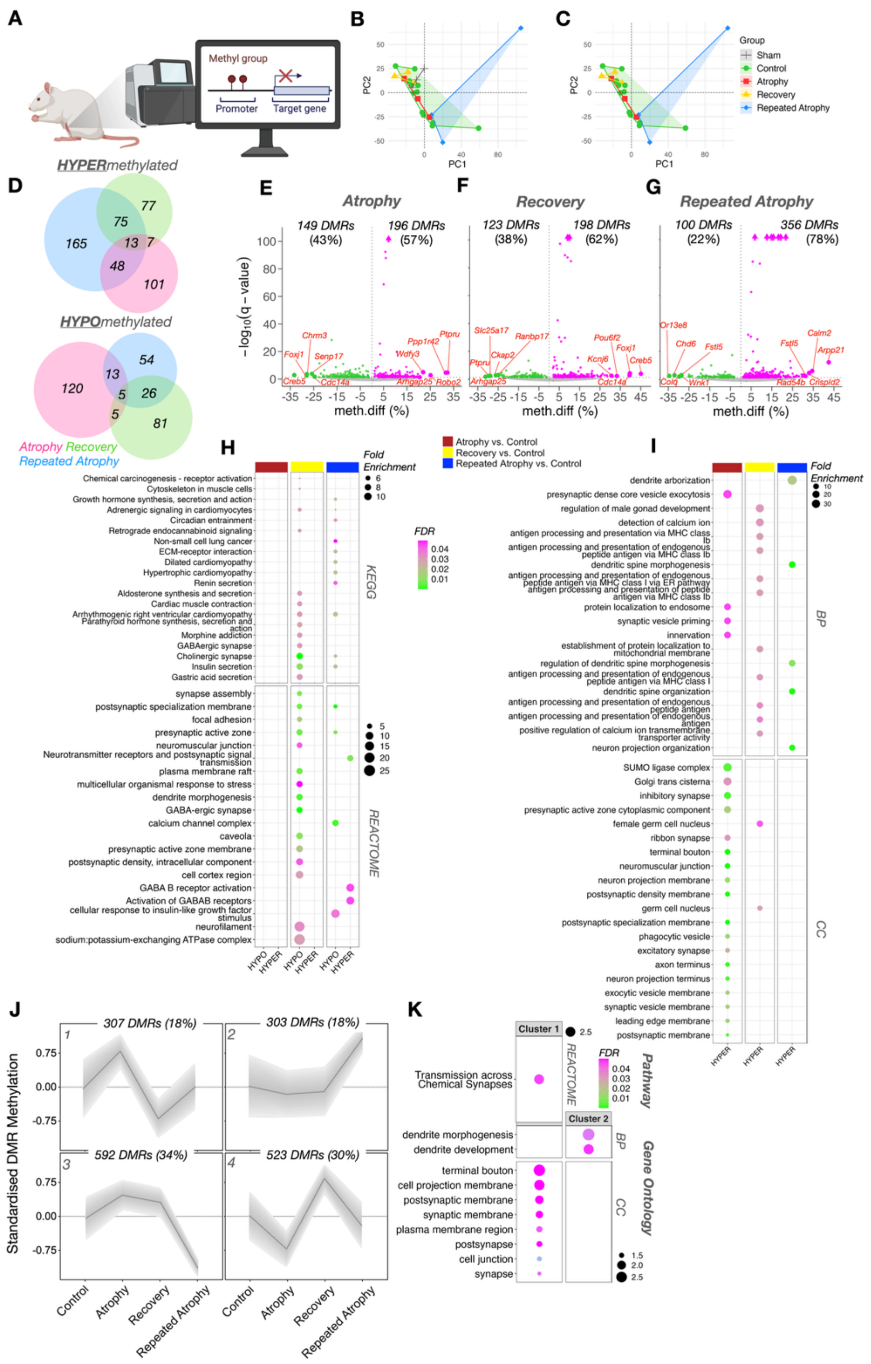


**Supplementary Figure 6.** (**A**) Schematic depicting the genome-wide DNA methylation (via Reduced Representation Seq) response to repeated disuse atrophy in aged rat skeletal muscle (created with BioRender.com). (**B & C**) Principal component analysis (PCA) with clustering by time point, including (**B**) and excluding (**C**) sham control samples. (**D**) Venn diagrams depicting the number of unique and common HYPO and HYPER differential methylated regions (DMRs) by gene symbol. (**E-G**) Volcano plots depicting the number of DMRs that were either HYPO- (green) or HYPER-methylated (magenta). Gene symbols within volcano plots represent the top 5 DMRs associated with those genes according to greatest methylation difference (meth.diff %) with FDR < 0.05. Over-representation analysis (ORA) of the top 20 enriched (**H**) pathway (KEGG, top 3 panels; REACTOME, bottom 3 panels) and (**I**) gene ontology (GO) terms for both HYPO- and HYPER-methylated DMRs across the main pairwise comparisons ("Atrophy vs. Control", "Recovery vs. Control", "Repeated Atrophy vs. Control"). (**J**) Self-Organizing Maps (SOM) analysis of 1725 DMRs by gene symbol identified gene clusters with similar temporal profiles across the time course of repeated disuse. Pathway (**J**) and (**K**) gene ontology (GO) analysis of these SOM cluster profiles revealed most enriched terms related to the neuromuscular junction development, structure and action potential. BP = biological process, CC = cellular component, MF = molecular function.
